# The loss of a supergene in obligately polygynous *Formica* wood ant species

**DOI:** 10.1101/2024.09.20.613865

**Authors:** Hanna Sigeman, Ina Satokangas, Matthieu De Lamarre, Riddhi Deshmukh, Pierre Nouhaud, Heikki Helanterä, Michel Chapuisat, Jonna Kulmuni, Lumi Viljakainen

## Abstract

Some of the most striking examples of phenotypic variation within species are controlled by supergenes. However, most research on supergenes has focused on their emergence and long-term maintenance, leaving the later stages of their life cycle largely unexplored. Specifically, what happens to a derived supergene haplotype when the trait it controls reaches fixation? Here we answer this question using the ancient supergene system of *Formica* ants, where (monogynous) single-queen colonies carry only the ancestral haplotype M while the derived haplotype P is exclusive to (polygynous) colonies with multiple queens. Through comparative genomics of all seven European wood ant species, we found that the P haplotype was present in only one of the three obligate polygynous species (*F. polyctena*). In the two others (*F. aquilonia* and *F. paralugubris*) the P haplotype was completely missing except for duplicated P-specific paralogs of two genes, *Zasp52* and *TTLL2*, with *Zasp52* being directly involved in wing muscle development. We hypothesize that these genes play a direct role in polygyny and contribute to differences in body size and/or dispersal behaviour between monogynous and polygynous queens. A complete lack of P/P genotypes among the 136 workers suggest strong selection against such genotypes. While our analyses did not reveal evidence of increased mutation load on the P, it is possible that this skew in genotype distributions is driven by a few loci with strong fitness effects. We propose that selection to escape P-associated fitness costs underlies the loss of this haplotype in obligate polygynous wood ants.

## Introduction

Supergenes underlie some of the most dramatic within-species polymorphisms in nature, such as color morphs in mimetic butterflies (Clarke & Sheppard, 1960; Joron et al., 2006; Kimura, 1956), male plumage morphs in birds (Küpper et al., 2016; Lamichhaney et al., 2016), and colony social organization in ants (Brelsford et al., 2020; Purcell et al., 2014; Wang et al., 2013). They often form through structural rearrangements which block recombination with the ancestral haplotype and can be selectively beneficial and rise in frequency if they capture multiple co-adapted alleles (Charlesworth, 2016; John R. G. Turner, 1967; Kimura, 1956; Thompson & Jiggins, 2014). Novel, or derived, supergene haplotypes are likely to have either higher or lower fitness than the ancestral haplotype, leading to their fixation or elimination (Schwander et al., 2014). However, supergenes can also exist as stable polymorphisms maintained by balancing selection for millions of years and across numerous speciation events (Brelsford et al., 2020; Purcell et al., 2014; Wang et al., 2013). Empirical work on supergenes is predominantly aimed at understanding the selection regimes and molecular dynamics underlying their emergence and maintenance in species with phenotypic variation, leaving the later stages of their evolutionary trajectory largely unexplored. Specifically, what happens to a supergene system in species where the phenotypic polymorphism it controls have disappeared?

In species where the derived trait has become fixed one might intuitively also expect (1) fixation of the derived haplotype, as seems to have happened in two species of mimetic swallowtail butterflies (Zhang et al., 2017). However, this is not possible in all supergene systems since derived haplotypes often become degenerated due to inefficient selection against deleterious mutations, leading to strong negative fitness effects or even homozygous lethality (Jay et al., 2021; Llaurens et al., 2017; Schwander et al., 2014). Alternative outcomes are then to either (2) maintain the supergene polymorphism despite lacking the phenotypic variation, as shown possible from population genetic models (Tafreshi et al., 2022), or to (3) retain only a subset of genes from the derived haplotype needed to uphold the derived trait, while otherwise reverting to fixation of the ancestral haplotype. This last outcome would allow purging of deleterious mutations accumulated on the derived haplotype. Here, we test between these molecular outcomes using the ancient supergene system of *Formica* ants (∼20-40 my; Brelsford et al., 2020; Purcell et al., 2021).

Many *Formica* ants display intra-species variation in reproductive queen number, with some colonies being headed by a single queen (monogyny) while others contain multiple, often non-related, queens (polygyny) (Chapuisat et al., 2004; DeHeer & Herbers, 2004; Goropashnaya et al., 2001; Rosset & Chapuisat, 2007; Seppä et al., 2004; Sundström, 1993). In such socially polymorphic *Formica* species, queen number is associated with a suite of behavioral, phenotypic, and life-history traits, collectively referred to as the ‘polygyny syndrome’ (sensu Keller, 1993). These include differences in colony size, queen and worker body size, and dispersal behaviour (Boomsma et al., 2014; Keller, 1991; Rosset & Chapuisat, 2007). The polygyny syndrome is genetically controlled by a ∼10 Mb supergene on chromosome 3, which formed through multiple inversion events in a shared ancestor (Purcell et al. 2014; Brelsford et al., 2020). In monogynous *Formica* colonies, all individuals exclusively carry the ancestral haplotype M (diploid workers and queens are M/M; haploid males are M), while the derived P haplotype is always present within polygynous colonies though not necessarily in every individual (Brelsford et al., 2020, 2020; Lagunas-Robles et al., 2021; McGuire et al., 2022; Pierce et al., 2022; Purcell et al., 2014). The P haplotype, while carrying genes adapted to polygyny, is also associated with recessive fitness costs. This is evidenced by lower survival rates of P/P females in *F. selysi* compared to other genotypes (Blacher et al., 2023), and by a scarcity of P/P workers outside the clade including *F. selysi*, *F. cinerea*, and *F. lemani* (Avril et al. 2019, Purcell et al. 2014, Brelsford et al. 2020, Purcell et al. 2020, Pierce et al., 2022).

In contrast to previous studies, we are investigating the evolutionary fate of the supergene in species where multi-queen colonies have become a fixed trait (obligate polygyny). We targeted all seven European species of the *Formica rufa* (“wood ants” or *Formica s.str* group) complex, in which three are obligate polygynous (*F. aquilonia*, *F. paralugubris* and *F. polyctena*) and four are socially polymorphic (*F. lugubris*, *F. pratensis*, *F. rufa*, and *F. truncorum*). Using comparative population genomic analyses, we show that the molecular outcome (see above; 1-3) of the supergene system varies between the obligate polygynous species. We also show that species in which a derived supergene-associated trait has become fixed are powerful study systems for revealing genes involved in these traits.

## Results and discussion

### Obligate polygynous Formica species are lacking the polygyny-associated haplotype P

We determined supergene genotype distributions across the seven *Formica* species using whole-genome sequencing data of 139 individuals (Table S1) with an average sequencing depth of 16.2x (Table S2). Diploid individuals (workers and queens) with M/P supergene genotypes are expected to display high heterozygosity levels across the supergene region on chromosome 3. Of the 136 workers in our dataset, 16 were deemed as highly heterozygous based on inbreeding coefficients (F_IS_ scores < −0.5; Figure 1a; Table S3). The heterozygous SNPs in these workers were concentrated within a 10.7 Mb central region of chromosome 3 positioned between 1.9 and 11.6 Mb (Figure 1b; Figure S1). This region is highly syntenic to the supergene region in other *Formica* species (Brelsford et al., 2020; Lagunas-Robles et al., 2021; Purcell et al., 2021), confirming their status as M/P heterozygotes (n = 5 *F. lugubris*, n = 8 *F. pratensis*, and n = 3 *F. polyctena*).

**Figure 1.**
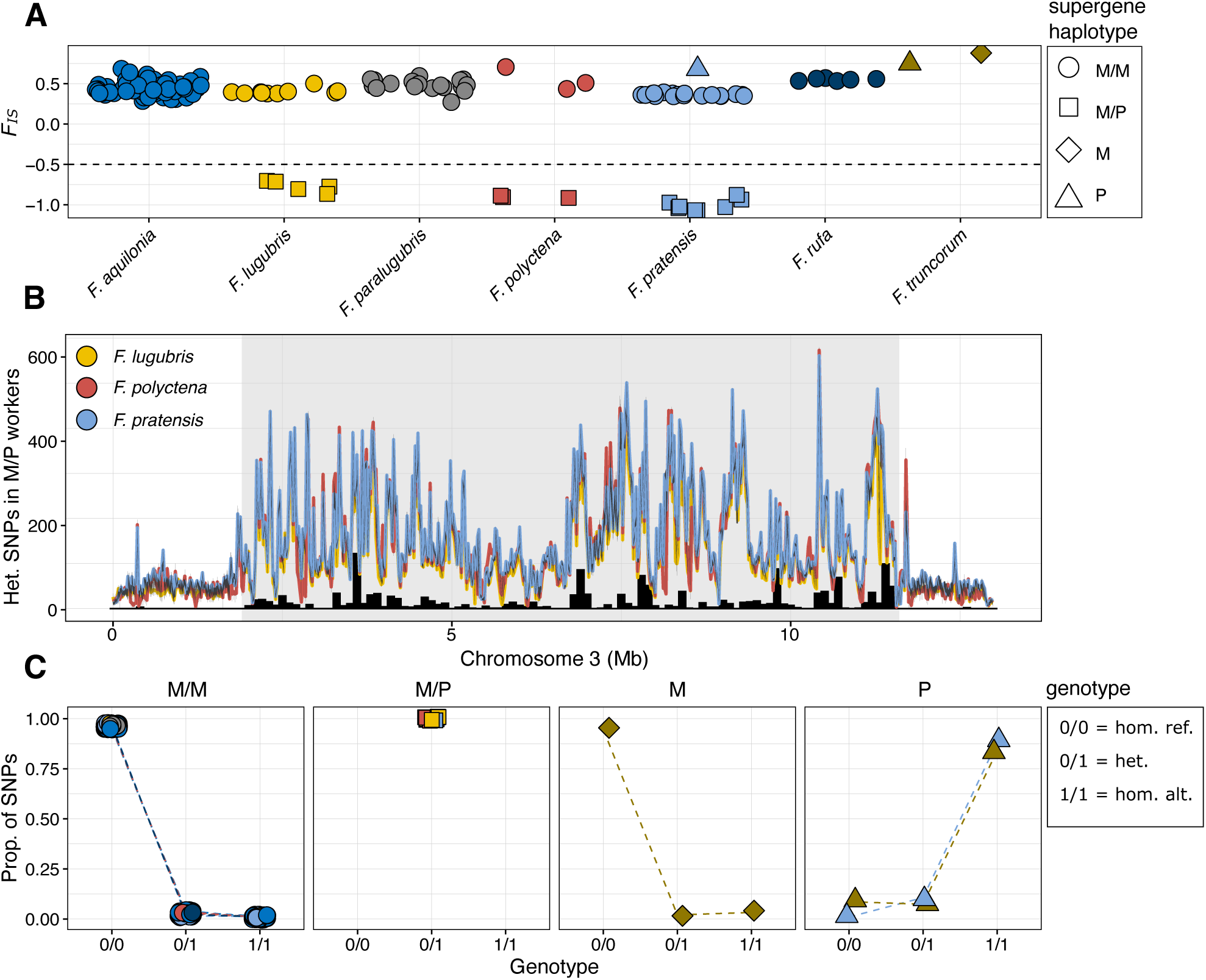
(**A**) Inbreeding coefficients (F_IS_ scores) for all samples (n = 136 diploid workers, n = 3 haploid males). The shape of the data points shows the supergene haplotype, as revealed by the results in **A**, **B** and **C**. The 16 workers with inbreeding coefficients <-0.5 (below the dashed line) were classified as M/P heterozygotes. (**B**) The lines show mean number of heterozygous SNPs (20kb windows) per species, for the 16 M/P workers (n = 5 F. lugubris, n = 8 F. pratensis, and n = 3 F. polyctena). The supergene region (1.9−11.6 Mb) is highlighted with a gray background colour. The black histogram shows the number of sites that were heterozygous in all 16 workers (“trans-species haplotype-specific SNPs”; n = 21,790). (**C**) Relative proportions of genotype counts (0/0 = reference homozygous, 0/1 = heterozygous, 1/1 = alternative homozygous) across the 21,790 trans-species haplotype-specific SNPs for each individual (i.e., the total value for each individual amount to 1.00). A dashed line connects the data points from the same individual. The shape and colour of the data points are consistent with **A**.

The genotypes of the remaining workers (n = 120; F_IS_ scores > -0.5; Figure 1a; Table S3) were determined based on their allele distribution at the 21,790 sites on chromosome 3 which were heterozygous across all 16 workers with M/P genotypes (referred to as “trans-species haplotype-specific SNPs”; Figure 1b,c), and by the inclusion of three (haploid) males from a previous study with known supergene status (Table S1; Brelsford et al., 2020). All non-M/P workers were almost exclusively homozygous for the reference allele (Figure 1c; Table S4), similarly to the *F. truncorum* male with a known M genotype (Figure 1c; Table S4), and therefore classified as M/M homozygotes. These results also show that the reference genome, which was constructed from a single male from a *Formica aquilonia x polyctena* hybrid population in southern Finland (Nouhaud et al., 2022), contains the M haplotype. This conclusion is further supported by the *F. pratensis* and *F. truncorum* males with known P genotypes, which both had the alternative allele across almost all sites (Figure 1c; Figure S1-2; Table S4).

Counter-intuitively, the polygyny-associated P haplotype was completely lacking from two of the three obligate polygynous species (*F. aquilonia* and *F. paralugubris*; Figure 2; Table S5). This is a surprising finding as the species share multiple polygyny syndrome characteristics assumed to be linked to the P haplotype, such as high queen number per colony, large colony sizes, and colony formation mainly through budding (Seifert, 2018). While we lack direct evidence on colony social structure for the workers in this study, we can safely assume that these M/M workers (Figure 2) originate from polygynous colonies. *F. aquilonia* is known to be polygynous across all studied European populations (Pamilo et al., 1992), and the workers in this study were collected from numerous colonies over a wide geographical range (n = 64; Finland, Russia, Scotland, and Switzerland; Table S1). The *F. paralugubris* workers were collected from a single population in Switzerland (n = 19; Table S1), where their highly polygynous social organization has been well characterized (Chapuisat et al., 1997; Pamilo et al., 1992). The absence of P in these species shows that fixation of the polygyny syndrome did not occur through fixation of the polygyny-associated haplotype (thus ruling out “molecular outcome 1”; see Introduction). It also shows that the ancient and stable *Formica* supergene polymorphism has disappeared from these obligate polygynous species (i.e., ruling out also “molecular outcome 2”).

**Figure 2.**
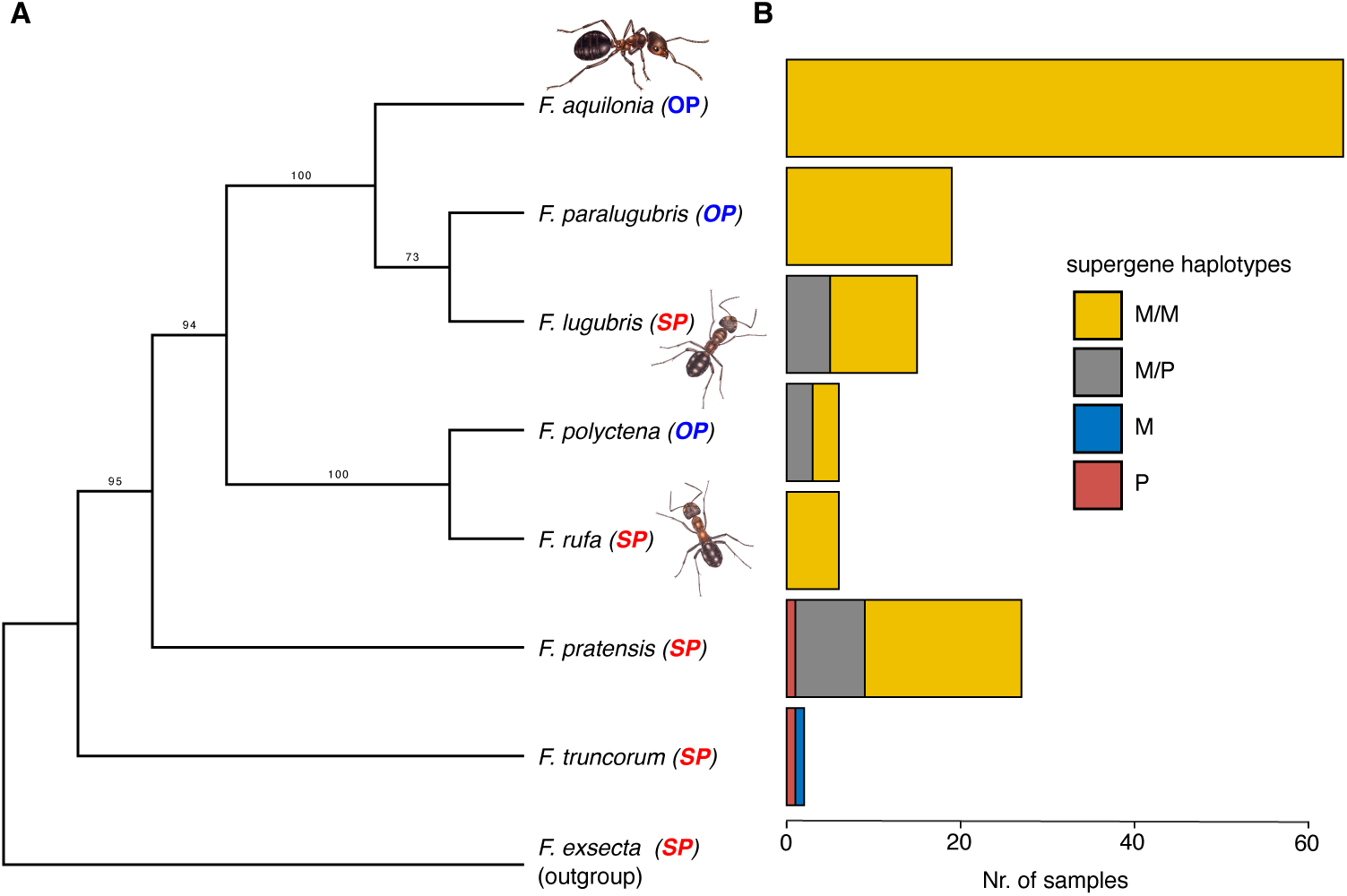
(**A**) Phylogeny of the seven studied species, including an outgroup (F. exsecta, also carrying the supergene), based on SNPs from chromosome 3 outside the supergene region (see Methods). Bootstrap values are displayed as branch labels. OP: obligate polygynous, SP: socially polymorphic. (**B**) The distribution of supergene genotypes for each species (Table S5). Ant drawings by Lizzie Harper.

In contrast, both supergene haplotypes M and P were found within individuals of the obligate polygynous species *F. polyctena* (n = 3/6; Figure 2b; Table S5; i.e., “molecular outcome 2”), and within three of the four socially polymorphic species (n = 5/15 *F. lugubris*, n = 7/25, *F. pratensis*, and n = 1/2 *F. truncorum*; Figure 2b; Table S5). It is highly likely that the individuals from socially polymorphic species carrying P haplotypes originate from polygynous colonies. This is because all *Formica* workers from monogynous colonies studied so far have been exclusively M/M (Brelsford et al., 2020; McGuire et al., 2022; Pierce et al., 2022; Purcell et al., 2014). Note that the polygynous social colony structure is confirmed for the two males with P haplotypes (Figure 2b; Table S5; Brelsford et al., 2020). Based on the supergene haplotype distributions in other *Formica* species, the colony social structure associated with the M/M genotypes can not be inferred with certainty (Figure 2). In the outgroup *Formica* species *F. selysi* and *F. exsecta*, M/M workers are exclusively found in monogynous colonies (Brelsford et al., 2020). However, M/M workers are present in both monogynous and polygynous colonies of *F. neoclara* (McGuire et al., 2022) and *F. francoeuri* (Pierce et al., 2022). Lastly, while it is possible that the absence of P haplotypes in *F. rufa* is representative of the species (Figure 2), we find this result more likely to be an effect of the low sample size and the mainly monogynous colony social structure within the sampling range (n = 6 workers, each collected from independent populations in Finland; Table S1).

### Two candidate genes for the maintenance of polygyny

So how is the polygyny syndrome maintained in *F. aquilonia* and *F. paralugubris* despite lacking the P haplotype? One explanation would be if a subset of P-specific gene copies necessary for upholding polygyny remains on chromosome 3 in these species (“molecular outcome 3”), which could be achieved through double crossing over or gene conversion between the supergene haplotypes. This hypothesis is plausible in the *Formica* system as rare recombination events are known to have homogenized varying parts of the M and P haplotypes within different socially polymorphic species (Brelsford et al., 2020). Previous studies have suggested that genes containing trans-species P-specific SNPs (i.e., which have not experienced recombination between haplotypes) are especially likely to play a functional role in maintaining the polygyny syndrome in *Formica* ants (Brelsford et al., 2020; Purcell et al., 2021). We therefore searched for genes with elevated numbers of such P-specific alleles (defined as alternative homozygous or heterozygous SNPs) in obligate polygynous M/M workers compared to socially polymorphic M/M workers. This is because we do not expect the M haplotype in socially polymorphic species to contain genes coding for polygynous traits.

We found P-specific copies of only two genes, *Zasp52* (“Z band alternatively spliced PDZ-motif protein 52”) and *TTLL2* (“Tubulin Tyrosine Ligase Like 2”), in M/M workers of obligate polygynous species but not in M/M workers of socially polymorphic species (Figure 3). Both genes, which are adjacently positioned (9.80−9.85 Mb; Figure 3c), had significantly more P-specific SNPs in the obligate polygynous species (*Zasp52*: n = 28 SNPs, two-tailed t-test: *p*=0.024; *TTLL2*: n = 8 SNPs; two-tailed t-test: *p*=0.029; Figure 3a,b; Table S6,7). After Bonferroni correction for the number of tests (n = 292), all p-values obtained the value of 1. The SNPs within these genes were almost completely heterozygous in all M/M workers of *F. aquilonia* (n = 64/64), most M/M workers of *F. paralugubris* (n = 15/19) and at lower frequencies in M/M workers of *F. polyctena* (n = 1/3), similarly to all 16 M/P workers of the socially polymorphic species (Figure 3d). In contrast, the M/M workers of the socially polymorphic species were mainly homozygous for the reference (i.e., ancestral) allele. Note, however, that the M/M workers of the socially polymorphic species were mainly alternative homozygous at 6 of the 36 trans-species haplotype-specific SNPs, suggesting that the M haplotype in the *F. aquilonia x polyctena reference* genome contain some P-specific alleles (Figure 3d).

**Figure 3.**
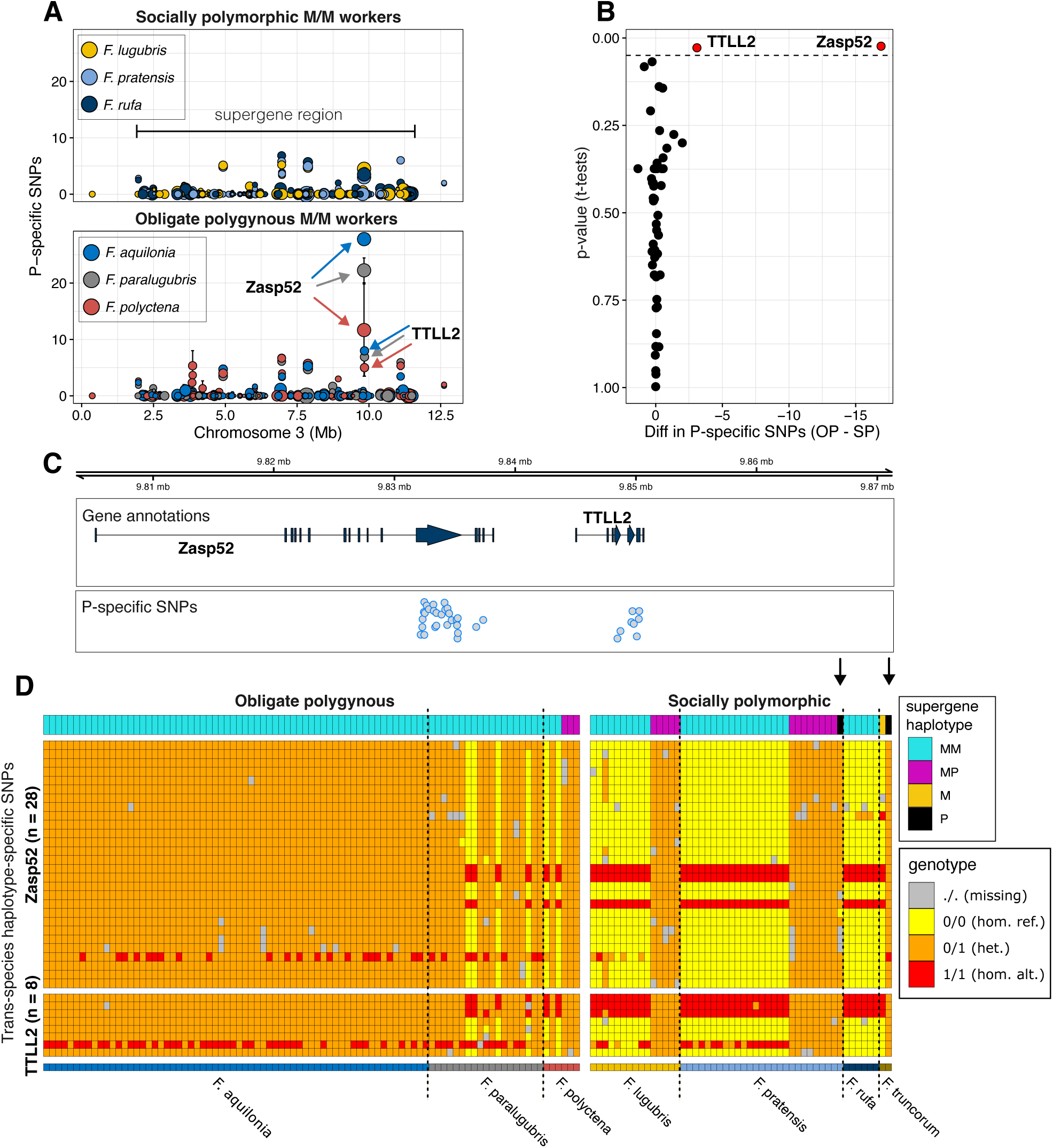
(**A**) Mean number of P-specific SNPs per gene and species (error bars represent ±SE), plotted separately for socially polymorphic (top panel) and obligate polygynous (bottom panel) species. (**B**) X-axis: Differences in mean number of P-specific SNPs for socially polymorphic (SP) and obligate polygynous (OP) species. Y-axis: P-values from t-tests between the number of P-specific SNPs in socially polymorphic and obligate polygynous species. The dashed line marks the 0.05 significance level, and genes with significant differences are labelled and coloured in red. (**C**) Gene annotations and genome positions of the two genes, Zasp52 and TTLL2, with significantly more P-specific SNPs in the obligate polygynous species. Data points for the P-specific SNPs are shown below, jittered along the y-axis for increased visibility. (**D**) Heatmap showing from the top: (i) supergene haplotype, (ii) genotypes at the trans-species haplotype-specific SNPs of the genes Zasp52 and TTLL2, and (iii) species. The dashed vertical lines also separate the species. The black arrows highlight the two P males (n = 1 F. pratensis, n = 1 F. truncorum), which both appear completely heterozygous despite being haploid, suggesting P-specific duplications of these genes.

We propose that the P-specific copies of *Zasp52* and *TTLL2* on the M haplotype may be responsible for upholding the polygyny syndrome in *F. aquilonia* and *F. paralugubris*. *Zasp52* is a well-studied central gene for muscle tissue development in insects, specifically involved in the formation and maintenance of Z-discs which act as boundaries between muscle sarcomeres (Chechenova et al., 2013; Katzemich et al., 2013; K. A. Liao et al., 2016). It is reasonable to imagine that such a gene could be involved in the polygyny syndrome, as studies of socially polymorphic species have shown that queens from polygynous colonies are generally smaller and disperse over shorter distances than those from monogynous colonies. Interestingly, flight muscles of polygynous *F. truncorum* queens have fewer mitochondria and sarcomeres, and may deteriorate faster (shown by heavily distorted Z-discs in dealate queens) than those of monogynous queens (Johnson et al., 2005). Note, however, that in other (non-wood ant) *Formica* species, polygyny is significantly associated with higher muscle mass in males but not in queens (*F. exsecta*), or not associated with colony social structure in either sex (*F. pressilabris*; (Hakala, 2020)). *TTLL2* has not been studied in insects but is in humans almost exclusively expressed in sperm (Fagerberg et al., 2014).

Combined evidence suggests that a duplication event created paralogs of *Zasp52* and *TTLL2* on the P haplotype in a shared ancestor of all wood ants, and that these P-specific gene copies later translocated to the M haplotype in the obligate polygynous species. Firstly, the two male samples with P haplotypes (*F. pratensis* and *F. truncorum*) are completely heterozygous for all trans-species haplotype-specific SNPs within these two genes (Figure 3d). This points towards a P-specific duplication of these genes, as (haploid) males cannot be truly heterozygous (see e.g., (Purcell et al., 2021)). This is further supported by heightened genome coverage values within this region in individuals with high heterozygosity (Figure S3, S4). Secondly, the high number of trans-species haplotype-specific SNPs shared by the M/P workers as well as the P males suggest that the duplication occurred in a shared ancestor of all wood ants (Figure 3d). And thirdly, *de novo* genome assemblies based on *F. aquilonia* and *F. polyctena* males with M haplotypes (see Methods; Table S8, S9) revealed two paralogs each of *Zasp52* and *TTLL2*, on separate contigs (Table S10), indicating translocation of the P-specific copies to the M haplotype background in obligate polygynous wood ants (Figure 4). In both genome assemblies, one gene copy was highly similar to the gene copy in the *F. aquilonia x polyctena* reference genome annotation, and to the closest non-wood ant relative *F. exsecta*. These were classified as the original gene copies (Figure 4; tip labels in blue). The other gene copy in the *F. aquilonia* and *F. polyctena* genomes, which were highly similar to each other, shared nucleotide sequence similarity to their paralogs of 70% (*Zasp52*) and 92% (*TTLL2*). These were classified as the duplicated, P-specific, paralogous gene copies (Figure 4; in red). The functionality of both gene copies was confirmed by gene expression using RNA data from 20 pooled *F. aquilonia* workers (see Methods; Figure S5).

**Figure 4.**
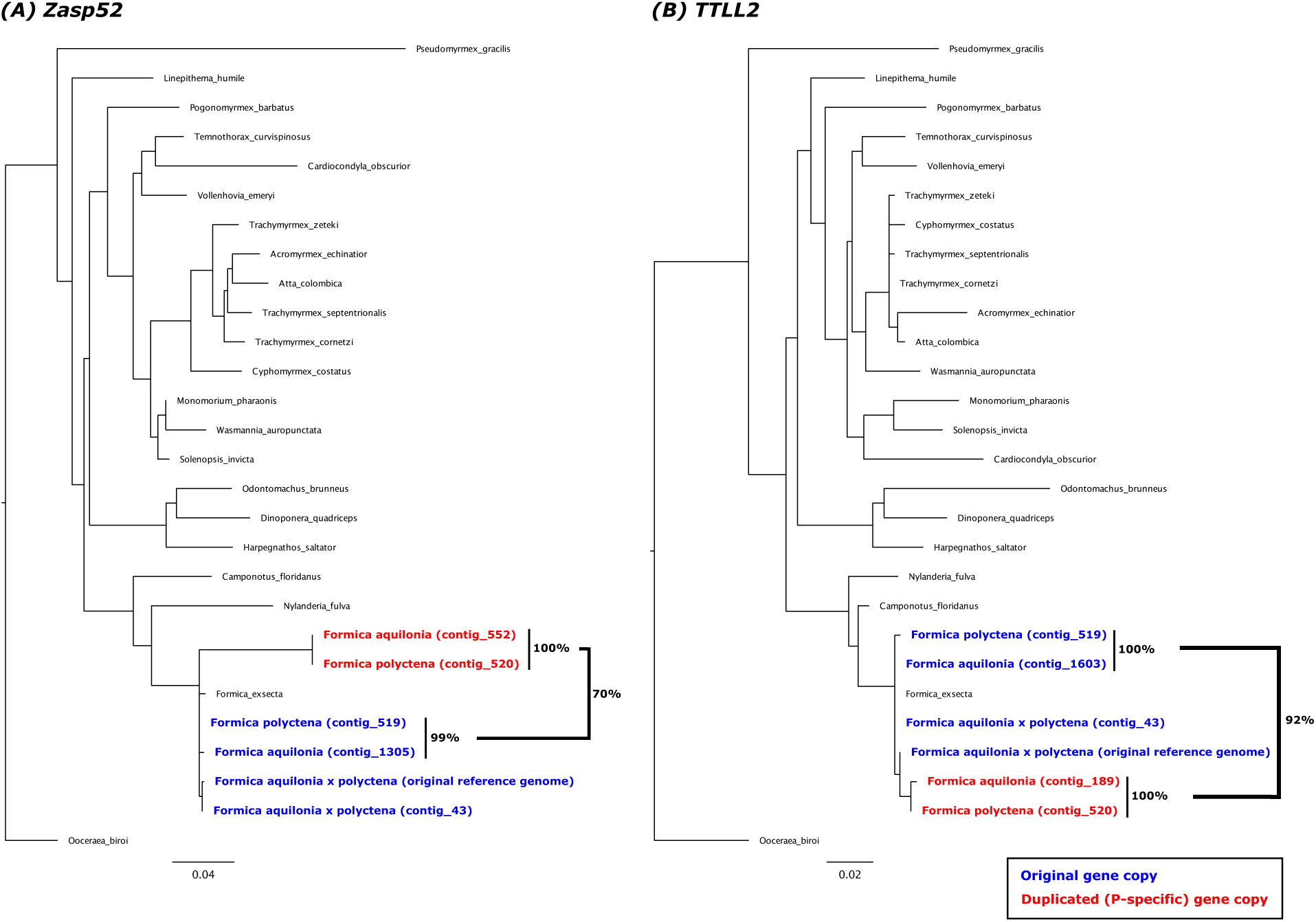
Amino acid gene trees of (**A**) Zasp52 and (**B**) TTLL2. Gene sequences are from the (1) Formica aquilonia x polyctena reference genome, de novo assemblies of PacBio data from males of (2) F. aquilonia, (3) F. polyctena, and the (3) Formica aquilonia x polyctena reference genome individual, as well as (4) 22 outgroup ant species. The original (blue) and duplicated (P-specific; red) gene copies from the Formica rufa complex species are highlighted, as well as the sequence similarities (%) between some of the gene copies.

Since the species carrying both gene copies of *Zasp52* and *TTLL2* on the M haplotype (*F. aquilonia*, *F. paralugubris*, and *F. polyctena*) are not monophyletic (Figure 2), we find the most parsimonious interpretation of these patterns to be that a recombination event between the M and P haplotype in one of the three obligate polygynous species (most likely *F. aquilonia* or *F. paralugubris*, see below) created a “minimal P” haplotype, specifically a M haplotype containing the P-specific copies of *Zasp52* and *TLLL2*, followed by adaptive introgression of this new haplotype to the other species (see e.g., (Stolle et al., 2022)). Alternatively, the translocation could have originated in a shared ancestor of the closely related species *F. aquilonia* and *F. paralugubris* (Figure 2), in which it is found in high frequencies (*F. aquilonia*: n = 64/64; *F. paralugubris*: n = 15/19) and more recently introgressed to *F. polyctena* in which the translocation occurs at lower frequencies (n = 1/3 M/M workers). A third and less likely scenario is if the translocation occurred more basally in the group, e.g., after the split between *F. pratensis* and the other wood ant species, followed by repeated losses across the socially polymorphic species. The P-specific duplication was not found among six individuals belonging to five outgroup *Formica* species carrying P haplotypes (n = 1 P/P *F. exsecta*, n = 1 P/P *F. cinerea*, n = 2 P/P *F. lemani*, n = 1 M/P *F. obscuripes*, n = 1 P *F. selysi*; Table S1; (Purcell et al., 2021), as there were no heterozygous SNPs within these genes.

### The Knockout gene does not maintain polygyny in F. aquilonia and F. paralugubris

Previous research on the *Formica* supergene system have revealed other genes hypothesized to play a role in the polygyny syndrome. The main candidate is the gene *Knockout* (Table S6), where multiple trans-species haplotype-specific SNPs are conserved throughout large parts of the *Formica* phylogeny (Purcell et al., 2021). In wood ants, 19 SNPs within the coding-region of *Knockout* were similarly trans-species haplotype-specific, but none were P-specific in the two obligate polygynous species lacking P haplotypes (*F. aquilonia* and *F. paralugubris*; Figure S6). It is therefore unlikely that *Knockout* is crucial for upholding the polygyny syndrome in these species. We also identified single-copy orthologs to four of the six additional candidate genes described in Purcell et al. (Purcell et al., 2021), all harboring large numbers of trans-species haplotype-specific SNPs between a wide range of *Formica* species (*AmGR10*, *FMFFaR*, *Single-minded*, and *ZPF148*; Table S6). Among the three genes that displayed trans-species haplotype-specific SNPs also within the wood ants (*AmGR10*: n = 10; *Single-minded*: n = 11; *ZPF148*: n = 5), no SNPs were P-specific in *F. aquilonia* or *F. paralugubris* (Figure S6). Note that *Zasp52* and *TTLL2* have not been proposed as candidate genes underlying the polygyny syndrome in other *Formica* species, suggesting that the genetic architecture of polygyny may vary across the group.

### P/P lethality and mutation load as drivers behind the loss of the P haplotype

Three alternative molecular outcomes for a derived supergene haplotype in species where the trait controlled by this haplotype has reached fixation were previously outlined: (1) fixation of the derived P haplotype, (2) maintenance of the supergene polymorphism, or (3) retention only of a subset of P-specific genes needed to uphold the derived trait. We found that these molecular outcomes vary between the obligate polygynous species in this study. In *F. polyctena*, the supergene polymorphism remains (molecular outcome 2), while the derived supergene haplotype P has become limited to only a few genes in *F. aquilonia* and F*. paralugubris* (molecular outcome 3). Similarly to previous studies across many *Formica* ants (Avril et al. 2019, Purcell et al. 2014, Brelsford et al. 2020, Purcell et al. 2020, Pierce et al., 2022), we did not find any P/P homozygotes among the 136 workers in our dataset. This suggests that P/P homozygotes are either unviable or subject to strong negative selection (Blacher et al., 2023; Purcell et al., 2021). Based on these observations, it is likely that selection against P/P genotypes has prevented fixation of the P haplotype within obligate polygynous *Formica* wood ants (molecular outcome 1). Strong selection against P/P genotypes would also affect the fitness of M/P queens mating with a P male. Such effects were demonstrated in *F. selysi*, one of the few Formica species where P/P individuals have been found, where M/P and P/P queens laid smaller broods and had a smaller proportion of offspring reaching adulthood compared to M/M queens (Blacher et al., 2023). Interestingly, the study further showed that fewer M/P and P/P queens were fertile compared to M/M queens, and that M/P and P/P workers had a lower survival probability compared to M/M workers (Blacher et al., 2023). This suggests a mutation load for the P haplotype also when occurring in heterozygous form and may contribute to why a recombinant supergene haplotype in wood ants (molecular outcome 3) containing only a subset of genes from the P haplotype could have reached high frequencies (in *F. paralugubris*) and even fixation (in *F. aquilonia*).

To estimate whether the wood ant P haplotype carries a mutation load, we predicted the functional effect of all intragenic SNPs within the supergene region of chromosome 3 (categories “high”, “moderate”, and “low”; see Figure 5 legend for descriptions of each category). The M/P workers had a higher number of SNPs compared to M/M workers across all three impact-categories; high (Figure 5a; Wilcoxon: W = 0; p < 0.001), moderate (Figure 5b; Wilcoxon: W = 0; p < 0.001;), and low (Figure 5c; Wilcoxon: W = 0; p < 0.001). This is to be expected as we are using a reference genome with an M genotype. However, the proportion of high- and moderate-impact mutations were not higher in the M/P workers than in the M/M workers (Figure 5c-d), as would have been expected from a P haplotype carrying a high mutation load. Contrary to expectations, the M/M workers had a significantly higher proportion of high-impact mutations compared to the M/P workers (Figure 5d; Wilcoxon: W = 1435, p = 0.001), while there was no difference between the proportions of moderate-(Figure 5e; Wilcoxon: W = 938, p-value = 0.8845) or low-impact mutations (Figure 5f; Wilcoxon: W = 888, p-value = 0.6291). It is possible that recombination or gene conversion events between the M and P haplotypes, as observed within *Formica* ants (Brelsford et al., 2020), has repeatedly purged deleterious mutations from the P. This would be similar to how rare recombination events between the sex chromosomes in European tree frogs have purged deleterious mutation from the Y chromosome (Guerrero et al., 2012). While we do observe numerically higher numbers of SNPs with predicted high impacts on gene function (i.e., deleterious mutations) among the M/P workers than the M/M workers (Figure 5a), it is difficult to establish how many of these excess mutations are false positives created by the large evolutionary distance between the P and the gene models predicted based on the M haplotype in the reference genome. Future studies using a P haplotype reference genome will be able to determine the effect of these predicted high impact mutations on gene functionality. The weak signature of genetic load on the P suggest that the lack of P/P individuals in Formica wood ants may be the result of a few mutations with high deleterious effects rather than a general accumulation of deleterious mutations across the supergene.

**Figure 5.**
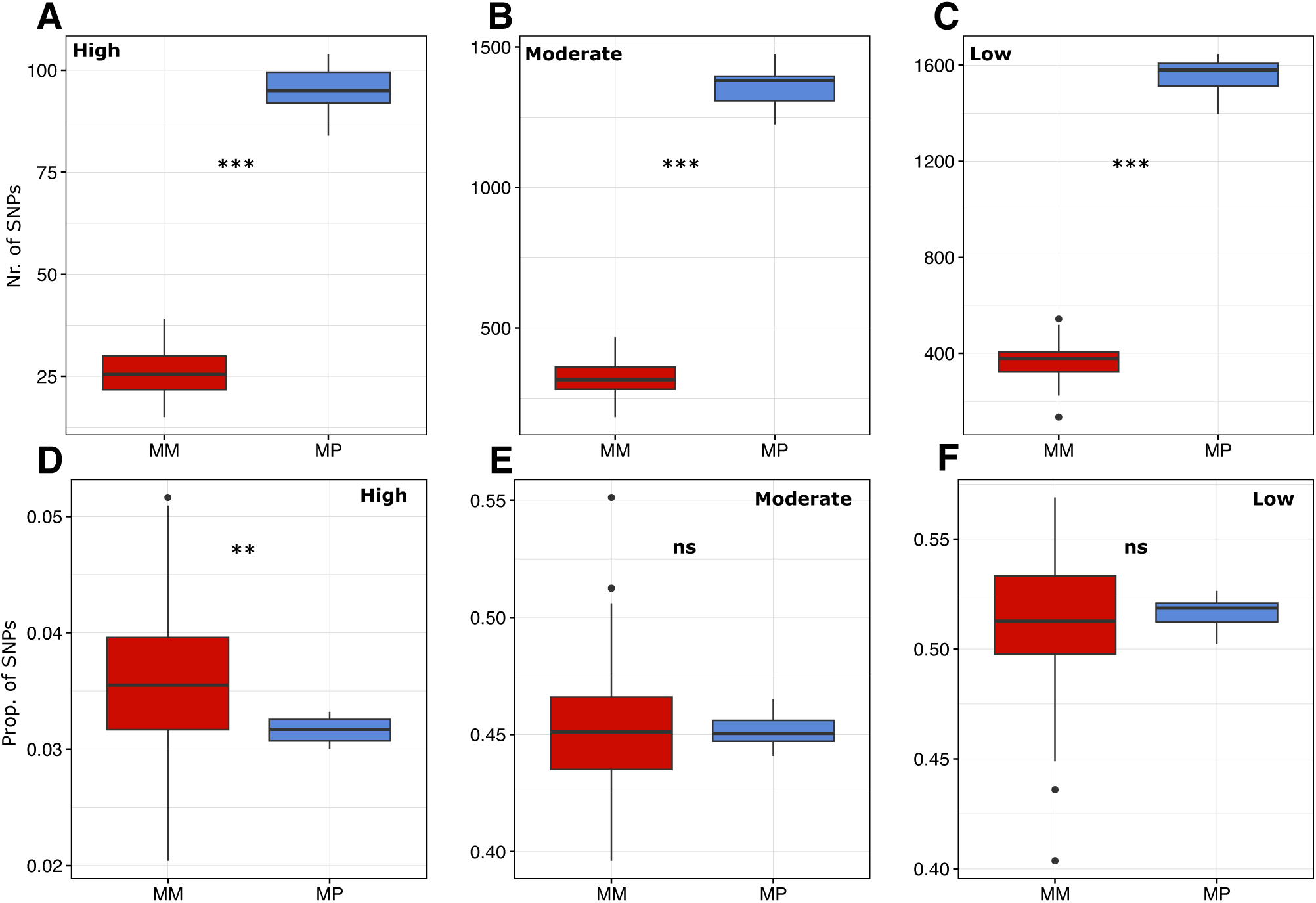
Predictions of the effect of intragenic variants on gene function; (**A**, **D**) high (e.g., frame-shift variants, whole-exon deletions, stop-codons), (**B**, **E**) moderate (e.g., non-synonymous mutation, in-frame deletion, in-frame insertion) and (**C**, **F**) low (e.g., synonymous substitution). Only SNPs within the supergene region on chromosome 3 (1.9-11.6 Mb) are included. The line within each boxplot shows median values across M/M workers (n = 120) and M/P workers (n = 16). (**A**-**C**) The total number of genetic variants of each class. (**D**-**F**) The relative proportion of each class. The number of SNPs in each category and sample is also provided as Table S11.

## Conclusion

Supergenes are powerful study systems for revealing candidate genes controlling complex phenotypic traits. However, their non-recombining nature results in large haplotypes (often containing hundreds of genes) typically being inherited as a single unit, which makes it hard to pinpoint the causal genes. The retention of only two P-specific genes in the obligate polygynous species *F. aquilonia* and *F. paralugubris* have, conveniently, narrowed down the list of candidate genes significantly. Future studies will reveal the precise function of the *Zasp52* and *TTLL2* genes in *Formica* ants, and their potential role in upholding the polygyny syndrome in obligate polygynous species lacking the full P haplotype.

## Materials and Methods

### Supergene haplotype distributions of seven wood ant species

We analyzed Illumina paired-end data from 145 individuals within the wood ant complex. Of these samples, three are previously published haploid males with known supergene status (n = 1 *F. pratensis* with a P haplotype, n = 1 *F. truncorum* with a M haplotype, and n = 1 *F. truncorum* with a P haplotype; Table S1; Brelsford et al., 2020). The remaining samples are diploid workers (females) with unknown supergene distributions, of the following species: *F. aquilonia* (n = 65), *F. lugubris* (n = 19), *F. paralugubris* (n = 20), *F. polyctena* (n = 6), *F. pratensis* (n = 26) and *F. rufa* (n = 6) (Table S1). Of the 142 workers, 72 have been previously described and published (Portinha et al., 2022; Satokangas et al., 2023) while the remaining 70 samples are new for this study (Table S1). Details about DNA extraction method and sequencing technology for all non-published samples are in Table S1.

The data from the three males were trimmed using trimmomatic v0.39 (Bolger et al., 2014) with options “SLIDINGWINDOW:4:20 MINLEN:25 ILLUMINACLIP:adapters/TruSeq3-PE-2.fa:2:40:15”. All samples were then aligned to the *Formica aquilonia* x *polyctena* reference genome (constructed from a haploid male hybrid individual; Nouhaud et al., 2022) using bwa mem v0.7.17, sorted using samtools sort v1.16.1 (Danecek et al., 2021) and deduplicated using picard MarkDuplicates v2.27 (http://broadinstitute.github.io/picard/). The resulting BAM files were filtered to only include reads mapping to chromosome 3 (Scaffold03) using samtools. We used deepvariant v1.4.0 (Poplin et al., 2018) to call variants for all BAM files separately (n = 145) with option --model_type WGS, and then performed joint variant calling using GLnexus v1.4.1-0-g68e25e5 (Yun et al., 2021) using options --config DeepVariantWGS and --trim-uncalled-alleles. We normalized the resulting VCF file using bcftools norm v1.16, and filtered variants using vcftools v0.1.17 (Danecek et al., 2011) with options --min-alleles 2 --max-alleles 2 --minQ 20 --minDP 3 --non-ref-ac 5 --max-missing 0.6 --mac 5 -- remove-filtered-all. We excluded samples with missing genotypes on ≥10% of called sites (n = 6), leaving 139 individuals (n = 136 workers; n = 3 males) with read depths between 9.1x and 59.9x (Table S2).

We calculated inbreeding coefficients (F_IS_ scores) for all samples using vcftools with option --het to identify samples that are supergene heterozygous (M/P). Of the 136 workers, 16 had extremely low F values (< –0.5), meaning that the observed homozygous sites were fewer than expected. These 16 samples (n = 5 *F. lugubris*, n = 3 *F. polyctena* and n = 8 *F. pratensis*) were labeled as M/P genotypes (Table S1). To determine the supergene haplotype of the other samples, we then filtered the VCF file keeping sites where all 16 M/P workers were heterozygous while allowing <20% missing genotypes. This left 21,790 trans-species haplotype-specific SNPs. For all samples, we then counted the proportion of reference homozygous sites (0/0), heterozygous sites (0/1) and alternative homozygous sites (1/1) using bcftools v1.16 (Danecek et al., 2021) query with the flag -f ‘[%SAMPLE\t%GT\n]’.

### Species phylogeny

To construct a species phylogeny, we called joint variants from the deepvariant VCF files again using GLnexus (see above) for the 139 samples, this time with two additional outgroup individuals from *Formica exsecta* (Table S1). These outgroup samples were trimmed and aligned in the same way as the male samples. From the joint VCF file, we extracted SNPs from chromosome 3 outside of the supergene region (>10kb away from chromosome ends, and >200kb away from the supergene region) using vcftools. We also excluded any genetic variants with heterozygosity in all 16 M/P individuals, leaving 37,862 SNPs. These were filtered using vcftools with the following options: --min-alleles 2 --max-alleles 2 --minQ 20 --minDP 3 --max-missing 0.3 --remove-filtered-all --remove-indels. The remaining 31,911 SNPs were filtered for minimum 100 bp distance using vcftools option –thin 100, leaving 7,805 SNPs. We used the script convert_vcf_to_nexus.rb (https://github.com/mmatschiner/tutorials/blob/master/species_tree_inference_with_snp_data/src/convert_vc f_to_nexus.rb; accessed 26^th^ of June, 2024) to convert the VCF file to NEXUS format, from which we constructed a phylogenetic tree for all samples using the svdQuartets (Chifman & Kubatko, 2014) algorithm, implemented in PAUP v4.0a168 (Swofford & Sullivan, 2003). We excluded 16 samples which did not cluster monophyletically in the resulting tree (Figure S7; n = 6 *F. aquilonia*, n = 7 F. lugubris, n = 1 *F. paralugubris*, n = 2 *F. polyctena*) from the VCF file containing the 7,805 SNPs (see above) and re-converted the output VCF to NEXUS format. We then used svdQuartets to construct a species-level phylogeny using a taxon partition file with the standard number of bootstraps.

### P-specific SNPs in obligate polygynous species with M/M genotypes

For each sample (n = 139), we counted the number of bi-allelic trans-species haplotype-specific SNPs (see above) occurring in CDS regions (n = 1,546 SNPS) with bcftools query using the flag -f ‘[%SAMPLE\t%GT\n]’. We interpreted a site as “P-specific” if the SNP genotype was either “0/1” or “1/1”. We used linear models in R v4.2.1 (R Core Team, 2022) to test if any genes had significantly different numbers of P-specific SNPs between M/M workers of obligate polygynous species (n = 3) and socially polymorphic species (n = 3). We also calculated sequencing depth for each CDS region and all samples using bamstats04 from Jvarkit v36b5fa3 (Lindenbaum, 2015). These values were normalized between samples, and the per-sample average was calculated for each gene. Two genes (“jg3505” and jg3507” in the *Formica aquilonia x polyctena* gene annotation; Table S6) had significantly more P-specific SNPs in the obligate polygynous species.

We functionally annotated these two genes in the following way: First, we identified orthologs between the *Formica aquilonia x polyctena* reference genome, two outgroup ant species (*Camponotus floridanus*, GCF_003227725.1; *Solenopsis invicta*, GCF_016802725.1) and a fruit fly (*Drosophila melanogaster*, GCF_000001215.4) using orthofinder v2.5.4 (Emms & Kelly, 2019). Secondly, using the two resulting orthogroups (containing jg3505 and jg3507, respectively), we confirmed that all transcripts were orthologous also in the orthoDB v11 (Kuznetsov et al., 2023) database. And thirdly, we confirmed the gene structure by blasting the gene sequences, as well as those of adjacently positioned gene annotations, on the NCBI website. We found that jg3504 and jg3505 belong to the same gene (*Zasp52*; Table S6) and manually annotated these into a new gene annotation (*Zasp52*; Table S7), using additional evidence from downloaded amino acid sequences from orthoDB (accessed 12^th^ of March; see orthogroup ID in Table S6).

The transcript jg3507, which belongs to the *TTLL2* gene (Table S6), and *Zasp52*, are both putative P-specific duplications (see Results and Discussion). To see if the duplication covers the entire transcripts, we extracted sequencing depth values using jvarkit (as above) for all SNPs remaining after the quality filtering (see above) within CDS regions of these genes. We found that individuals carrying a P haplotype, and workers of obligate polygynous species with M/M genotypes, had higher sequencing depth across the entire *TTLL2* gene (Figure S3, S4) but only across parts of the *Zasp52* gene (exons 10-16 in the curated transcript “Zasp52”; Figure S3, S4; Table S7). We therefore created a separate transcript which contained only exons 10-16 and adjusted the first exon to include an additional eight codons where the first one was a start codon (“Zasp52-dup”; Figure S8; Table S7). These eight codons were included to aid the homology-based annotation (see below) and were not included in any of the results in the Results and Discussion section. Note also that the curated transcripts Zasp52 and Zasp52-dup contain the same trans-species haplotype-specific SNPs as the original transcript jg3505 (n = 28; Figure S8).

### Confirming duplication of Zasp52 and TTLL2 using de novo assemblies of F. aquilonia and F. polyctena

To verify that the P-specific copies of *Zasp52* and *TTLL2* are duplicated on the M haplotype in the obligate polygynous species, we aligned raw PacBio data from (a) the *Formica aquilonia x polyctena* male from which the reference genome was constructed, (b) five pooled *F. aquilonia* males, and (c) five pooled *F. polyctena* males (Table S8), to the *Formica aquilonia x polyctena* reference genome using pbmm2 v1.13.0 (https://github.com/PacificBiosciences/pbmm2) with option --preset SUBREAD. The aligned reads were sorted and indexed using samtools, and genetic variants were called across Scaffold03 using longshot v.0.4.1 (Edge & Bansal, 2019). Of the 21,790 trans-species haplotype-specific SNPs (see above) only a small fraction of the SNPs from the PacBio data was heterozygotic or alternative homozygotic across the three datasets; 499 (2.3%) for the *Formica aquilonia x polyctena* dataset, 687 (3.2%) for the *F. aquilonia* dataset, and 958 (4.4%) for the *F. polyctena* dataset. We therefore concluded that the *F. aquilonia* and *F. polyctena* data exclusively contain the M haplotype, similarly to the *Formica aquilonia x polyctena* reference genome. We assembled the PacBio reads from all three datasets using Flye v.2.9.3 (Kolmogorov et al., 2019) with default parameters (genome statistics in Table S9). We used GeMoMa v1.6.4 (Keilwagen et al., 2019) to do homology-based annotations for all three *de novo* assemblies based on the annotation file for the *Formica aquilonia x polyctena* reference genome (https://doi.org/10.6084/m9.figshare.14186522.v1). For the *F. aquilonia* assembly, the genes *Zasp52* and *TTLL2* each occurred as three gene copies on separate contigs (Table S10). We ran the pipeline purge_dups v1.2.5 (Guan et al., 2020) on this assembly and found that contig_1719 was a haplotig of contig_189, and gene annotations from this contig were not used for further analysis (Table S10).

From the amino acid sequences of *Zasp52* and *TTLL2* downloaded from OrthoDB (see above), we selected all ant species (n = 22) and removed short sequences (≥ 600 for *Zasp52* and ≥ 400 for *TTLL2*; cut-offs based on manual inspection) using seqkit v0.16.0 (Shen et al., 2016). For each gene, we aligned these 22 sequences together with the predicted proteins from GeMoMa using clustalo v1.2.4 (Sievers & Higgins, 2018) with option -t Protein. The aligned sequences were manually inspected and then trimmed twice using trimAL v1.4.rev15 (Capella-Gutiérrez et al., 2009), first using the option -strictplus and then using the option -gt 0.8. Note that this trimming removed the first 8 amino acids in the beginning of the first exon of the transcript Zasp52_dup, which were adjusted to include a start codon. The final alignments were 1080 (*Zasp52*) and 470 (*TTLL2*) amino acids long. We used iqtree2 v2.2.2.7 (Minh et al., 2020) to create phylogenetic trees from the trimmed alignments using options -m LG+F+G -nt AUTO. We calculated pairwise sequence identity between all sequences using a custom python script. We determined which gene copies in the *de novo* assemblies were the duplicated paralogs based on the topology between the sequences and sequence similarity to the *Formica aquilonia x polyctena* reference genome (Figure 4; Table S10).

We trimmed, aligned, and called variants from n = 5 published *Formica* samples outside the wood ant complex (Table S1) using the same method as for the other Illumina samples (see above). We extracted SNPs at the positions of the wood ant trans-species haplotype-specific SNPs occurring in CDS regions using bcftools.

### Mutation load

The genome assembly GFF3 file was transformed to GTF2.2 format using the script agat_convert_sp_gff2gtf.pl v.1.0.0 from agat v1.0.0 (Dainat, n.d.), and protein and CDS sequence files were created from this file using gffread v0.12.7 (Pertea & Pertea, 2020). A snpEff v5.2 (Cingolani et al., 2012) database was build based on these files. From the filtered VCF file containing 139 individuals (see above), we extracted SNPs from each worker using vcftools and the option –keep <IND> --mac 1. The functional effects of all SNPs were predicted using snpEff ann. From this output, we retained genetic variants from primary transcripts with high, moderate, and low predicted effects.

### Alignment, variant calling, and expression levels of RNA-seq data

We indexed the *Formica aquilonia* x *polyctena* and the *F. aquilonia* reference genomes using STAR v2.7.11a (Dobin et al., 2013) with option --sjdbOverhang 149. We aligned the RNA data from a pool of 20 *F. aquilonia* workers (read length 150bp; this study) to both reference genomes using STAR with option --twopassMode Basic. The reads were sorted and indexed using sambamba v0.7.0 (Tarasov et al., 2015), duplicates were marked using picard MarkDuplicates v2.27.5, and GATK SplitNCigarReads v4.5.0.0 (Van der Auwera et al., 2020) was used to split reads. We called variants for each sample separately using GATK Haplotypecaller. We counted aligned reads per gene using featureCounts v.2.0.1 (Y. Liao et al., 2014) and calculated transcripts per million (TPM) per gene using the following formula:

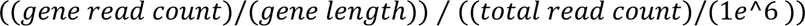

We defined the expression level of genes based on the cut-off values used by the EMBL-EBI Expression Atlas (last accessed 20^th^ March 2024).

## Data Availability

All code for running these analyses is available as Supplementary Code, and supporting DNA sequences will be uploaded with the published manuscript.

## Author Contributions

H.S. and L.V. conceived the study. H.S. performed the bioinformatic analyses and wrote the initial draft of the manuscript with critical input from all co-authors I.S., P.N., H.H., J.K., M.D.L., R.D., and M.C. provided data or samples, performed molecular lab work, and/or contributed to data interpretation. L.V. secured funding for the project. All authors reviewed and approved the final version of the manuscript.

## Acknowledgements

This work was supported by the Research Council of Finland (to L.V., grant no. 24303855) and the Swiss National Science Foundation (to M.C., grant no. 310030_207642).

## References

Blacher, P., De Gasperin, O., Grasso, G., Sarton-Lohéac, S., Allemann, R., & Chapuisat, M. (2023). Cryptic recessive lethality of a supergene controlling social organization in ants. Molecular Ecology, 32(5), 1062– 1072. 10.1111/mec.16821

Bolger, A. M., Lohse, M., & Usadel, B. (2014). Trimmomatic: A flexible trimmer for Illumina sequence data. Bioinformatics, 30(15), 2114–2120. 10.1093/bioinformatics/btu170

Boomsma, J. J., Huszár, D. B., & Pedersen, J. S. (2014). The evolution of multiqueen breeding in eusocial lineages with permanent physically differentiated castes. Animal Behaviour, 92, 241–252. 10.1016/j.anbehav.2014.03.005

Brelsford, A., Purcell, J., Avril, A., Tran Van, P., Zhang, J., Brütsch, T., Sundström, L., Helanterä, H., & Chapuisat, M. (2020). An Ancient and Eroded Social Supergene Is Widespread across Formica Ants. Current Biology, 30(2), 304–311.e4. 10.1016/j.cub.2019.11.032

Capella-Gutiérrez, S., Silla-Martínez, J. M., & Gabaldón, T. (2009). trimAl: A tool for automated alignment trimming in large-scale phylogenetic analyses. Bioinformatics, 25(15), 1972–1973. 10.1093/bioinformatics/btp348

Chapuisat, M., Bocherens, S., & Rosset, H. (2004). Variable queen number in ant colonies: No impact on queen turnover, inbreeding, and population genetic differentiation in the ant Formica selysi. Evolution; International Journal of Organic Evolution, 58(5), 1064–1072. 10.1111/j.0014-3820.2004.tb00440.x

Chapuisat, M., Goudet, J., & Keller, L. (1997). MICROSATELLITES REVEAL HIGH POPULATION VISCOSITY AND LIMITED DISPERSAL IN THE ANT *FORMICA PARALUGUBRIS*. Evolution, 51(2), 475–482. 10.1111/j.1558-5646.1997.tb02435.x

Charlesworth, D. (2016). The status of supergenes in the 21st century: Recombination suppression in B atesian mimicry and sex chromosomes and other complex adaptations. Evolutionary Applications, 9(1), 74–90. 10.1111/eva.12291

Chechenova, M. B., Bryantsev, A. L., & Cripps, R. M. (2013). The Drosophila Z-disc Protein Z(210) Is an Adult Muscle Isoform of Zasp52, Which Is Required for Normal Myofibril Organization in Indirect Flight Muscles. Journal of Biological Chemistry, 288(6), 3718–3726. 10.1074/jbc.M112.401794

Chifman, J., & Kubatko, L. (2014). Quartet Inference from SNP Data Under the Coalescent Model. Bioinformatics, 30(23), 3317–3324. 10.1093/bioinformatics/btu530

Cingolani, P., Platts, A., Wang, L. L., Coon, M., Nguyen, T., Wang, L., Land, S. J., Lu, X., & Ruden, D. M. (2012). A program for annotating and predicting the effects of single nucleotide polymorphisms, SnpEff: SNPs in the genome of Drosophila melanogaster strain w 1118; iso-2; iso-3. Fly, 6(2), 80–92. 10.4161/fly.19695

Clarke, C. A., & Sheppard, P. M. (1960). Super-genes and mimicry. Heredity, 14(1–2), 175–185. 10.1038/hdy.1960.15

Dainat, J. (n.d.). AGAT: Another Gff Analysis Toolkit to handle annotations in any GTF/GFF format. (Version v0.8.0) [Dataset]. Zenodo. https://www.doi.org/10.5281/zenodo.3552717

Danecek, P., Auton, A., Abecasis, G., Albers, C. A., Banks, E., DePristo, M. A., Handsaker, R. E., Lunter, G., Marth, G. T., Sherry, S. T., McVean, G., Durbin, R., & 1000 Genomes Project Analysis Group. (2011). The variant call format and VCFtools. Bioinformatics, 27(15), 2156–2158. 10.1093/bioinformatics/btr330

Danecek, P., Bonfield, J. K., Liddle, J., Marshall, J., Ohan, V., Pollard, M. O., Whitwham, A., Keane, T., McCarthy, S. A., Davies, R. M., & Li, H. (2021). Twelve years of SAMtools and BCFtools. GigaScience, 10(2), giab008. 10.1093/gigascience/giab008

DeHeer, C. J., & Herbers, J. M. (2004). Population genetics of the socially polymorphic ant Formica podzolica. Insectes Sociaux, 51(4), 309–316. 10.1007/s00040-004-0745-1

Dobin, A., Davis, C. A., Schlesinger, F., Drenkow, J., Zaleski, C., Jha, S., Batut, P., Chaisson, M., & Gingeras, T. R. (2013). STAR: Ultrafast universal RNA-seq aligner. Bioinformatics, 29(1), 15–21. 10.1093/bioinformatics/bts635

Edge, P., & Bansal, V. (2019). Longshot enables accurate variant calling in diploid genomes from single-molecule long read sequencing. Nature Communications, 10(1), 4660. 10.1038/s41467-019-12493-y

Emms, D. M., & Kelly, S. (2019). OrthoFinder: Phylogenetic orthology inference for comparative genomics. Genome Biology, 20(1), 238. 10.1186/s13059-019-1832-y

Fagerberg, L., Hallström, B. M., Oksvold, P., Kampf, C., Djureinovic, D., Odeberg, J., Habuka, M., Tahmasebpoor, S., Danielsson, A., Edlund, K., Asplund, A., Sjöstedt, E., Lundberg, E., Szigyarto, C. A.-K., Skogs, M., Takanen, J. O., Berling, H., Tegel, H., Mulder, J., … Uhlén, M. (2014). Analysis of the Human Tissue-specific Expression by Genome-wide Integration of Transcriptomics and Antibody-based Proteomics. Molecular & Cellular Proteomics, 13(2), 397–406. 10.1074/mcp.M113.035600

Goropashnaya, A. V., Seppä, P., & Pamilo, P. (2001). Social and genetic characteristics of geographically isolated populations in the ant *Formica cinerea*. Molecular Ecology, 10(12), 2807–2818. 10.1046/j.0962-1083.2001.01410.x

Guan, D., McCarthy, S. A., Wood, J., Howe, K., Wang, Y., & Durbin, R. (2020). Identifying and removing haplotypic duplication in primary genome assemblies. Bioinformatics, 36(9), 2896–2898. 10.1093/bioinformatics/btaa025

Guerrero, R. F., Kirkpatrick, M., & Perrin, N. (2012). Cryptic recombination in the ever-young sex chromosomes of H ylid frogs. Journal of Evolutionary Biology, 25(10), 1947–1954. 10.1111/j.1420-9101.2012.02591.x

Hakala, S. M. (2020). Social polymorphism and dispersal in Formica ants. Helsingin yliopisto.

Jay, P., Chouteau, M., Whibley, A., Bastide, H., Parrinello, H., Llaurens, V., & Joron, M. (2021). Mutation load at a mimicry supergene sheds new light on the evolution of inversion polymorphisms. Nature Genetics, 53(3), 288–293. 10.1038/s41588-020-00771-1

John R. G. Turner. (1967). On Supergenes. I. The Evolution of Supergenes. The American Naturalist, 101(919), 195–221.

Johnson, C. A., Sundström, L., & Billen, J. (2005). Development of alary muscles in single-and multiple-queen populations of the wood ant Formica truncorum. Annales Zoologici Fennici, *42*(3,), 225–234.

Joron, M., Papa, R., Beltrán, M., Chamberlain, N., Mavárez, J., Baxter, S., Abanto, M., Bermingham, E., Humphray, S. J., Rogers, J., Beasley, H., Barlow, K., H. ffrench-Constant, R., Mallet, J., McMillan, W. O., & Jiggins, C. D. (2006). A Conserved Supergene Locus Controls Colour Pattern Diversity in Heliconius Butterflies. PLoS Biology, 4(10), e303. 10.1371/journal.pbio.0040303

Katzemich, A., Liao, K. A., Czerniecki, S., & Schöck, F. (2013). Alp/Enigma Family Proteins Cooperate in Z-Disc Formation and Myofibril Assembly. PLoS Genetics, 9(3), e1003342. 10.1371/journal.pgen.1003342

Keilwagen, J., Hartung, F., & Grau, J. (2019). GeMoMa: Homology-Based Gene Prediction Utilizing Intron Position Conservation and RNA-seq Data. In M. Kollmar (Ed.), Gene Prediction: Methods and Protocols (pp. 161–177). Springer New York. 10.1007/978-1-4939-9173-0_9

Keller, L. (1991). Queen number, mode of colony founding, and queen reproductive success in ants (Hymenoptera Formicidae). Ethology Ecology & Evolution, 3(4), 307–316. 10.1080/08927014.1991.9525359

Keller, L. (1993). The Assessment of Reproductive Success of Queens in Ants and Other Social Insects. Oikos, 67(1), 177. 10.2307/3545107

Kimura, M. (1956). A MODEL OF A GENETIC SYSTEM WHICH LEADS TO CLOSER LINKAGE BY NATURAL SELECTION. Evolution, 10(3), 278–287. 10.1111/j.1558-5646.1956.tb02852.x

Kolmogorov, M., Yuan, J., Lin, Y., & Pevzner, P. A. (2019). Assembly of long, error-prone reads using repeat graphs. Nature Biotechnology, 37(5), 540–546. 10.1038/s41587-019-0072-8

Küpper, C., Stocks, M., Risse, J. E., Dos Remedios, N., Farrell, L. L., McRae, S. B., Morgan, T. C., Karlionova, N., Pinchuk, P., Verkuil, Y. I., Kitaysky, A. S., Wingfield, J. C., Piersma, T., Zeng, K., Slate, J., Blaxter, M., Lank, D. B., & Burke, T. (2016). A supergene determines highly divergent male reproductive morphs in the ruff. Nature Genetics, 48(1), 79–83. 10.1038/ng.3443

Kuznetsov, D., Tegenfeldt, F., Manni, M., Seppey, M., Berkeley, M., Kriventseva, E. V., & Zdobnov, E. M. (2023). OrthoDB v11: Annotation of orthologs in the widest sampling of organismal diversity. Nucleic Acids Research, 51(D1), D445–D451. 10.1093/nar/gkac998

Lagunas-Robles, G., Purcell, J., & Brelsford, A. (2021). Linked supergenes underlie split sex ratio and social organization in an ant. Proceedings of the National Academy of Sciences, 118(46), e2101427118. 10.1073/pnas.2101427118

Lamichhaney, S., Fan, G., Widemo, F., Gunnarsson, U., Thalmann, D. S., Hoeppner, M. P., Kerje, S., Gustafson, U., Shi, C., Zhang, H., Chen, W., Liang, X., Huang, L., Wang, J., Liang, E., Wu, Q., Lee, S. M.-Y., Xu, X., Höglund, J., … Andersson, L. (2016). Structural genomic changes underlie alternative reproductive strategies in the ruff (Philomachus pugnax). Nature Genetics, 48(1), 84–88. 10.1038/ng.3430

Liao, K. A., González-Morales, N., & Schöck, F. (2016). Zasp52, a Core Z-disc Protein in Drosophila Indirect Flight Muscles, Interacts with α-Actinin via an Extended PDZ Domain. PLOS Genetics, 12(10), e1006400. 10.1371/journal.pgen.1006400

Liao, Y., Smyth, G. K., & Shi, W. (2014). featureCounts: An efficient general purpose program for assigning sequence reads to genomic features. Bioinformatics, 30(7), 923–930. 10.1093/bioinformatics/btt656

Lindenbaum, P. (2015). JVarkit: Java-based utilities for Bioinformatics [Dataset]. 10.6084/m9.figshare.1425030

Llaurens, V., Whibley, A., & Joron, M. (2017). Genetic architecture and balancing selection: The life and death of differentiated variants. Molecular Ecology, 26(9), 2430–2448. 10.1111/mec.14051

McGuire, D., Sankovitz, M., & Purcell, J. (2022). A novel distribution of supergene genotypes is present in the socially polymorphic ant Formica neoclara. BMC Ecology and Evolution, 22(1), 47. 10.1186/s12862-022-02001-0

Minh, B. Q., Schmidt, H. A., Chernomor, O., Schrempf, D., Woodhams, M. D., Von Haeseler, A., & Lanfear, R. (2020). IQ-TREE 2: New Models and Efficient Methods for Phylogenetic Inference in the Genomic Era. Molecular Biology and Evolution, 37(5), 1530–1534. 10.1093/molbev/msaa015

Nouhaud, P., Beresford, J., & Kulmuni, J. (2022). Assembly of a Hybrid *Formica aquilonia* × *F. polyctena* Ant Genome From a Haploid Male. Journal of Heredity, 113(3), 353–359. 10.1093/jhered/esac019

Pamilo, P., Chautems, D., & Cherix, D. (1992). Genetic differentiation of disjunct populations of the antsFormica aquilonia andFormica lugubris in Europe. Insectes Sociaux, 39(1), 15–29. 10.1007/BF01240528

Pertea, G., & Pertea, M. (2020). GFF Utilities: GffRead and GffCompare. F1000Research, 9, ISCB Comm J-304. 10.12688/f1000research.23297.2

Pierce, D., Sun, P., Purcell, J., & Brelsford, A. (2022). A socially polymorphic *Formica* ant species exhibits a novel distribution of social supergene genotypes. Journal of Evolutionary Biology, 35(8), 1031–1044. 10.1111/jeb.14038

Poplin, R., Chang, P.-C., Alexander, D., Schwartz, S., Colthurst, T., Ku, A., Newburger, D., Dijamco, J., Nguyen, N., Afshar, P. T., Gross, S. S., Dorfman, L., McLean, C. Y., & DePristo, M. A. (2018). A universal SNP and small-indel variant caller using deep neural networks. Nature Biotechnology, 36(10), 983–987. 10.1038/nbt.4235

Portinha, B., Avril, A., Bernasconi, C., Helanterä, H., Monaghan, J., Seifert, B., Sousa, V. C., Kulmuni, J., & Nouhaud, P. (2022). Whole-genome analysis of multiple wood ant population pairs supports similar speciation histories, but different degrees of gene flow, across their European ranges. Molecular Ecology, 31(12), 3416–3431. 10.1111/mec.16481

Purcell, J., Brelsford, A., Wurm, Y., Perrin, N., & Chapuisat, M. (2014). Convergent Genetic Architecture Underlies Social Organization in Ants. Current Biology, 24(22), 2728–2732. 10.1016/j.cub.2014.09.071

Purcell, J., Lagunas-Robles, G., Rabeling, C., Borowiec, M. L., & Brelsford, A. (2021). The maintenance of polymorphism in an ancient social supergene. Molecular Ecology, 30(23), 6246–6258. 10.1111/mec.16196

R Core Team. (2022). R: A Language and Environment for Statistical Computing. R Foundation for Statistical Computing, Vienna [Dataset]. https://www.R-project.org

Rosset, H., & Chapuisat, M. (2007). Alternative life-histories in a socially polymorphic ant. Evolutionary Ecology, 21(5), 577–588. 10.1007/s10682-006-9139-3

Satokangas, I., Nouhaud, P., Seifert, B., Punttila, P., Schultz, R., Jones, M. M., Sirén, J., Helanterä, H., & Kulmuni, J. (2023). Semipermeable species boundaries create opportunities for gene flow and adaptive potential. Molecular Ecology, 32(15), 4329–4347. 10.1111/mec.16992

Schwander, T., Libbrecht, R., & Keller, L. (2014). Supergenes and Complex Phenotypes. Current Biology, 24(7), R288–R294. 10.1016/j.cub.2014.01.056

Seifert, B. (2018). *The ants of Central and North Europe* (E. J. H. Robinson, Trans.; 1. Auflage). lutra.

Seppä, P., Gyllenstrand, N., Corander, J., & Pamilo, P. (2004). COEXISTENCE OF THE SOCIAL TYPES: GENETIC POPULATION STRUCTURE IN THE: ANT FORMICA EXSECTA. Evolution, 58(11), 2462–2471. 10.1111/j.0014-3820.2004.tb00875.x

Shen, W., Le, S., Li, Y., & Hu, F. (2016). SeqKit: A Cross-Platform and Ultrafast Toolkit for FASTA/Q File Manipulation. PLOS ONE, 11(10), e0163962. 10.1371/journal.pone.0163962

Sievers, F., & Higgins, D. G. (2018). Clustal Omega for making accurate alignments of many protein sequences. Protein Science, 27(1), 135–145. 10.1002/pro.3290

Stolle, E., Pracana, R., López-Osorio, F., Priebe, M. K., Hernández, G. L., Castillo-Carrillo, C., Arias, M. C., Paris, C. I., Bollazzi, M., Priyam, A., & Wurm, Y. (2022). Recurring adaptive introgression of a supergene variant that determines social organization. Nature Communications, 13(1), 1180. 10.1038/s41467-022-28806-7

Sundström, L. (1993). Genetic population structure and sociogenetic organisation in Formica truncorum (Hymenoptera; Formicidae). Behavioral Ecology and Sociobiology, 33(5), 345–354. 10.1007/BF00172934

Swofford, D. L., & Sullivan, J. (2003). Phylogeny inference based on parsimony and other methods using PAUP*. *The Phylogenetic Handbook: A Practical Approach to DNA and Protein Phylogeny*, Cáp, 7, 160– 206.

Tafreshi, A. G., Otto, S. P., & Chapuisat, M. (2022). Unbalanced selection: The challenge of maintaining a social polymorphism when a supergene is selfish. Philosophical Transactions of the Royal Society B: Biological Sciences, 377(1856), 20210197. 10.1098/rstb.2021.0197

Tarasov, A., Vilella, A. J., Cuppen, E., Nijman, I. J., & Prins, P. (2015). Sambamba: Fast processing of NGS alignment formats. Bioinformatics, 31(12), 2032–2034. 10.1093/bioinformatics/btv098

Thompson, M. J., & Jiggins, C. D. (2014). Supergenes and their role in evolution. Heredity, 113(1), 1–8. 10.1038/hdy.2014.20

Van der Auwera, G., O’Connor, B., & Safari, an O. M. C. (2020). Genomics in the Cloud. O’Reilly Media, Incorporated. https://books.google.se/books?id=dC42zgEACAAJ

Wang, J., Wurm, Y., Nipitwattanaphon, M., Riba-Grognuz, O., Huang, Y.-C., Shoemaker, D., & Keller, L. (2013). A Y-like social chromosome causes alternative colony organization in fire ants. Nature, 493(7434), 664—668. 10.1038/nature11832

Yun, T., Li, H., Chang, P.-C., Lin, M. F., Carroll, A., & McLean, C. Y. (2021). Accurate, scalable cohort variant calls using DeepVariant and GLnexus. Bioinformatics, 36(24), 5582–5589. 10.1093/bioinformatics/btaa1081

Zhang, W., Westerman, E., Nitzany, E., Palmer, S., & Kronforst, M. R. (2017). Tracing the origin and evolution of supergene mimicry in butterflies. Nature Communications, 8(1), 1269. 10.1038/s41467-017-01370-1

