## Supplementary Figures 1-8 for "The loss of a supergene in obligately polygynous *Formica* wood ant species"

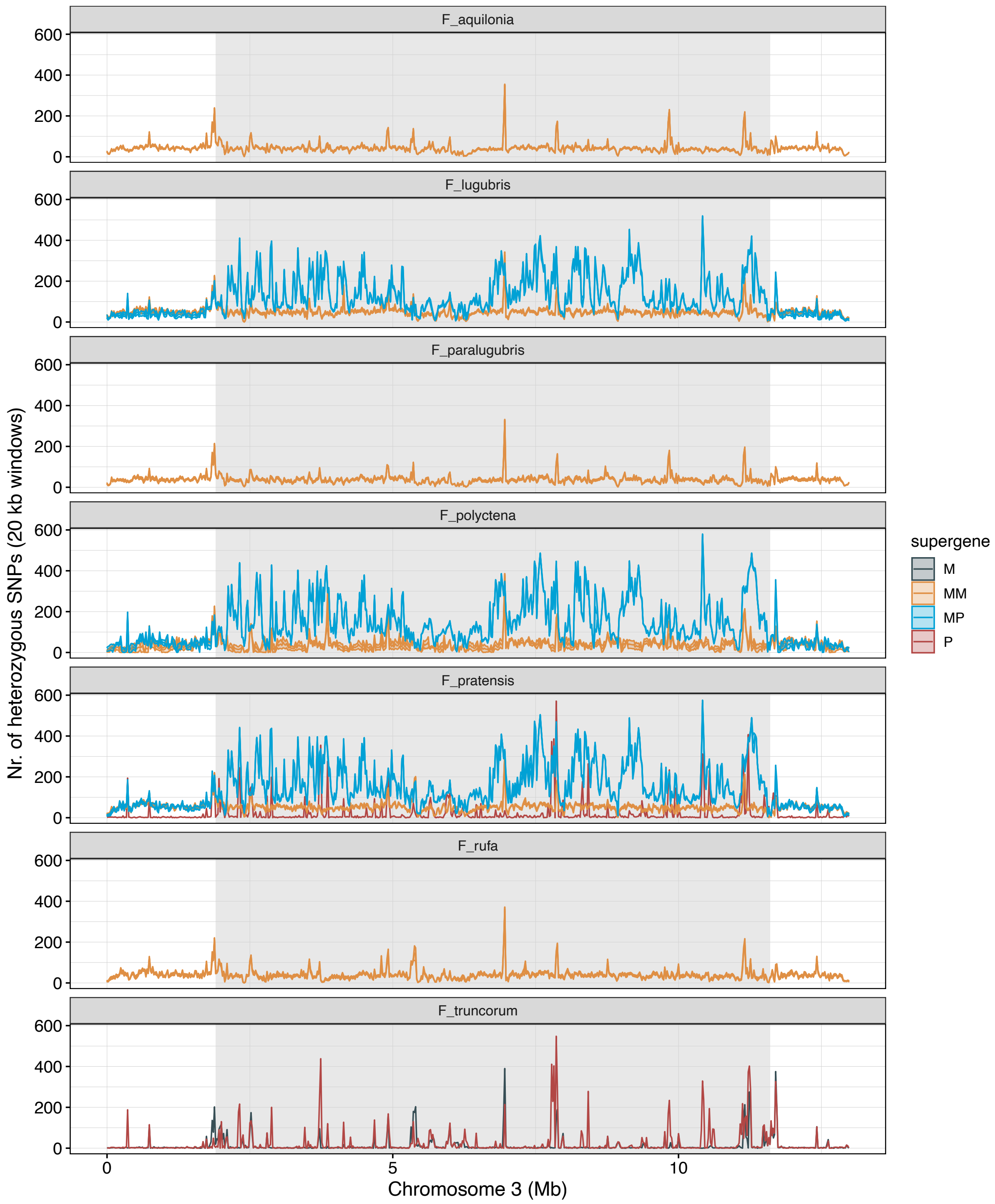


**Figure S1**. Mean values (±SE) of heterozygous SNPs (0/1) across 20 kb windows, plotted separately per species and supergene haplotype.

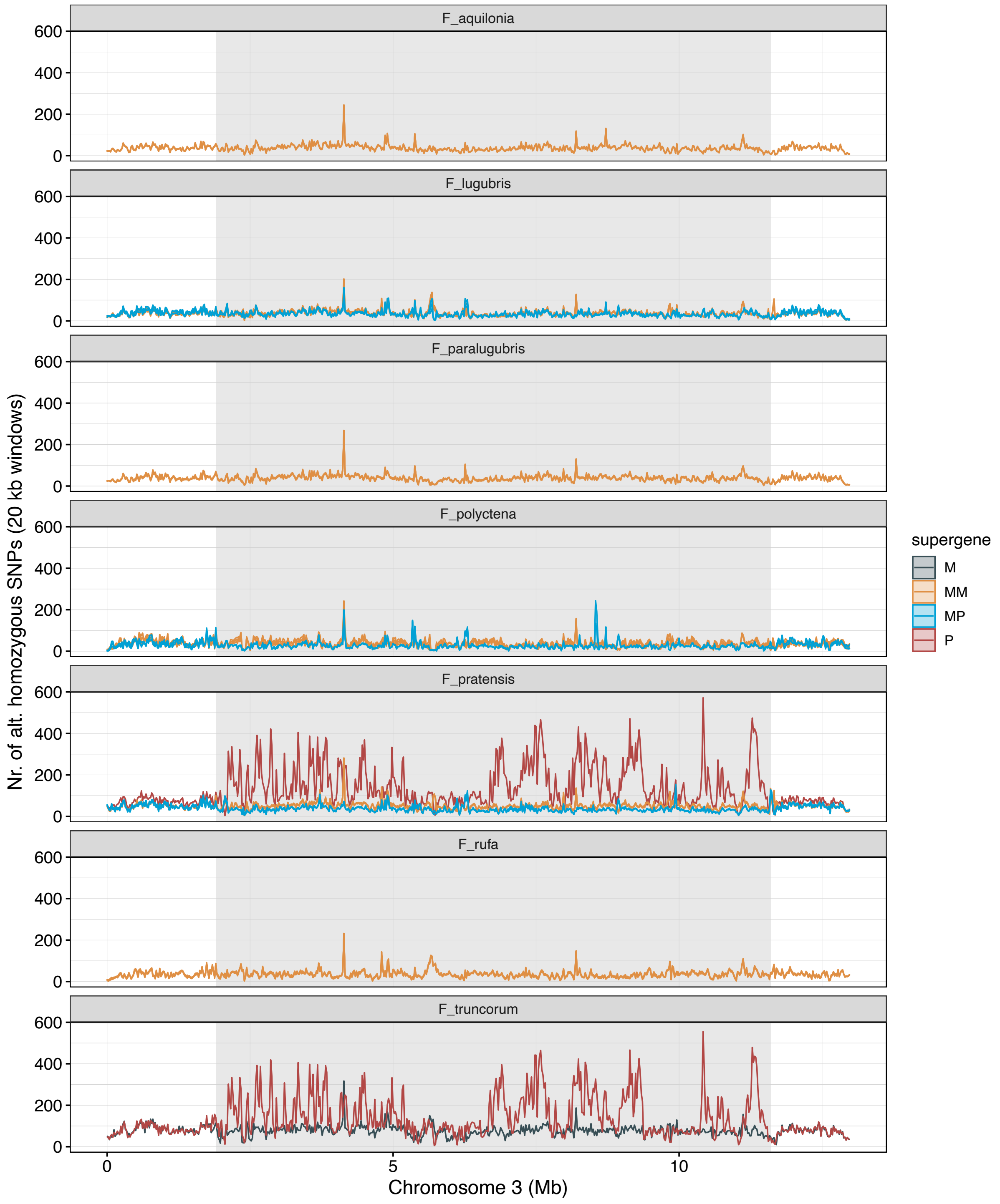


**Figure S2**. Mean values (±SE) of alternative homozygous SNPs (1/1) across 20 kb windows, plotted separately per species and supergene haplotype.

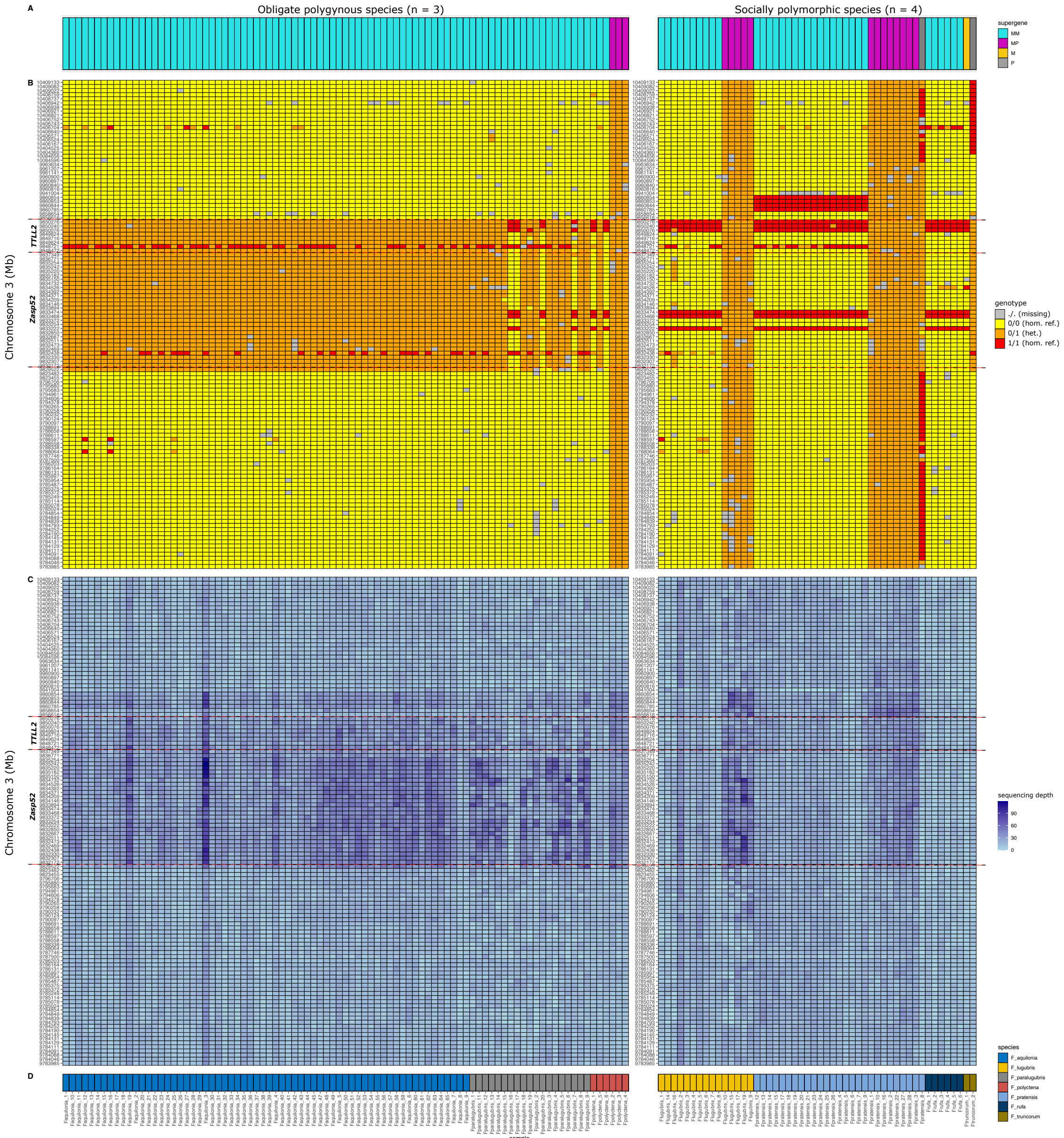


**Figure S3**. Heatmap showing (**A**) supergene genotype, (**B**) genotypes at the trans-species haplotype-specific SNPs in the genome region around *Zasp52* and *TTLL2* (Scaffold03:9782000-10410000, (**C**) sequencing depth values for the same genome region, and (**D**) species.

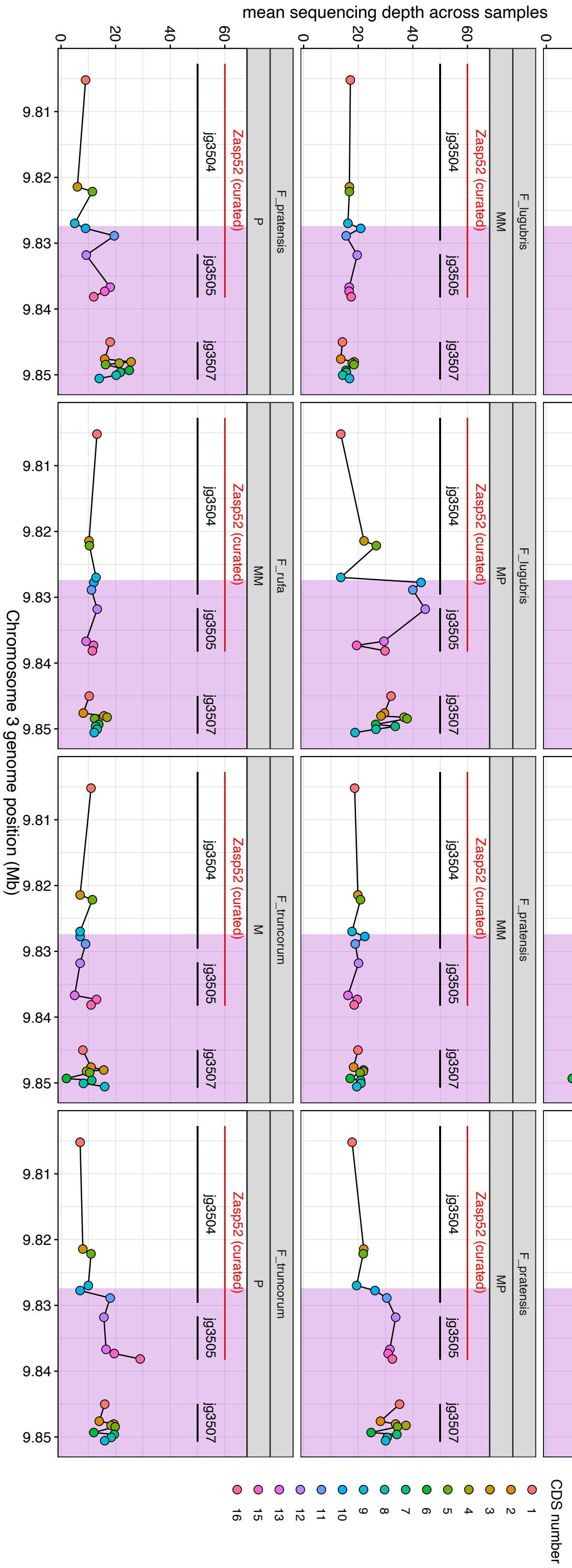


**Figure S4**. Per-CDS sequencing depth values for the genes *Zasp52* (jg3504 and jg3505) and *TTLL2* (jg3507). The values are averages across all SNPs in the region. The black lines mark the extent of the original transcripts and the red line show the extent of the manually curated *Zasp52* gene annotation.

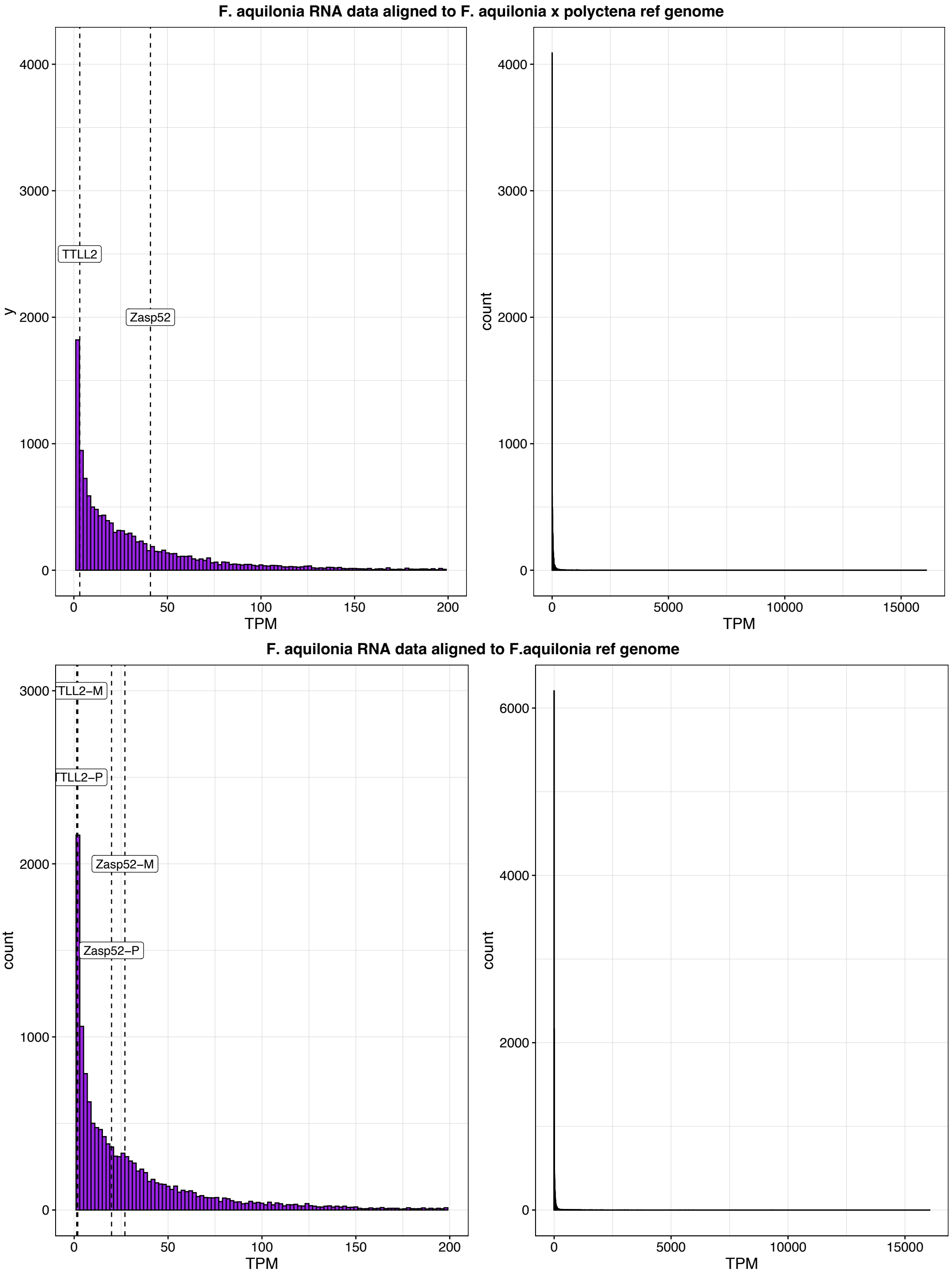


**Figure S5**. TPM values for the genes *TTLL2* and *Zasp52*, based on RNAseq data from 20 *F. aquilonia* workers aligned to (top) the *F. aquilonia x polyctena* and (bottom) the *de novo F. aquilonia* reference genome. The left-side panels show TPM values ≥200 (for increased visibility) while the right-side panels show all values.

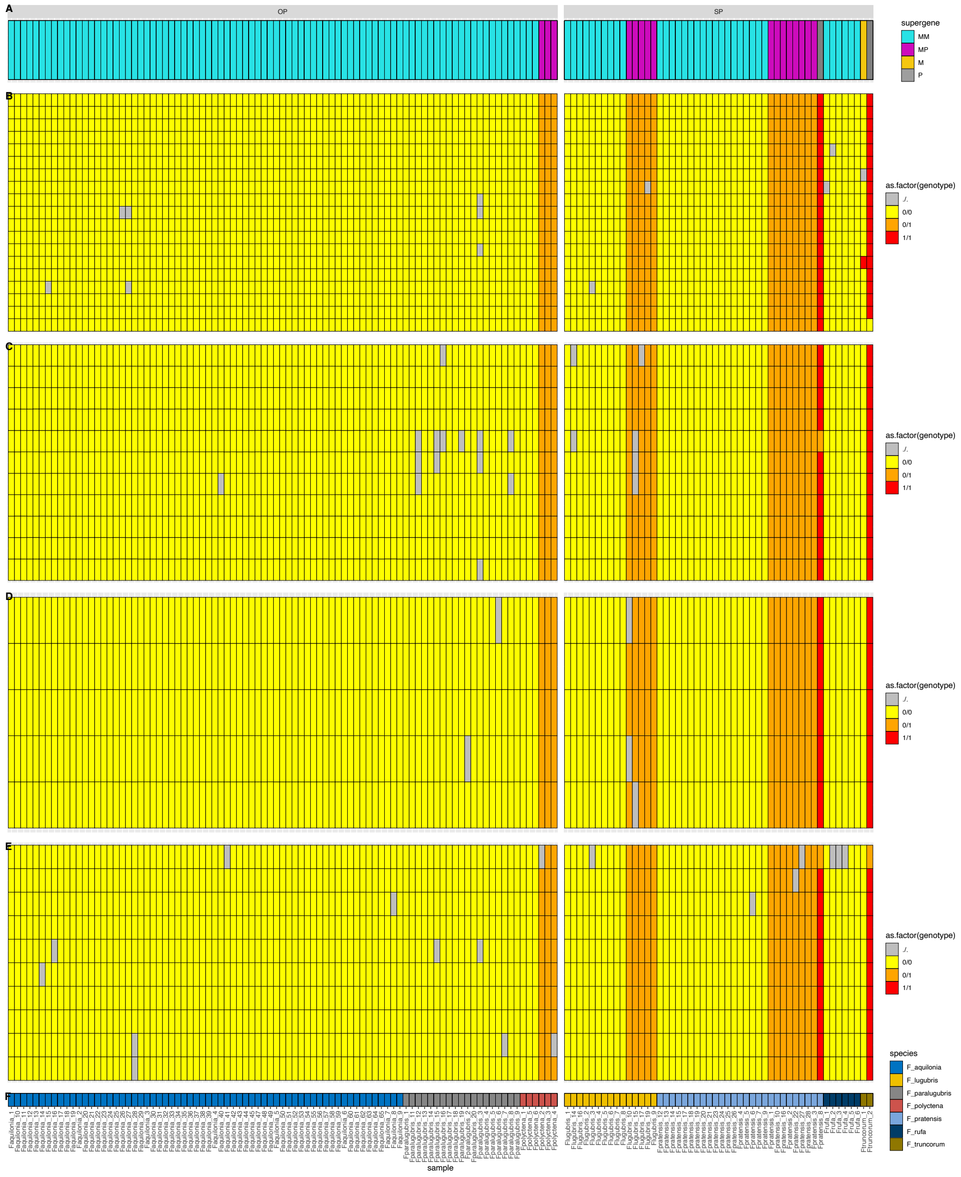


**Figure S6.** Heatmap showing (**A**) supergene genotype, and genotypes at the trans-species haplotype-specific SNPs at the genes (**B**) *Knockout*, (**C**) *Single-minded*, (**D**) *ZPF148*, and (**E**) *AmGR10*. The last row shows (**F**) Sample ID's and species.

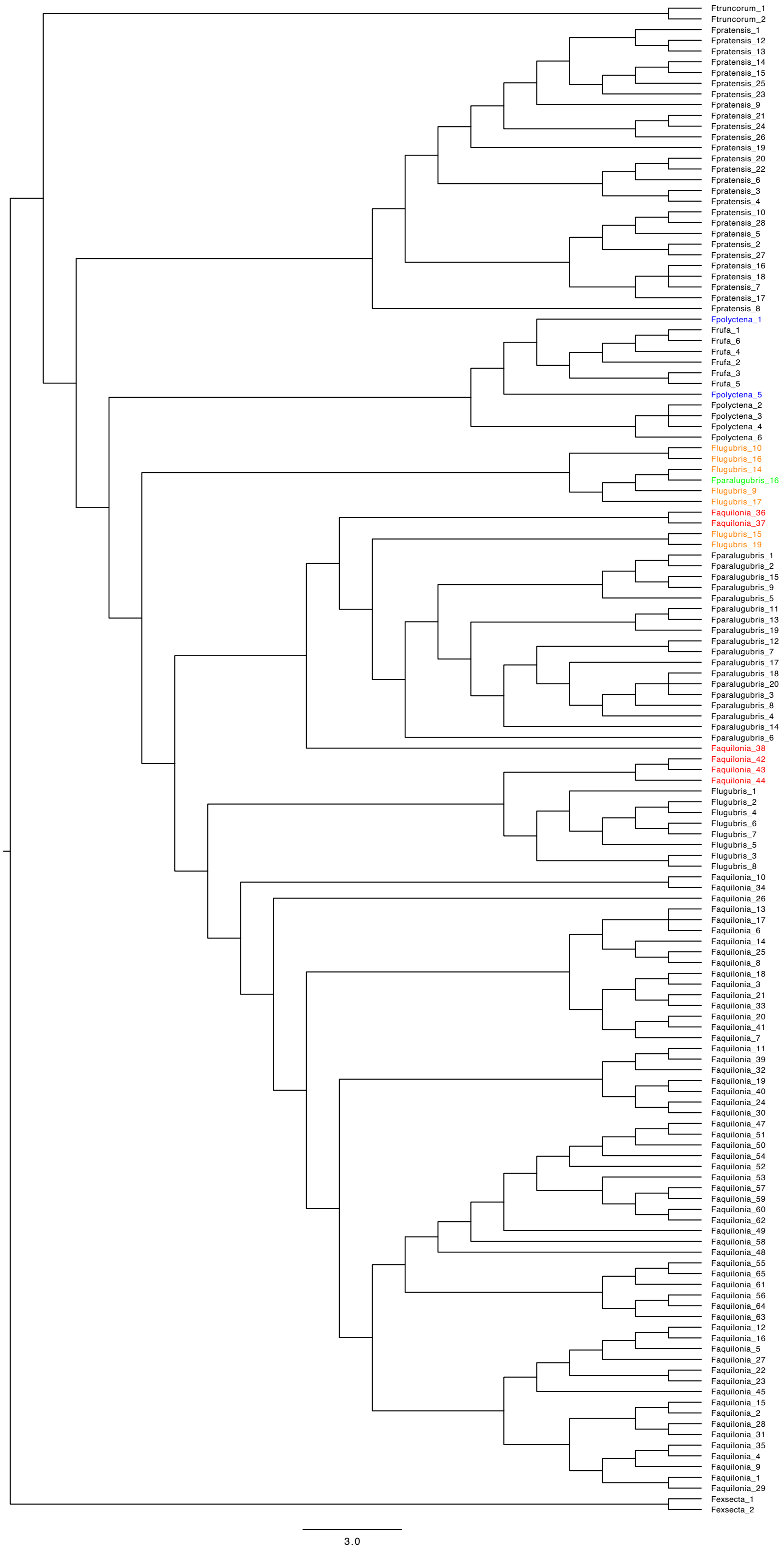


**Figure S7**. Output tree from svdQuartets of all 139 samples (Table S1). The highlighted samples (i.e., non-black tip labels) were removed from the input VCF file before constructing the final species tree (Figure 2a; see Methods).

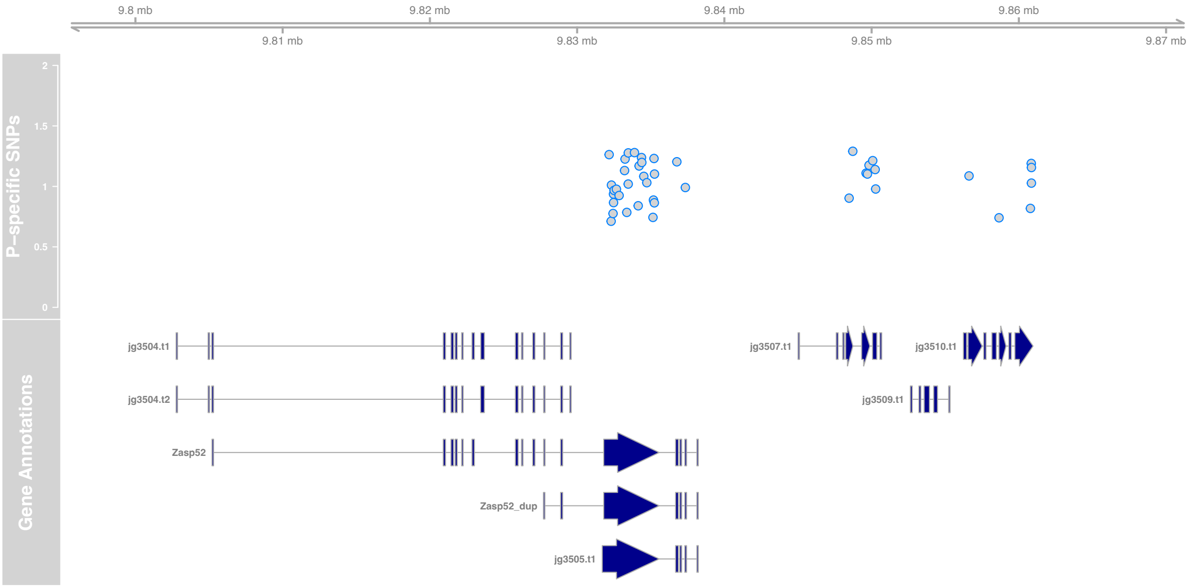


**Figure S8**. Original and curated gene models of *Zasp52* and *TTLL2* (see Main Text for details). The gene models starting with "jg" are from the Formica aquilonia x polyctena gene annotation, while the others are manually curated (provided in Table S7).
