## Supplementary Code for "The loss of a supergene in obligately polygynous *Formica* wood ant species": Supplementary Code 3702782bed6b4b8ca818cbea26b5421f.html

### Supplementary Code

> The code is arranged in the order it is presented in the Methods section in the Main Text.

**Supergene haplotype distributions of seven wood ant species**

Step 1: Run snakemake files available at https://github.com/hsigeman/wood-ant-supergene

```
git clone https://github.com/hsigeman/wood-ant-supergene
cd wood-ant-supergene

snakemake -s code/snakemake/snakefile_filtered -j 40 --cluster-config cluster_puhti_mem.yml --cluster " sbatch -A {cluster.account} -t {cluster.time} -c {cluster.cpus-per-task} --mem-per-cpu {cluster.mem-per-cpu}" -k
```

Step 2: Run code in the following page for supergene genotyping:

Code for supergene genotyping

Species phylogeny

Step 1: Run code in the following page to reproduce Figure 2a:

Code for phylogenetic tree (Figure 2a)

**P-specific SNPs in obligate polygynous species with M/M genotypes** 

Step 1: Run code in the following page to identify P-specific gene copies in M/M workers:

Code for CDS SNPs

Step 2: Run code in the following page to run an orthology analysis between the F. aquilonia x polyctena reference genome and outgroup insect genomes:

Code for orthology analysis

**Confirming duplication of** ***Zasp52*** **and** ***TTLL2*** **using** ***de novo*** **assemblies of** ***F. aquilonia*** **and** ***F. polyctena***

Step 1: Run code in the following page to identify supergene genotypes of the PacBio data:

Code for determining the supergene genotype for PacBio data 

Step 2: Run code in the following page to produce de novo genome assemblies and do an orthology analysis of the “minimal-P” region:

Code to produce de novo genome assemblies and orthology analyses

**Mutation load**

Step 1: Run code in the following page to quantify mutation load for each worker:

Code for mutation load analyses

**Alignment, variant calling, and expression levels of RNA-seq data**

Step 1: Run code in the following page to align RNAseq data to reference genomes, do variant calling and expression level analyses:

Code for RNA-seq data analyses
