## Supplementary Code for "The loss of a supergene in obligately polygynous *Formica* wood ant species": Code for CDS SNPs dc2ddc7c1d5345bab9d6635c4ebc8696.html

### Code for CDS SNPs

```
bcftools query -f '[%POS\t%SAMPLE\t%GT\t%DP\n]' intermediate/joint_vcfs/Scaff03_final.woodAnt.hetSites.CDS.hom10.vcf.gz | awk -v OFS="\t" '{print "Scaffold03",$1-1,$1,$2,$3,$4}' | bedtools intersect -a stdin -b ../data/external_raw/genome/Faqxpol_genome_annotation_v1.gff3 -wa -wb | grep mRNA | grep "\.t1" | cut -f 1,2,4,5,6,10,11,15 | tr ' ' \\t > scratch/Scaff03_final.woodAnt.refP.CDS.hom10.genotypeCount.perSample.position.DP.tsv
```

Rcode

```
library(ggplot2)
library(plyr)
library(dplyr)
library(doBy)
library(reshape2)
library(psych)
library(ggsci)
library(data.table)
library(ggpubr)
library(tidyr)
library(gplots)
library("scales")
library(tibble)
library(ggforce)
library(ggrepel)
library(cowplot)

setwd("/scratch/project_2004676/hanna_sigeman/ants/wood-ant-supergene")

text_size_colour = list(theme_bw(base_family="Helvetica", base_size = 12) + 
                          theme(axis.text.x= element_text(colour="black", size=12)) +
                          theme(axis.text.y= element_text(colour="black", size=12)) +
                          theme(axis.title.x = element_text(colour="black",size=14)) + 
                          theme(axis.title.y = element_text(colour="black",size=14)) + 
                          theme(axis.ticks = element_line(colour = "black", size = 0.5)) +
                          theme(panel.border = element_rect(colour = "black", fill=NA)) +
                          theme(plot.title=element_text(family="Courier", size=14, colour="black", hjust = 0.5)) +
                          theme(panel.grid.major = element_line(size = 0.15, linetype = 'solid',
                                                                colour = "lightgrey"), 
                                panel.grid.minor = element_line(size = 0.15, linetype = 'solid',
                                                                colour = "lightgrey")))

theme_title <- function(...) {
  theme_gray(base_family = "Helvetica") + 
    theme(plot.title = element_text(face = "bold"))
}
title_theme <- calc_element("plot.title", theme_title())

theme_description <- function(...) {
  theme_gray(base_family = "Helvetica") + 
    theme(plot.title = element_text(face = "plain",size = 12, hjust = 0))
}
theme_description <- calc_element("plot.title", theme_description())

MyColour <- c("#0073C2FF", "#EFC000FF", "#868686FF", "#CD534CFF", "#7AA6DCFF", "#003C67FF", "#8F7700FF")
names(MyColour) <- c("F_aquilonia", "F_lugubris", "F_paralugubris", "F_polyctena", "F_pratensis", "F_rufa", "F_truncorum")
MyShapes <- c(21, 22, 23, 24)
names(MyShapes) <- c("MM", "MP", "M", "P")

mypal <- pal_jco()(9)
mypal

show_col(mypal)

data <- read.delim("scratch/Scaff03_final.woodAnt.refP.CDS.hom10.genotypeCount.perSample.position.DP.tsv", sep="\t", header = F)
gencov.data <- read.table("results/CDS_coverage/all_samples.Scaffold03.bamstat04.depth", header = T) # Per-exon genome c
samples <- read.delim("config/samples_supergene_haplotype_species.tsv", header = F)
samples <- samples %>% rename(sample = V1, species = V2, supergene = V3)

samples$gyny <- "SP"
samples$gyny[which(c(samples$species =="F_aquilonia" | samples$species =="F_paralugubris" | samples$species =="F_polyctena" ))] <- "OP"

gencov.data$gene <- gsub("\\..*","",gencov.data$CDS)
gencov.data.small <- gencov.data %>% select(gene, sample, avgcov, mediancov, percentcovered)


data$V8 <- gsub("ID=", "", data$V8)
data$V8 <- gsub(";.*", "", data$V8)
data$V8 <- gsub("\\.t1", "", data$V8)

data <- data %>% rename(chr = V1, SNP_pos = V2, sample = V3, genotype = V4, depth = V5, start = V6, end = V7, gene = V8)

data_count_SNPs <- data %>% group_by(gene, sample) %>% summarize(nrSNPs = n())

mean_values <- data_count_SNPs %>% group_by(gene) %>% summarize(mean_nrSNPs = mean(nrSNPs, na.rm = TRUE))


# Count the occurrences of each unique combination of V1, V3, V4, V5, and their corresponding V2 values
grouped <- data %>% group_by(sample, genotype, start, end, gene) %>% summarise(count = n()) %>% ungroup()

# Calculate the proportion for each combination
grouped <- grouped %>% group_by(sample, start, end, gene) %>% mutate(sum = sum(count))

# Display the result
print(grouped)

grouped_wide <- grouped %>% pivot_wider(id_cols = c(sample, start, end, gene, sum),
                                        names_from = genotype, values_from = count, values_fill = 0)

grouped_sp <- merge(grouped_wide, samples, by.x = "sample", by.y = "sample")

grouped_sp$P_sites <- grouped_sp$`0/1` + grouped_sp$`1/1`

grouped_sp_avg <- grouped_sp %>% group_by(species, gyny, gene, start, supergene, sum) %>% summarize(mean_value = mean(P_sites), se = sd(P_sites) / sqrt(n()))

grouped_sp_avg_MM <- subset(grouped_sp_avg, c(grouped_sp_avg$supergene == "MM" |grouped_sp_avg$supergene == "M"))
grouped_sp_avg_MM$gyny_f = factor(grouped_sp_avg_MM$gyny, levels=c('SP','OP'))

grouped_sp_avg_MM_shuffled <- grouped_sp_avg_MM[sample(nrow(grouped_sp_avg_MM)), ]

pdf("results/plots/Pspecific_SNPs_gene_SP_OP.pdf", height = 6, width = 12)
ggplot(grouped_sp_avg_MM_shuffled, aes(x = start/1000000, y = mean_value, shape = supergene, fill = species, label = gene)) + 
  annotate("rect", xmin = 1.9, xmax = 11.6, ymin = -Inf, ymax = Inf, alpha = 0.5, fill = "lightgray") +
  geom_errorbar(aes(ymin=mean_value-se, ymax=mean_value+se), width=0.1) +
  geom_point(aes(size = sum)) +
  scale_shape_manual(values = MyShapes) +
  scale_fill_manual(values = MyColour) + 
  xlab("Chromosome 3 (Mb)") +  ylab(expression(paste("P-specific SNPs in MM (M) genotypes \n (average nr per species)"))) +
  text_size_colour + facet_wrap(~gyny_f, ncol = 1) + theme(plot.margin = margin(1, 1, 1, 1, "cm"))
dev.off()


grouped_sp_avg$gyny_f = factor(grouped_sp_avg$gyny, levels=c('SP','OP'))


grouped_sp_avg_MM <- subset(grouped_sp_avg_MM, grouped_sp_avg_MM$supergene == "MM")

# T-tests 
u_genes <- unique(grouped_sp_avg_MM$gene)
length(u_genes)
p_vals <- data.frame(gene = u_genes, p = rep(0, length(u_genes)))
for(i in 1:length(u_genes)){
  mod <- lm(mean_value ~ gyny, data=grouped_sp_avg_MM[which(grouped_sp_avg_MM$gene == u_genes[i]),])
  p_vals$p[i] <- summary(mod)$coefficients[,4][2] 
  
}
hist(p_vals$p)
p_vals$gene[which(p_vals$p < 0.1)]


wide_data <- pivot_wider(grouped_sp_avg_MM, 
                         id_cols = c("gene"),
                         names_from = "species",
                         values_from = c("mean_value"))


wide_data <- wide_data %>%
  column_to_rownames(var = "gene")


#wide_data = log2(wide_data[,1:7])

wide_data <- wide_data %>%
  select("F_aquilonia", "F_paralugubris", "F_polyctena", "F_lugubris", "F_pratensis", "F_rufa")  # Specify the column name


#calculate the mean of each gene per control group
control = apply(wide_data[,4:6], 1, mean)

#calcuate the mean of each gene per test group
test = apply(wide_data[, 1:3], 1, mean) 


#confirming that we have a vector of numbers
class(control) 


foldchange <- control - test 

foldchange_df <- data.frame(
  gene = names(foldchange),
  fold_change = as.numeric(as.character(foldchange))
)

hist(foldchange, xlab = "log2 Fold Change (Control vs Test)")

p_log2fold <- merge(p_vals, foldchange_df, by = "gene")
p_log2fold <- merge(p_log2fold, mean_values, by = "gene")


wide_data$gene <- rownames(wide_data)
p_log2fold <- merge(p_log2fold, wide_data, by = "gene")

p_log2fold <- p_log2fold %>%
  mutate(fold_change = ifelse(fold_change == Inf, 0, fold_change))

p_log2fold$colour <- "1"
p_log2fold$colour[which(p_log2fold$p < 0.05)] <- "2"

write.table(p_log2fold, "results/tables/p_log2fold.tsv", quote=FALSE, sep="\t", row.names = F, col.names = T, na = "NA")

p_log2fold$padj <- p.adjust(p_log2fold$p, method = "bonferroni")


ggplot(data = p_log2fold, aes(x = fold_change, y = mean_nrSNPs, colour = colour)) + geom_point()

ggplot(data = p_log2fold, aes(x = fold_change, y =  -1*log10(p), colour = p)) + geom_point()

ggplot(data = p_log2fold, aes(x = fold_change, y =  p, fill = p)) + geom_point(pch = 21, size = 5) + scale_y_reverse() + text_size_colour

p_log2fold = mutate(p_log2fold, sig=ifelse(p_log2fold$p<0.05, "p<0.05", "Not Sig"))


p2 <- ggplot(data = p_log2fold, aes(x = fold_change, y =  p, fill = sig)) + 
  geom_hline(yintercept = 0.05, linetype = "dashed") + 
  scale_fill_manual(values=c("black", "red")) + 
  geom_point(pch = 21, size = 3) + scale_y_reverse() + text_size_colour +
  geom_text_repel(data=filter(p_log2fold, p<0.05), aes(label=gene))

p1 <- ggplot(grouped_sp_avg_MM_shuffled, aes(x = start/1000000, y = mean_value, shape = supergene, fill = species, label = gene)) + 
  annotate("rect", xmin = 1.9, xmax = 11.6, ymin = -Inf, ymax = Inf, alpha = 0.5, fill = "lightgray") +
  # annotate("rect", xmin = 9.805202, xmax = 9.870675, ymin = -Inf, ymax = Inf, alpha = 0.5, fill = "lightgray") +
  #annotate("rect", xmin = 9.805202, xmax = 9.870675, ymin = 33, ymax = 36, fill = "black") +
  geom_errorbar(aes(ymin=mean_value-se, ymax=mean_value+se), width=0.1) +
  geom_point(aes(size = sum)) +
  scale_shape_manual(values = MyShapes) +
  scale_fill_manual(values = MyColour) + 
  xlab("Chromosome 3 (Mb)") +  ylab(expression(paste("P-specific SNPs"))) +
  text_size_colour + facet_wrap(~gyny_f, ncol = 1) + theme(plot.margin = margin(1, 1, 1, 1, "cm")) 


pdf("results/plots/Pspecific_SNPs_gene_SP_OP_stat_test.pdf", height = 6, width = 12)
plot_grid(p1 + theme(legend.position = 'none'), p2 + theme(legend.position = 'none'), align = "h", axis = "bt", rel_widths = c(3, 1.5))
dev.off()

summary(lm(mean_value ~ gyny, data=grouped_sp_avg_MM[which(grouped_sp_avg_MM$gene == "jg3505"),]))
summary(lm(mean_value ~ gyny, data=grouped_sp_avg_MM[which(grouped_sp_avg_MM$gene == "jg3507"),]))


#### PLOT GENE ANNOTATIONS


library("GenomicFeatures")
library("Gviz")
library(BiocManager)
library(karyoploteR)
library(ape)
library(rtracklayer)


options(ucscChromosomeNames=FALSE)

#gff.file <- import.gff("intermediate/manual_curation/Faqxpol_genome_annotation_v1.ZaspEdit.Scaffold03.cleaned.gff3")

gff.file <- import.gff("../intermediate/manual_curation/Faqxpol_genome_annotation_v1.minP.start.stop.gff3")


gtrack <- GenomeAxisTrack()

atrack <- AnnotationTrack(F.genome, name="Scaffold03")
grtrack <- GeneRegionTrack(gff.file, genome = "F.genome", parent = gff.file$Parent, transcript = gff.file$ID, showID=T,name = "Gene Model")
#plotTracks(list(gtrack, atrack, grtrack), from = 9805202, to = 9875675, extend.right = 1000, extend.left = 1000)


gff.file.df <- as.data.frame(gff.file)
gff.file.df.cds <- subset(gff.file.df, gff.file.df$type=="CDS")
aTrack.groups <- AnnotationTrack(start = gff.file.df.cds$start, 
                                 width = gff.file.df.cds$end - gff.file.df.cds$start,
                                 chromosome = "Scaffold03",
                                 strand = gff.file.df.cds$strand,
                                 group = gff.file.df.cds$Parent, 
                                 genome = "F.genome", name = "Gene Annotations",
                                 stacking = "squish")
feature(aTrack.groups) <- ifelse(strand(aTrack.groups)=="+", "plus", "minus")

pdf("results/plots/minimal_P_gene_annotations_FULL.pdf", width = 12, height = 6)
plotTracks(list(gtrack, aTrack.groups), groupAnnotation = "group", 
           plus="darkblue", minus="darkblue",
           from = 9805202, to = 9870675, extend.right = 1000, extend.left = 10000)
dev.off()


library(stringr)
gff.file.df.cds.curated <- gff.file.df.cds%>%filter(str_detect(Parent,"Zasp52_M|Zasp52_P|jg3507.t1|jg3509.t1|jg3510.t1"))
aTrack.groups2 <- AnnotationTrack(start = gff.file.df.cds.curated$start, 
                                  width = gff.file.df.cds.curated$end - gff.file.df.cds.curated$start,
                                  chromosome = "Scaffold03",
                                  strand = gff.file.df.cds.curated$strand,
                                  group = gff.file.df.cds.curated$Parent, 
                                  genome = "F.genome", name = "Gene Annotations",
                                  stacking = "squish")
feature(aTrack.groups2) <- ifelse(strand(aTrack.groups2)=="+", "plus", "minus")


pdf("results/plots/minimal_P_gene_annotations_CURATED.pdf", width = 12, height = 3)
plotTracks(list(gtrack, aTrack.groups2), groupAnnotation = "group", 
           plus="darkblue", minus="darkblue", shape = "arrow",
           from = 9805202, to = 9870675, extend.right = 1000, extend.left = 10000)
dev.off()


data(twoGroups)
dTrack <- DataTrack(twoGroups, name = "uniform")
plotTracks(dTrack)

# Plot genotype hetmaps for the Zasp52 and TTLL2 genes

MyGenoColours <- c("gray", "yellow", "orange", "red")
names(MyGenoColours) <- c("./.", "0/0", "0/1", "1/1")


h_data <- data
h_data$genotype_ID <- 0
h_data$genotype_ID[which(h_data$genotype == "0/1")] <- 1
h_data$genotype_ID[which(h_data$genotype == "1/1")] <- 2
h_data <- merge(h_data, samples, by.x = "sample", by.y = "sample")

# Convert species and supergene to factors with desired order
h_data$species <- factor(h_data$species, levels = c("F_aquilonia", "F_paralugubris", "F_polyctena", 
                                                    "F_lugubris", "F_pratensis", "F_rufa", "F_truncorum"))
h_data$supergene <- factor(h_data$supergene, levels = c("MM", "M", "MP", "PP"))

# Reorder samples based on species and then supergene
h_data$sample <- factor(h_data$sample, 
                        levels = unique(h_data[order(h_data$species, h_data$supergene), "sample"]))


ggplot(h_data, aes(depth, colour = species)) + 
  geom_density() 


p_heatmap_settings <- list(theme(strip.background = element_blank(), strip.text = element_blank()) +
                             theme(axis.title.x=element_blank(),
                                   axis.text.x=element_blank(),
                                   axis.ticks.x=element_blank()) + 
                             theme(axis.title.y=element_blank(),
                                   axis.text.y=element_blank(),
                                   axis.ticks.y=element_blank()))


##Zasp52
heatmap_jg3505 <- subset(h_data, h_data$gene=="jg3505")


h_supergene <- ggplot(heatmap_jg3505, aes(sample, 1)) + 
  geom_tile(aes(fill = supergene), color = "black") +
  ggforce::facet_row(vars(gyny), scales = 'free', space = 'free') +
  scale_fill_manual(values = MyShapes) +
  theme(axis.title.x=element_blank(),
        axis.text.x=element_blank(),
        axis.ticks.x=element_blank()) + 
  theme(axis.title.y=element_blank(),
        axis.text.y=element_blank(),
        axis.ticks.y=element_blank()) 

h_jg3505 <- ggplot(heatmap_jg3505, aes(sample, as.factor(SNP_pos))) + 
  geom_tile(aes(fill = as.factor(genotype)), color = "black") +
  scale_fill_manual(values = MyGenoColours) + 
  ggforce::facet_row(vars(gyny), scales = 'free', space = 'free') +
  p_heatmap_settings


#ggarrange(h_supergene, h_jg3505, h_jg3505_depth, h_names, ncol = 1, heights = c(0.2,1, 1, 0.5), align = "v")


##TTLL2
heatmap_jg3507 <- subset(h_data, h_data$gene=="jg3507")

h_jg3507 <- ggplot(heatmap_jg3507, aes(sample, as.factor(SNP_pos))) + 
  geom_tile(aes(fill = as.factor(genotype)), color = "black") +
  scale_fill_manual(values = MyGenoColours) + 
  ggforce::facet_row(vars(gyny), scales = 'free', space = 'free') +
  p_heatmap_settings

h_names <- ggplot(heatmap_jg3507, aes(sample, 1)) + 
  geom_tile(aes(fill = species), color = "black") +
  scale_fill_manual(values = MyColour) + 
  ggforce::facet_row(vars(gyny), scales = 'free', space = 'free') +
  theme(strip.background = element_blank(), strip.text = element_blank()) +
  theme(axis.text.x = element_text(angle = 90, vjust = 0.5, hjust=1)) +
  theme(axis.title.y=element_blank(),
        axis.text.y=element_blank(),
        axis.ticks.y=element_blank())

pdf("results/plots/Zasp52_TTLL2_heatmap_genotype.pdf", width = 12, height = 7)
ggarrange(h_supergene, h_jg3505, h_jg3507, h_names, ncol = 1, heights = c(0.2,1, 0.35, 0.5), align = "v")
dev.off()

h_jg3505_depth <- ggplot(heatmap_jg3505, aes(sample, as.factor(SNP_pos))) + 
  geom_tile(aes(fill = depth), color = "black") +
  scale_fill_gradient(low="lightblue", high="darkblue") +
  ggforce::facet_row(vars(gyny), scales = 'free', space = 'free') +
  p_heatmap_settings

h_jg3507_depth <- ggplot(heatmap_jg3507, aes(sample, as.factor(SNP_pos))) + 
  geom_tile(aes(fill = depth), color = "black") +
  scale_fill_gradient(low="lightblue", high="darkblue") +
  ggforce::facet_row(vars(gyny), scales = 'free', space = 'free') +
  p_heatmap_settings

pdf("results/plots/Zasp52_heatmap_genotype_depth.pdf", width = 18, height = 12)
ggarrange(h_supergene, h_jg3505, h_jg3505_depth, h_names, ncol = 1, heights = c(0.3,1, 1, 0.5), align = "v")
dev.off()

pdf("results/plots/TTLL2_heatmap_genotype_depth.pdf", width = 18, height = 12)
ggarrange(h_supergene, h_jg3507, h_jg3507_depth, h_names, ncol = 1, heights = c(0.3,1, 1, 0.5), align = "v")
dev.off()

depth_minP <- subset(h_data, c(h_data$start>9782000 & h_data$end<10410000))

p_minP_geno <- ggplot(depth_minP, aes(sample, as.factor(SNP_pos))) + 
  geom_tile(aes(fill = as.factor(genotype)), color = "black") +
  scale_fill_manual(values = MyGenoColours) + 
  ylab(expression(paste("Chromosome 3  (Mb)"))) +
  ggforce::facet_row(vars(gyny), scales = 'free', space = 'free') +
  p_heatmap_settings


p_minP_depth <- ggplot(depth_minP, aes(sample, as.factor(SNP_pos))) + 
  geom_tile(aes(fill = depth), color = "black") +
  scale_fill_gradient(low="lightblue", high="darkblue") +
  ylab(expression(paste("Chromosome 3 (Mb)"))) +
  ggforce::facet_row(vars(gyny), scales = 'free', space = 'free') +
  p_heatmap_settings

pdf("results/plots/minP_heatmap_genotype_depth.pdf", width = 25, height = 28)
ggarrange(h_supergene, p_minP_geno, p_minP_depth, h_names, ncol = 1, heights = c(0.15, 1, 1, 0.15), align = "v", labels = "AUTO")
dev.off()


# Other genes:
## Knockout 
p_knockout <- ggplot(subset(h_data, h_data$gene=="jg3690"), aes(sample, as.factor(SNP_pos))) + 
  geom_tile(aes(fill = as.factor(genotype)), color = "black") +
  scale_fill_manual(values = MyGenoColours) + 
  ggforce::facet_row(vars(gyny), scales = 'free', space = 'free') +
  p_heatmap_settings

## Single-minded
p_singleminded <- ggplot(subset(h_data, h_data$gene=="jg2509"), aes(sample, as.factor(SNP_pos))) + 
  geom_tile(aes(fill = as.factor(genotype)), color = "black") +
  scale_fill_manual(values = MyGenoColours) + 
  ggforce::facet_row(vars(gyny), scales = 'free', space = 'free') +
  p_heatmap_settings

## ZFP148
p_ZFP148 <- ggplot(subset(h_data, h_data$gene=="jg3646"), aes(sample, as.factor(SNP_pos))) + 
  geom_tile(aes(fill = as.factor(genotype)), color = "black") +
  scale_fill_manual(values = MyGenoColours) + 
  ggforce::facet_row(vars(gyny), scales = 'free', space = 'free') +
  p_heatmap_settings

## FMRFaR no SNPs
#p_FMRFaR <- ggplot(subset(h_data, h_data$gene=="jg3766"), aes(sample, as.factor(SNP_pos))) + 
#  geom_tile(aes(fill = as.factor(genotype)), color = "black") +
#  scale_fill_manual(values = MyGenoColours) + 
#  ggforce::facet_row(vars(gyny), scales = 'free', space = 'free') +
#  p_heatmap_settings

## AmGR10
p_AmGR10 <- ggplot(subset(h_data, h_data$gene=="jg3649"), aes(sample, as.factor(SNP_pos))) + 
  geom_tile(aes(fill = as.factor(genotype)), color = "black") +
  scale_fill_manual(values = MyGenoColours) + 
  ggforce::facet_row(vars(gyny), scales = 'free', space = 'free') +
  p_heatmap_settings

pdf("results/plots/duplicated_genes_Purcell.pdf", width = 18, height = 22)
ggarrange(h_supergene, p_knockout, p_singleminded, p_ZFP148, p_AmGR10, h_names, ncol = 1, heights = c(0.35, 1, 1, 1, 1, 0.35), align = "v", labels = "AUTO")
dev.off()
```
