## Supplementary Code for "The loss of a supergene in obligately polygynous *Formica* wood ant species": Code for determining the supergene genotype for Pa 3f46460f903b422495f3896b0e6f23b6.html

Code for determining the supergene genotype for PacBio data  

### Code for determining the supergene genotype for PacBio data

PacBio reads are aligned to the *F. aquilonia x polyctena* reference genome with the snakemake file

```
conda create -c bioconda -n longshot longshot

# Call variants with longshot

#!/bin/bash
#SBATCH --job-name=longshot
#SBATCH --output=longshot.out.log
#SBATCH --error=longshot.err.log
#SBATCH --account=project_2004676
#SBATCH --partition=small
#SBATCH --time=1-00:00:00
#SBATCH --ntasks=1
#SBATCH --mem-per-cpu=6000
#SBATCH --cpus-per-task=10

source activate longshot

longshot --bam intermediate/dedup/Faqxpol_pacbio.sorted.bam --ref ../data/external_raw/genome/Formica_hybrid_v1_wFhyb_Sapis.fa --out intermediate/variant_calling/Faqxpol_pacbio.Scaffold03.longshot.vcf -r Scaffold03

longshot --bam intermediate/dedup/Faql_pacbio.sorted.bam --ref ../data/external_raw/genome/Formica_hybrid_v1_wFhyb_Sapis.fa --out intermediate/variant_calling/Faql_pacbio.Scaffold03.longshot.vcf -r Scaffold03

longshot --bam intermediate/dedup/Fpol_pacbio.sorted.bam --ref ../data/external_raw/genome/Formica_hybrid_v1_wFhyb_Sapis.fa --out intermediate/variant_calling/Fpol_pacbio.Scaffold03.longshot.vcf -r Scaffold03


vcftools --vcf intermediate/variant_calling/Faqxpol_pacbio.Scaffold03.longshot.vcf --positions intermediate/joint_vcfs/Scaff03_final.MPind.hetSites.pos --recode --stdout > scratch/Faqxpol_pacbio_MPind.hetSites.vcf

bgzip -c scratch/Faqxpol_pacbio_MPind.hetSites.vcf > scratch/Faqxpol_pacbio_MPind.hetSites.vcf.gz
tabix -p vcf scratch/Faqxpol_pacbio_MPind.hetSites.vcf.gz

bcftools query -f '[%SAMPLE\t%GT\n]' scratch/Faqxpol_pacbio_MPind.hetSites.vcf.gz | sed 's/ /\t/g' | wc -l # 499 SNPs


vcftools --vcf intermediate/variant_calling/Faql_pacbio.Scaffold03.longshot.vcf --positions intermediate/joint_vcfs/Scaff03_final.MPind.hetSites.pos --recode --stdout > scratch/Faql_pacbio_MPind.hetSites.vcf

bgzip -c scratch/Faql_pacbio_MPind.hetSites.vcf > scratch/Faql_pacbio_MPind.hetSites.vcf.gz
tabix -p vcf scratch/Faql_pacbio_MPind.hetSites.vcf.gz

bcftools query -f '[%SAMPLE\t%GT\n]' scratch/Faql_pacbio_MPind.hetSites.vcf.gz | sed 's/ /\t/g' | wc -l # 687 SNPs


vcftools --vcf intermediate/variant_calling/Fpol_pacbio.Scaffold03.longshot.vcf --positions intermediate/joint_vcfs/Scaff03_final.MPind.hetSites.pos --recode --stdout > scratch/Fpol_pacbio_MPind.hetSites.vcf

bgzip -c scratch/Fpol_pacbio_MPind.hetSites.vcf > scratch/Fpol_pacbio_MPind.hetSites.vcf.gz
tabix -p vcf scratch/Fpol_pacbio_MPind.hetSites.vcf.gz

bcftools query -f '[%SAMPLE\t%GT\n]' scratch/Fpol_pacbio_MPind.hetSites.vcf.gz | sed 's/ /\t/g' | wc -l # 958 SNPs
```
