## Supplementary Code for "The loss of a supergene in obligately polygynous *Formica* wood ant species": Code for mutation load analyses 678fd29e42dd44dd8c53af1917365f18.html

### Code for mutation load analyses

Build snpEff database

```
sinteractive --account project_2005765 --cores 7 --time 6:00:00 --mem 50000 --tmp 100

conda activate snpsift
module load biojava
module load openjdk/17.0.3_7

mkdir -p code/snpEff/data/Faqxpol_scaffolds_v1
# snpEff database
cp ../data/external_raw/genome/Faqxpol_scaffolds_v1.fa code/snpEff/data/Faqxpol_scaffolds_v1/sequences.fa

cp ../data/external_raw/genome/Faqxpol_genome_annotation_v1.gff3 code/snpEff/data/Faqxpol_scaffolds_v1/genes.gff

gff=../data/external_raw/genome/Faqxpol_genome_annotation_v1.gff3
ref=../data/external_raw/genome/Faqxpol_scaffolds_v1.fa
agat="apptainer exec -B /scratch:/scratch /projappl/project_2004676/hanna_sigeman/agat_1.0.0--pl5321hdfd78af_0.sif"

${agat} agat_convert_sp_gff2gtf.pl -gff $gff --gtf_version 2.2 --out ../data/external_raw/genome/Faqxpol_genome_annotation_v1.agat.gtf

cp ../data/external_raw/genome/Faqxpol_genome_annotation_v1.agat.gtf code/snpEff/data/Faqxpol_scaffolds_v1/genes.gtf

gffread ../data/external_raw/genome/Faqxpol_genome_annotation_v1.agat.gtf -g ../data/external_raw/genome/Faqxpol_scaffolds_v1.fa -V -y code/snpEff/data/Faqxpol_scaffolds_v1/protein.fa
gffread ../data/external_raw/genome/Faqxpol_genome_annotation_v1.agat.gtf -g ../data/external_raw/genome/Faqxpol_scaffolds_v1.fa -V -x code/snpEff/data/Faqxpol_scaffolds_v1/cds.fa


echo "Faqxpol_scaffolds_v1.genome: Faqxpol_scaffolds_v1" >> code/snpEff/snpEff.config

echo "Faqxpol_scaffolds_v1.genome: Faqxpol_scaffolds_v1" >> /projappl/project_2004676/hanna_sigeman/envs/snpsift/share/snpeff-5.2-0/snpEff.config


snpEff -Xmx8g build -gtf22 -v Faqxpol_scaffolds_v1 -dataDir /scratch/project_2004676/hanna_sigeman/ants/wood-ant-supergene/code/snpEff/data -nodownload
```

Run snpSift

```
outdir=intermediate/mutation_load/allSamples
vcf=intermediate/joint_vcfs/Scaff03_final.vcf.gz
mkdir -p $outdir

cat config/samples_supergene_haplotype_species.tsv | cut -f 1 | while IFS= read -r string; do echo "$string" > "${outdir}/$string"; done

cat config/samples_supergene_haplotype_species.tsv | cut -f 1 | while read sample ; do vcftools --gzvcf $vcf --keep ${outdir}/${sample} --from-bp 1900000 --to-bp 11600000 --chr Scaffold03 --mac 1 --recode --stdout > ${outdir}/${sample}.vcf ; done

cat config/samples_supergene_haplotype_species.tsv | cut -f 1 | while read sample ; do snpEff ann -v Faqxpol_scaffolds_v1 ${outdir}/${sample}.vcf -dataDir /scratch/project_2004676/hanna_sigeman/ants/wood-ant-supergene/code/snpEff/data -csvStats ${outdir}/${sample}.snpEff.report > ${outdir}/${sample}.snpEff.vcf ; done

cat config/samples_supergene_haplotype_species.tsv | cut -f 1 | while read sample ; do cat ${outdir}/${sample}.snpEff.report.genes.txt | grep ".t1" | cut -f 1,5-7 | awk '{print "'"$sample"'",$0}' | sed 's/ /\t/g' ; done > results/allSamples_mutationLoad_snpEff_HIGHMODERATELOWMODIFIER_t1.tsv
```

Rcode for plotting and statistics

```
library(ggplot2)
library(plyr)
library(dplyr)
library(doBy)
library(reshape2)
library(psych)
library(ggsci)
library(data.table)
library(ggpubr)
library(tidyr)
library(gplots)
library("scales")
library(tibble)
library(ggforce)
library(stringr)
library(ggnewscale)
setwd("~/work/wood-ant-supergene/results/")

options(scipen = 999)
text_size_colour = list(theme_bw(base_family="Helvetica", base_size = 12) + 
                          theme(axis.text.x= element_text(colour="black", size=12)) +
                          theme(axis.text.y= element_text(colour="black", size=12)) +
                          theme(axis.title.x = element_text(colour="black",size=14)) + 
                          theme(axis.title.y = element_text(colour="black",size=14)) + 
                          theme(axis.ticks = element_line(colour = "black", size = 0.5)) +
                          theme(panel.border = element_rect(colour = "black", fill=NA)) +
                          theme(plot.title=element_text(family="Courier", size=14, colour="black", hjust = 0.5)) +
                          theme(panel.grid.major = element_line(size = 0.15, linetype = 'solid',
                                                                colour = "lightgrey"), 
                                panel.grid.minor = element_line(size = 0.15, linetype = 'solid',
                                                                colour = "lightgrey")))

theme_title <- function(...) {
  theme_gray(base_family = "Helvetica") + 
    theme(plot.title = element_text(face = "bold"))
}
title_theme <- calc_element("plot.title", theme_title())

theme_description <- function(...) {
  theme_gray(base_family = "Helvetica") + 
    theme(plot.title = element_text(face = "plain",size = 12, hjust = 0))
}
theme_description <- calc_element("plot.title", theme_description())

samples <- read.delim("../config/samples_supergene_haplotype_species.tsv", header = F)
samples <- samples %>% rename(Sample = V1, Species = V2, Supergene = V3)
genes <- read.delim("Faqxpol_scaffolds_v1_genes_Scaffold03_snpEff.small.bed", sep="\t", header = F)
genes <- genes %>% select(V1, V2, V3, V4) %>% rename(c(Chr = V1, Start = V2, End = V3, Gene = V4))


#snpEff <- read.delim("Ftruncroum_snpEffgenes.tsv", sep="\t", header = F)
snpEff <- read.delim("allSpecies.genes.1ind.txt", sep="\t", header = F)

dnds <- read.delim("allSamples_mutationLoad_snpEff_missenseSilent.tsv", sep="\t", header = F)
HIGHMODERATELOW <- read.delim("allSamples_mutationLoad_snpEff_HIGHMODERATELOWMODIFIER_t1.tsv", sep="\t", header = F)
HIGHMODERATELOW <- HIGHMODERATELOW %>% rename(c(Sample = V1, Gene = V2, HIGH = V3, LOW = V4, MODERATE = V5))


HIGHMODERATELOW <- merge(HIGHMODERATELOW, samples, by = "Sample")

HIGHMODERATELOW_MM_MP <- subset(HIGHMODERATELOW, c(HIGHMODERATELOW$Supergene=="MM" | HIGHMODERATELOW$Supergene=="MP"))

HIGHMODERATELOW_MM_MP$TotalCount <- HIGHMODERATELOW_MM_MP$HIGH + HIGHMODERATELOW_MM_MP$MODERATE + HIGHMODERATELOW_MM_MP$LOW

HIGHMODERATELOW_MM_MP_sum <- HIGHMODERATELOW_MM_MP %>%
  group_by(Sample) %>%
  summarize(
    HIGH_sum = sum(HIGH),
    LOW_sum = sum(LOW),
    MODERATE_sum = sum(MODERATE),
    TotalCount_sum = sum(TotalCount)
  )

HIGHMODERATELOW_MM_MP_sum <- merge(HIGHMODERATELOW_MM_MP_sum, samples, by = "Sample")
HIGHMODERATELOW_MM_MP_sum$Gyny <- "SP"
HIGHMODERATELOW_MM_MP_sum$Gyny[which(HIGHMODERATELOW_MM_MP_sum$Species=="F_aquilonia")] <- "OP"
HIGHMODERATELOW_MM_MP_sum$Gyny[which(HIGHMODERATELOW_MM_MP_sum$Species=="F_polyctena")] <- "OP"
HIGHMODERATELOW_MM_MP_sum$Gyny[which(HIGHMODERATELOW_MM_MP_sum$Species=="F_paralugubris")] <- "OP"

p1 <- ggplot(HIGHMODERATELOW_MM_MP_sum, aes(x = Supergene, y = HIGH_sum, fill = Supergene)) + 
  geom_boxplot() + stat_compare_means(method = "wilcox.test", vjust = 15, bracket.size = 0) +
  text_size_colour + scale_fill_startrek() + ylab("Nr. HIGH-impact SNPs") + theme(axis.title.x = element_blank()) 

p2 <- ggplot(HIGHMODERATELOW_MM_MP_sum, aes(x = Supergene, y = MODERATE_sum, fill = Supergene)) + 
  geom_boxplot() + stat_compare_means(method = "wilcox.test", vjust = 15, bracket.size = 0) +
  text_size_colour + scale_fill_startrek() + ylab("Nr. MODERATE-impact SNPs") + theme(axis.title.x = element_blank()) 

p3 <- ggplot(HIGHMODERATELOW_MM_MP_sum, aes(x = Supergene, y = LOW_sum, fill = Supergene)) + 
  geom_boxplot() + stat_compare_means(method = "wilcox.test", vjust = 15, bracket.size = 0) +
  text_size_colour + scale_fill_startrek() + ylab("Nr. LOW-impact SNPs") + theme(axis.title.x = element_blank()) 

p4 <- ggplot(HIGHMODERATELOW_MM_MP_sum, aes(x = Supergene, y = HIGH_sum/TotalCount_sum, fill = Supergene)) + 
  geom_boxplot() + stat_compare_means(method = "wilcox.test", vjust = 15, bracket.size = 0) +
  text_size_colour + scale_fill_startrek() + ylab("Prop. HIGH-impact SNPs") + theme(axis.title.x = element_blank()) 

p5 <- ggplot(HIGHMODERATELOW_MM_MP_sum, aes(x = Supergene, y = MODERATE_sum/TotalCount_sum, fill = Supergene)) + 
  geom_boxplot() + stat_compare_means(method = "wilcox.test", vjust = 15, bracket.size = 0) +
  text_size_colour + scale_fill_startrek() + ylab("Prop. MODERATE-impact SNPs") + theme(axis.title.x = element_blank()) 

p6 <- ggplot(HIGHMODERATELOW_MM_MP_sum, aes(x = Supergene, y = LOW_sum/TotalCount_sum, fill = Supergene)) + 
  geom_boxplot() + stat_compare_means(method = "wilcox.test", vjust = 15, bracket.size = 0) +
  text_size_colour + scale_fill_startrek() + ylab("Prop. LOW-impact SNPs") + theme(axis.title.x = element_blank()) 


pdf("plots/Figure5_new_boxplot.pdf", width = 12, height = 8)
ggarrange(p1, p2, p3, p4, p5, p6, ncol = 3, nrow = 2, legend = "none", common.legend = TRUE) 
dev.off()

table(HIGHMODERATELOW_MM_MP_sum$Supergene)


wilcox.test(HIGH_sum ~ Supergene, data = HIGHMODERATELOW_MM_MP_sum, paired = FALSE)
wilcox.test(MODERATE_sum ~ Supergene, data = HIGHMODERATELOW_MM_MP_sum, paired = FALSE)
wilcox.test(LOW_sum ~ Supergene, data = HIGHMODERATELOW_MM_MP_sum, paired = FALSE)
wilcox.test(HIGH_sum/TotalCount_sum ~ Supergene, data = HIGHMODERATELOW_MM_MP_sum, paired = FALSE)
wilcox.test(MODERATE_sum/TotalCount_sum ~ Supergene, data = HIGHMODERATELOW_MM_MP_sum, paired = FALSE)
wilcox.test(LOW_sum/TotalCount_sum ~ Supergene, data = HIGHMODERATELOW_MM_MP_sum, paired = FALSE)

outname <- sprintf("tables/TableS11.tsv")
write.table(HIGHMODERATELOW_MM_MP_sum, file = outname, sep = "\t", quote = FALSE, row.names = F)
```
