## Supplementary Code for "The loss of a supergene in obligately polygynous *Formica* wood ant species": Code for orthology analysis 9155d72cf1c74358a494bde6101e5af4.html

### Code for orthology analysis

Download genomes and annotations

```
mkdir intermediate/orthofinder

# Download C. floridanus
wget https://ftp.ncbi.nlm.nih.gov/genomes/all/GCF/003/227/725/GCF_003227725.1_Cflo_v7.5/GCF_003227725.1_Cflo_v7.5_protein.faa.gz
gunzip GCF_003227725.1_Cflo_v7.5_protein.faa.gz

# Download Drosophila
wget https://ftp.ncbi.nlm.nih.gov/genomes/all/GCF/000/001/215/GCF_000001215.4_Release_6_plus_ISO1_MT/GCF_000001215.4_Release_6_plus_ISO1_MT_genomic.fna.gz
wget https://ftp.ncbi.nlm.nih.gov/genomes/all/GCF/000/001/215/GCF_000001215.4_Release_6_plus_ISO1_MT/GCF_000001215.4_Release_6_plus_ISO1_MT_genomic.gff.gz
wget https://ftp.ncbi.nlm.nih.gov/genomes/all/GCF/000/001/215/GCF_000001215.4_Release_6_plus_ISO1_MT/GCF_000001215.4_Release_6_plus_ISO1_MT_protein.faa.gz 
 
# Download S. invicta
wget https://ftp.ncbi.nlm.nih.gov/genomes/all/GCF/016/802/725/GCF_016802725.1_UNIL_Sinv_3.0/GCF_016802725.1_UNIL_Sinv_3.0_protein.faa.gz
wget https://ftp.ncbi.nlm.nih.gov/genomes/all/GCF/016/802/725/GCF_016802725.1_UNIL_Sinv_3.0/GCF_016802725.1_UNIL_Sinv_3.0_protein.faa.gz
wget https://ftp.ncbi.nlm.nih.gov/genomes/all/GCF/016/802/725/GCF_016802725.1_UNIL_Sinv_3.0/GCF_016802725.1_UNIL_Sinv_3.0_genomic.fna.gz 

# Download F. exsecta
wget https://ftp.ncbi.nlm.nih.gov/genomes/all/GCF/003/651/465/GCF_003651465.1_ASM365146v1/GCF_003651465.1_ASM365146v1_protein.faa.gz 
wget https://ftp.ncbi.nlm.nih.gov/genomes/all/GCF/003/651/465/GCF_003651465.1_ASM365146v1/GCF_003651465.1_ASM365146v1_genomic.fna.gz 
wget https://ftp.ncbi.nlm.nih.gov/genomes/all/GCF/003/651/465/GCF_003651465.1_ASM365146v1/GCF_003651465.1_ASM365146v1_genomic.gff.gz 


ls | grep gz | grep GCF | while read file ; do gunzip $file ; done


mkdir intermediate/orthofinder/allSpecies
cp ../data/external_raw/genome/GCF_016802725.1_UNIL_Sinv_3.0_protein.faa intermediate/orthofinder/allSpecies/Sinv.pep.fa
cp ../data/external_raw/genome/GCF_000001215.4_Release_6_plus_ISO1_MT_protein.faa intermediate/orthofinder/allSpecies/Dmel.pep.fa
cp ../data/external_raw/genome/GCF_003227725.1_Cflo_v7.5_protein.faa intermediate/orthofinder/allSpecies/Cflo.pep.fa
cp intermediate/orthofinder/Faqxpol_Fselysi/Faqxpol.pep.fa intermediate/orthofinder/allSpecies/
cp ../data/external_raw/genome/GCF_003651465.1_ASM365146v1_protein.faa intermediate/orthofinder/allSpecies/Fexs.pep.fa
```

Orthology analysis

```
#!/bin/bash
#SBATCH --job-name=orthofinder
#SBATCH --account=project_2004676
#SBATCH --partition=small
#SBATCH --time=20:00:00
#SBATCH --ntasks=1
#SBATCH --mem-per-cpu=6000
#SBATCH --cpus-per-task=8


source activate snpsift
module load biokit
module load python-data
orthofinder -f intermediate/orthofinder/allSpecies/ -t 6 -n orthology_allSpecies
```
