## Supplementary Code for "The loss of a supergene in obligately polygynous *Formica* wood ant species": Code for phylogenetic tree (Figure 2a) 9513194fd1784310b703ed7a05b64f00.html

### Code for phylogenetic tree (Figure 2a)

Analysis according to tutorial: https://github.com/ForBioPhylogenomics/tutorials/tree/main/species\_tree\_inference\_with\_snp\_data

Install software

```
# Installation

### Download command-line PAUP: http://phylosolutions.com/paup-test/

gunzip paup4a168_osx.gz 
chmod a+x paup4a168_osx 
ls paup4a168_osx

### Download conversion script 
wget https://raw.githubusercontent.com/ForBioPhylogenomics/tutorials/main/week2_src/convert_vcf_to_nexus.rb
```

Make per-sample phylogenetic tree

```
# Create bed file from the non-supergene region on Scaffold03. Specifically, 10kb from the ends of each chromosome end and 200kb away from the supergene region:

echo -e "Scaffold03\t10000\t1696491\nScaffold03\t11913739\t12988361" > scratch/outsideSupergene.bed

# Filter and bgzip VCF file
vcftools --gzvcf intermediate/joint_vcfs/Scaff03.outgroup.deepvariant.vcf.gz --bed scratch/outsideSupergene.bed --recode --stdout > scratch/Scaff03.outgroup.deepvariant.outsideSupergene.vcf

vcftools --vcf scratch/Scaff03.outgroup.deepvariant.outsideSupergene.vcf --exclude-positions intermediate/joint_vcfs/Scaff03_final.MPind.hetSites.pos --recode --stdout > scratch/Scaff03.outgroup.deepvariant.outsideSupergene.noHetSites.vcf

bgzip -c scratch/Scaff03.outgroup.deepvariant.outsideSupergene.noHetSites.vcf > scratch/Scaff03.outgroup.deepvariant.outsideSupergene.noHetSites.vcf.gz
tabix -p vcf scratch/Scaff03.outgroup.deepvariant.outsideSupergene.noHetSites.vcf.gz -f

# Filter VCF and count SNPs
bcftools view -H scratch/Scaff03.outgroup.deepvariant.outsideSupergene.noHetSites.vcf.gz | wc -l # 37862 SNPs

vcftools --gzvcf scratch/Scaff03.outgroup.deepvariant.outsideSupergene.noHetSites.vcf.gz --min-alleles 2 --max-alleles 2 --minQ 20 --minDP 3 --max-missing 0.3 --remove-filtered-all --recode --stdout | bgzip -c > scratch/Scaff03.outgroup.deepvariant.outsideSupergene.noHetSites.filt.vcf.gz

bcftools view -H scratch/Scaff03.outgroup.deepvariant.outsideSupergene.noHetSites.filt.vcf.gz | wc -l # 33605

vcftools --gzvcf scratch/Scaff03.outgroup.deepvariant.outsideSupergene.noHetSites.vcf.gz --remove-indels --remove-filtered-all --recode --stdout | bgzip -c > scratch/Scaff03.outgroup.deepvariant.outsideSupergene.noHetSites.filt.noIndel.vcf.gz

bcftools view -H scratch/Scaff03.outgroup.deepvariant.outsideSupergene.noHetSites.filt.noIndel.vcf.gz | wc -l # 31911

vcftools --gzvcf scratch/Scaff03.outgroup.deepvariant.outsideSupergene.noHetSites.filt.noIndel.vcf.gz --thin 100 --remove-filtered-all --recode --stdout > scratch/Scaff03.outgroup.deepvariant.outsideSupergene.noHetSites.filt.noIndel.thin.vcf

grep -v "#" scratch/Scaff03.outgroup.deepvariant.outsideSupergene.noHetSites.filt.noIndel.thin.vcf | wc -l # 7805

### Convert VCF to Nexus
ruby ~/Downloads/convert_vcf_to_nexus.rb scratch/Scaff03.outgroup.deepvariant.outsideSupergene.noHetSites.filt.noIndel.thin.vcf scratch/Scaff03.outgroup.deepvariant.outsideSupergene.noHetSites.filt.noIndel.thin.nex

### First, we run the analysis on all samples to see which samples are not monophyltetic

### Run PAUP
~/Downloads/paup4a168_osx

execute scratch/Scaff03.outgroup.deepvariant.outsideSupergene.noHetSites.filt.noIndel.thin.nex

tstatus full # Check numbers of samples (to select the outgroups)

outgroup 65 66

svdQuartets # DONE!

saveTrees file='formica.tre' supportValues=nodeLabels
```

Exclude samples that are not monophyletic (samples highlighted in Figure S3)

```
# Samples to exclude 
echo "Fpolyctena_1
Fpolyctena_5
Fparalugubris_16
Flugubris_9
Flugubris_10
Flugubris_14
Flugubris_16
Flugubris_17
Flugubris_15
Flugubris_19
Faquilonia_36
Faquilonia_37
Faquilonia_38
Faquilonia_42
Faquilonia_43
Faquilonia_44" > exclude.list


vcftools --vcf scratch/Scaff03.outgroup.deepvariant.outsideSupergene.noHetSites.filt.noIndel.thin.vcf --remove exclude.list --remove-filtered-all --recode --stdout > scratch/Scaff03.outgroup.deepvariant.outsideSupergene.noHetSites.filt.noIndel.thin.rmHybr.vcf

ruby ~/Downloads/convert_vcf_to_nexus.rb scratch/Scaff03.outgroup.deepvariant.outsideSupergene.noHetSites.filt.noIndel.thin.rmHybr.vcf scratch/Scaff03.outgroup.deepvariant.outsideSupergene.noHetSites.filt.noIndel.thin.rmHybr.nex

~/Downloads/paup4a168_osx
execute scratch/Scaff03.outgroup.deepvariant.outsideSupergene.noHetSites.filt.noIndel.thin.rmHybr.nex

tstatus full

outgroup 59 60 

svdQuartets

saveTrees file='formica.rmHybr.tre' supportValues=nodeLabels
```

Make per-species phylogenetic tree

```
echo "BEGIN SETS;
  	TAXPARTITION SPECIES =
  		Faquilonia: 1-58,
  		Fexsecta: 59-60,
  		Flugubris: 61-68,
  		Fparalugubris: 69-86,
  		Fpolyctena: 87-90,
  		Fpratensis: 91-117,
  		Frufa: 118-123,
  		Ftruncorum: 124-125;
  END;" > taxpartitions.txt
  
cat scratch/Scaff03.outgroup.deepvariant.outsideSupergene.noHetSites.filt.noIndel.thin.rmHybr.nex taxpartitions.txt  > scratch/Scaff03.outgroup.deepvariant.outsideSupergene.noHetSites.filt.noIndel.thin.rmHybr.parts.nex

~/Downloads/paup4a168_osx
execute scratch/Scaff03.outgroup.deepvariant.outsideSupergene.noHetSites.filt.noIndel.thin.rmHybr.parts.nex

outgroup 59 60 
svdQuartets taxpartition=SPECIES bootstrap=standard nthreads=2
```
