## Supplementary Code for "The loss of a supergene in obligately polygynous *Formica* wood ant species": Code for RNA-seq data analyses 37af0039d4a34811947f4f2709327878.html

### Code for ***RNA-seq data analyses***

Code adapted from: https://gist.githubusercontent.com/PoisonAlien/c6c03539cf4b1ac41cf1/raw/43c0657734825c7ec2982ed016290beef155efc5/rna\_seq\_variant\_pipeline.sh

```
#!/bin/bash
#
# AUTHOR: Anand M.
# RNA-Seq variant calling pieline accoring to GATK Best practices.
# https://www.broadinstitute.org/gatk/guide/article?id=3891
#
# Call with following arguments
# bash rna_seq_variant_pipeline.sh <Input_Reads1.fq.gz> <Input_Reads2.fq.gz> <output_basename>
# 
# Assumes STAR aligner is under path
#
# GATK bundle set : one can obtain these from gatk ftp (knonw as gatk bundle)
# ftp://ftp.broadinstitute.org/bundle/2.8/hg19/
#


# sinteractive --account project_2004676 --cores 7 --time 1-00:00:00 --mem 50000 --tmp 100


#!/bin/bash
#SBATCH --job-name=STAR
#SBATCH --output=logs/mapping/STAR.txt
#SBATCH --error=logs/mapping/STAR.txt
#SBATCH --account=project_2004676
#SBATCH --partition=small
#SBATCH --time=2-00:00:00
#SBATCH --ntasks=1
#SBATCH --mem-per-cpu=6000
#SBATCH --cpus-per-task=8

module load biokit
module load gatk


# 151 read length
fwd=../data/internal_raw/fastq/Faqu_FKRN220303431-1A_H22GGDSX5_L4_1.fq.gz
rev=../data/internal_raw/fastq/Faqu_FKRN220303431-1A_H22GGDSX5_L4_2.fq.gz
bn=Faqu_FKRN220303431

#Path to reference genome and Index files.
star_ref="../data/external_raw/genome/star_index_149"
ref="../data/external_raw/genome/Faqxpol_scaffolds_v1.fa"
gtf="../data/external_raw/genome/Faqxpol_genome_annotation_v1.gtf"

opdir=intermediate/RNA_seq_GATK_pipeline/Faqxpol_scaffolds_v1/$bn

#STAR --runThreadN 6 --runMode genomeGenerate --genomeDir $star_ref --genomeFastaFiles $ref --sjdbGTFfile $gtf --sjdbOverhang 149 --genomeSAindexNbases 13

# picard CreateSequenceDictionary R=$ref O=$(echo $ref | sed 's/.fa//').dict


#Create an output directory
mkdir -p $opdir

#STAR 2 pass basic mode run
echo -e "["$(date)"]\tAligning.."
STAR --outFileNamePrefix $opdir/$bn --outSAMtype BAM Unsorted --outSAMstrandField intronMotif --outSAMattrRGline ID:$bn LB:PairedEnd PL:Illumina PU:Unknown SM:$bn --genomeDir $star_ref --runThreadN 6 --readFilesCommand zcat --readFilesIn $fwd $rev --twopassMode Basic

#sambamba sort
echo -e "["$(date)"]\tSorting.."
sambamba sort -o $opdir/$bn"_sorted.bam" -p -t 6 $opdir/$bn"Aligned.out.bam"

rm $opdir/$bn"Aligned.out.bam"

#sambamba index
echo -e "["$(date)"]\tIndexing.."
sambamba index -p -t 50 $opdir/$bn"_sorted.bam"

#picard mark duplicates
echo -e "["$(date)"]\tMarking duplicates.."
picard MarkDuplicates I=$opdir/$bn"_sorted.bam" O=$opdir/$bn"_dupMarked.bam" M=$opdir/$bn"_dup.metrics" TMP_DIR=./tmp CREATE_INDEX=true VALIDATION_STRINGENCY=SILENT 2>$opdir/$bn.MarkDuplicates.log

rm $opdir/$bn"_sorted.bam"
rm $opdir/$bn"_sorted.bam.bai"

#SplitNCigarReads
echo -e "["$(date)"]\tSpliting reads.."
gatk SplitNCigarReads -R $ref -I $opdir/$bn"_dupMarked.bam" -O $opdir/$bn"_split.bam" --tmp-dir ./tmp 2>$opdir/$bn.SplitNCigarReads.log

rm $opdir/$bn"_dupMarked.bam"
rm $opdir/$bn"_dupMarked.bai"

#Create targets for indel realignment
#echo -e "["$(date)"]\tCreating targets for indel realignment.."
#gatk -T RealignerTargetCreator -R $ref -I $opdir/$bn"_split.bam" -o $opdir/$bn".intervals" -nt 50 -known $millsIndels -known $KGIndels 2>$opdir/$bn.indel.log

#Perform indel realignment
#echo -e "["$(date)"]\tPerforming Indel Realignment.."
#gatk -T IndelRealigner -R $ref -I $opdir/$bn"_split.bam" -targetIntervals $opdir/$bn".intervals" -known $millsIndels -known $KGIndels -o $opdir/$bn"_processed.bam" 2>$opdir/$bn.indel2.log 

#rm $opdir/$bn"_split.bam"
#rm $opdir/$bn"_split.bai"

#Perform BQSR
#echo -e "["$(date)"]\tPerforming BQSR.."
#gatk -T BaseRecalibrator -I $opdir/$bn"_processed.bam" -R $ref -knownSites $KGIndels -knownSites $millsIndels -knownSites $dbSNP138 -o $opdir/$bn"_recal.table" 2>$opdir/$bn.BQSR.log

#Print recalibrated reads
#echo -e "["$(date)"]\tPrinting recalibrated reads.."
#java -d64 -jar $gatk -T PrintReads -R $ref -I $opdir/$bn"_processed.bam" -nct 50 -BQSR $opdir/$bn"_recal.table" -o $opdir/$bn"_recal.bam" 2>$opdir/$bn.BQSR2.log

#rm $opdir/$bn"_processed.bam"
#rm $opdir/$bn"_processed.bai"

#Run HaplotypeCaller
echo -e "["$(date)"]\tRunning HaplotypeCaller.."
gatk HaplotypeCaller -R $ref -I $opdir/$bn"_split.bam" -L Scaffold03 -O $opdir/$bn".Scaffold03.vcf" 2>$opdir/$bn.HaplotypeCaller.log

# Count number of fixed P-specific SNPs in Zasp52 (16)
bedtools intersect -b intermediate/joint_vcfs/Scaff03_final.woodAnt.hetSites.CDS.hom10.vcf -a intermediate/RNA_seq_GATK_pipeline/Faqxpol_scaffolds_v1/Faqu_FKRN220303431/Faqu_FKRN220303431.Scaffold03.vcf -wa -wb | cut -f 1-10 | awk '{print $1,$2-1,$2,$0}' | sed 's/ /\t/g' | bedtools intersect -a stdin -b meta/Faqxpol_genome_annotation_v1.agat.longest.isoform.CDS.bed -wa -wb | grep jg3505 | wc -l


# Allelic depth (for the manuscript)
bedtools intersect -b intermediate/joint_vcfs/Scaff03_final.woodAnt.hetSites.CDS.hom10.vcf -a intermediate/RNA_seq_GATK_pipeline/Faqxpol_scaffolds_v1/Faqu_FKRN220303431/Faqu_FKRN220303431.Scaffold03.vcf -wa -wb | cut -f 1-10 | awk '{print $1,$2-1,$2,$0}' | sed 's/ /\t/g' | bedtools intersect -a stdin -b meta/Faqxpol_genome_annotation_v1.agat.longest.isoform.CDS.bed -wa -wb | grep jg3505 | cut -f 12,13 | sed 's/:/\t/g' | cut -f 6,7 | sed 's/,/\t/g' | sort -k3,3g

bedtools intersect -b intermediate/joint_vcfs/Scaff03_final.woodAnt.hetSites.CDS.hom10.vcf -a intermediate/RNA_seq_GATK_pipeline/Faqxpol_scaffolds_v1/Faqu_FKRN220303431/Faqu_FKRN220303431.Scaffold03.vcf -wa -wb | cut -f 1-10 | awk '{print $1,$2-1,$2,$0}' | sed 's/ /\t/g' | bedtools intersect -a stdin -b meta/Faqxpol_genome_annotation_v1.agat.longest.isoform.CDS.bed -wa -wb | grep jg3507 | cut -f 12,13 | sed 's/:/\t/g' | cut -f 6,7 | sed 's/,/\t/g' | sort -k3,3g


#Filter variants
#echo -e "["$(date)"]\tFiltering Variants.."
#gatk VariantFiltration -R $ref -V $opdir/$bn".Scaffold03.vcf" -window 35 -cluster 3 -filterName FS -filter "FS > 30.0" -filterName QD -filter "QD < 2.0" -O $opdir/$bn"_filtered.vcf" 2>$opdir/$bn.VariantFilter.log

echo -e "["$(date)"]\tDONE!"
```

```
#!/bin/bash
#
# AUTHOR: Anand M.
# RNA-Seq variant calling pieline accoring to GATK Best practices.
# https://www.broadinstitute.org/gatk/guide/article?id=3891
#
# Call with following arguments
# bash rna_seq_variant_pipeline.sh <Input_Reads1.fq.gz> <Input_Reads2.fq.gz> <output_basename>
# 
# Assumes STAR aligner is under path
#
# GATK bundle set : one can obtain these from gatk ftp (knonw as gatk bundle)
# ftp://ftp.broadinstitute.org/bundle/2.8/hg19/
#


# sinteractive --account project_2004676 --cores 7 --time 1-00:00:00 --mem 50000 --tmp 100


#!/bin/bash
#SBATCH --job-name=STAR
#SBATCH --output=logs/mapping/STAR.txt
#SBATCH --error=logs/mapping/STAR.txt
#SBATCH --account=project_2004676
#SBATCH --partition=small
#SBATCH --time=2-00:00:00
#SBATCH --ntasks=1
#SBATCH --mem-per-cpu=6000
#SBATCH --cpus-per-task=30

module load biokit
module load gatk

threads=28

# 151 read length
fwd=../data/internal_raw/fastq/Faqu_FKRN220303431-1A_H22GGDSX5_L4_1.fq.gz
rev=../data/internal_raw/fastq/Faqu_FKRN220303431-1A_H22GGDSX5_L4_2.fq.gz
bn=Faqu_FKRN220303431

#Path to reference genome and Index files.
star_ref="../data/external_raw/genome/Faql_flye_noContig_1719_star_index_149"
ref="../data/external_raw/genome/Faql_flye.noContig_1719.fasta"
gtf="intermediate/gemoma/Faql_flye/final_annotation.gff"

opdir=intermediate/RNA_seq_GATK_pipeline/Faql_flye_noContig_1719/$bn


STAR --runThreadN $threads --runMode genomeGenerate --genomeDir $star_ref --genomeFastaFiles $ref --genomeSAindexNbases 13

#picard CreateSequenceDictionary R=$ref O=$(echo $ref | sed 's/.fa//').dict


#Create an output directory
mkdir -p $opdir

#STAR 2 pass basic mode run
echo -e "["$(date)"]\tAligning.."
STAR --outFileNamePrefix $opdir/$bn --outSAMtype BAM Unsorted --outSAMstrandField intronMotif --outSAMattrRGline ID:$bn LB:PairedEnd PL:Illumina PU:Unknown SM:$bn --genomeDir $star_ref --runThreadN $threads --readFilesCommand zcat --readFilesIn $fwd $rev --twopassMode Basic

#sambamba sort
echo -e "["$(date)"]\tSorting.."
sambamba sort -o $opdir/$bn"_sorted.bam" -p -t $threads $opdir/$bn"Aligned.out.bam" --tmpdir=./tmp

#rm $opdir/$bn"Aligned.out.bam"

#sambamba index
echo -e "["$(date)"]\tIndexing.."
sambamba index -p -t $threads $opdir/$bn"_sorted.bam"

#picard mark duplicates
echo -e "["$(date)"]\tMarking duplicates.."
picard MarkDuplicates I=$opdir/$bn"_sorted.bam" O=$opdir/$bn"_dupMarked.bam" M=$opdir/$bn"_dup.metrics" TMP_DIR=./tmp CREATE_INDEX=true VALIDATION_STRINGENCY=SILENT 2>$opdir/$bn.MarkDuplicates.log

rm $opdir/$bn"_sorted.bam"
rm $opdir/$bn"_sorted.bam.bai"

#SplitNCigarReads
echo -e "["$(date)"]\tSpliting reads.."
gatk SplitNCigarReads -R $ref -I $opdir/$bn"_dupMarked.bam" -O $opdir/$bn"_split.bam" --tmp-dir ./tmp 2>$opdir/$bn.SplitNCigarReads.log

rm $opdir/$bn"_dupMarked.bam"
rm $opdir/$bn"_dupMarked.bai"

#Run HaplotypeCaller
echo -e "["$(date)"]\tRunning HaplotypeCaller.."
gatk HaplotypeCaller -R $ref -I $opdir/$bn"_split.bam" -O $opdir/$bn".vcf" 2>$opdir/$bn.HaplotypeCaller.log


#Filter variants
#echo -e "["$(date)"]\tFiltering Variants.."
#gatk VariantFiltration -R $ref -V $opdir/$bn".Scaffold03.vcf" -window 35 -cluster 3 -filterName FS -filter "FS > 30.0" -filterName QD -filter "QD < 2.0" -O $opdir/$bn"_filtered.vcf" 2>$opdir/$bn.VariantFilter.log

echo -e "["$(date)"]\tDONE!"


ref="../data/external_raw/genome/Faql_flye.noContig_1719.fasta"

cat ${ref}.fai | awk '{if($1=="contig_1305" || $1=="contig_189" || $1=="contig_552" || $1=="contig_1603") print $1,"0",$2}' | sed 's/ /\t/g' > Faql_flye.minP.list
samtools view -L Faql_flye.minP.list $opdir/$bn"_split.bam"
samtools view -L Faql_flye.minP.list $opdir/$bn"_split.bam" -O BAM > $opdir/$bn"_split_minP.bam"
samtools index $opdir/$bn"_split_minP.bam"
```

Count features and then use https://github.com/AAlhendi1707/countToFPKM to quantify gene expression per gene

```
sinteractive --account project_2004676 --cores 6 --time 6:00:00 --mem 50000 --tmp 100


#!/bin/bash
#SBATCH --job-name=featureCounts
#SBATCH --account=project_2004676
#SBATCH --partition=small
#SBATCH --time=03:00:00
#SBATCH --ntasks=1
#SBATCH --mem-per-cpu=6000
#SBATCH --cpus-per-task=15

source activate snpsift

featureCounts -a ../data/external_raw/genome/Faqxpol_genome_annotation_v1.gtf intermediate/RNA_seq_GATK_pipeline/Faqxpol_scaffolds_v1/Faqu_FKRN220303431/Faqu_FKRN220303431_split.bam -o intermediate/RNA_seq_GATK_pipeline/Faqxpol_scaffolds_v1/Faqu_FKRN220303431/Faqu_FKRN220303431_split.featureCounts.tsv -p -O -g gene_id -t CDS -T 14


#cat intermediate/gemoma/Faql_flye/final_annotation.gff | grep -v contig_1719 > intermediate/gemoma/Faql_flye/final_annotation.noContig_1719.gff
#ants="/scratch/project_2004676/hanna_sigeman/ants/ants-snakemake2"
#proj="/projappl/project_2004676/hanna_sigeman"

#apptainer exec -B /scratch:/scratch ${proj}/agat_1.0.0--pl5321hdfd78af_0.sif agat_convert_sp_gff2gtf.pl --gff intermediate/gemoma/Faql_flye/final_annotation.noContig_1719.gff -o intermediate/gemoma/Faql_flye/final_annotation.noContig_1719.gtf

featureCounts -a intermediate/gemoma/Faql_flye/final_annotation.noContig_1719.gtf intermediate/RNA_seq_GATK_pipeline/Faql_flye_noContig_1719/Faqu_FKRN220303431/Faqu_FKRN220303431_split.bam -o intermediate/RNA_seq_GATK_pipeline/Faql_flye_noContig_1719/Faqu_FKRN220303431/Faqu_FKRN220303431_split.featureCounts.tsv -p -O -g gene_id -t CDS -T 14


# Input files
featureCounts_output="intermediate/RNA_seq_GATK_pipeline/Faqxpol_scaffolds_v1/Faqu_FKRN220303431/Faqu_FKRN220303431_split.featureCounts.tsv"
transcript_lengths="transcript_lengths.txt"


# Total number of reads
total_reads=$(awk -F'\t' '{sum+=$7;} END{print sum;}' "$featureCounts_output")

# Calculate TPM for each transcript
awk -v total_reads="$total_reads" 'NR > 2 {
    tpm = ($7 * 1e6) / ($6/1000) / total_reads
    print $1, tpm
}' "$featureCounts_output" > intermediate/RNA_seq_GATK_pipeline/Faqxpol_scaffolds_v1/Faqu_FKRN220303431/TPM_output.txt

cat ../data/external_raw/genome/Faqxpol_genome_annotation_v1.gff3 | awk '$3=="gene" {print}' | cut -f 1,4,5,9 | tr -d "ID=;" | while read scaff start end gene ; do cat intermediate/RNA_seq_GATK_pipeline/Faqxpol_scaffolds_v1/Faqu_FKRN220303431/TPM_output.txt | awk '$1=="'"$gene"'" {print "'"$scaff"'","'"$start"'","'"$end"'",$2,$1}'; done | sed 's/ /\t/g' > intermediate/RNA_seq_GATK_pipeline/Faqxpol_scaffolds_v1/Faqu_FKRN220303431/TPM_output_geneInfo.txt
 

featureCounts_output="intermediate/RNA_seq_GATK_pipeline/Faql_flye_noContig_1719/Faqu_FKRN220303431/Faqu_FKRN220303431_split.featureCounts.tsv"
transcript_lengths="transcript_lengths.txt"


# Total number of reads
total_reads=$(awk -F'\t' '{sum+=$7;} END{print sum;}' "$featureCounts_output")

# Calculate TPM for each transcript
awk -v total_reads="$total_reads" 'NR > 2 {
    tpm = ($7 * 1e6) / ($6/1000) / total_reads
    print $1, tpm
}' "$featureCounts_output" > intermediate/RNA_seq_GATK_pipeline/Faql_flye_noContig_1719/Faqu_FKRN220303431/TPM_output.txt

cat intermediate/gemoma/Faql_flye/final_annotation.noContig_1719.gtf | awk '$3=="transcript" {print}' | cut -f 1,4,5,9 | sed 's/ /\t/g' | cut -f 1-3,5,7 | tr -d "\";" | while read scaff start end gene trans ; do cat intermediate/RNA_seq_GATK_pipeline/Faql_flye_noContig_1719/Faqu_FKRN220303431/TPM_output.txt | awk '$1=="'"$gene"'" {print "'"$scaff"'","'"$start"'","'"$end"'",$2,$1,"'"$trans"'"}'; done | sed 's/ /\t/g' > intermediate/RNA_seq_GATK_pipeline/Faql_flye_noContig_1719/Faqu_FKRN220303431/TPM_output_geneInfo.txt

# Faqxpol_JG3505.T1_R2 / nbis-gene-16075 = P-copy 
# Faqxpol_JG3505.T1_R1 / nbis-gene-16074 = M-copy 


picard CollectInsertSizeMetrics I=intermediate/RNA_seq_GATK_pipeline/Faqxpol_scaffolds_v1/Faqu_FKRN220303431/Faqu_FKRN220303431_split.bam O=intermediate/RNA_seq_GATK_pipeline/Faqxpol_scaffolds_v1/Faqu_FKRN220303431/Faqu_FKRN220303431_split_insert_size_metrics.txt M=0.5 H=intermediate/RNA_seq_GATK_pipeline/Faqxpol_scaffolds_v1/Faqu_FKRN220303431/Faqu_FKRN220303431_split_insert_size_metrics.histogram.pdf

echo -e "Faqu_FKRN220303431\t335.895385" > RNA-seq.samples.metrics.txt

start-r


library(countToFPKM)

file.readcounts <- read.delim("test2.featureCounts.tsv", sep = "\t")
file.annotations <- system.file("extdata", "Biomart.annotations.hg38.txt", package="countToFPKM")
file.sample.metrics <- system.file("extdata", "RNA-seq.samples.metrics.txt", package="countToFPKM")

gencov.data <- read.table("results/CDS_coverage/all_samples.Scaffold03.bamstat04.depth", header = T) # Per-exon genome c
gene.start <- aggregate(start ~ CDS, gencov.data, function(x) min(x))
gene.end <- aggregate(end ~ CDS, gencov.data, function(x) max(x))

gene.start.end <- merge(gene.start, gene.end , by = ("CDS"))
gene.start.end <- plyr::rename(gene.start.end, c("start"="Start", "end"="End", "CDS"="Gene_ID"))
gene.start.end$length <- gene.start.end$End - gene.start.end$Start
gene.start.end$Gene_ID <- gsub("\\..*","",gene.start.end$Gene_ID)


folder <- "./"
# reading in featureCounts output
counts <- read.table(paste0(folder, "test2.featureCounts.tsv"), header=TRUE)
counts <- plyr::rename(counts, c("intermediate.RNA_seq_GATK_pipeline.Faqxpol_scaffolds_v1.Faqu_FKRN220303431.Faqu_FKRN220303431_split.bam"="Faql"))

counts$RPK <- counts$Faql/(counts$Length/1000)
sum(counts$Faql) # 28460795
number <- 28460795 / 1000000
counts$TPM <- counts$RPK / number

counts <- counts[order(match(counts[,1],gene.start.end[,1])),]


# Plot log10(FPKM+1) heatmap of top 30 highly variable features
fpkmheatmap(fpkm_matrix, topvar=30, showfeaturenames=TRUE, return_log = TRUE)
```

Plotting

```
library(ggplot2)
library(plyr)
library(dplyr)
library(doBy)
library(reshape2)
library(psych)
library(ggsci)
library(data.table)
library(ggpubr)
library(tidyr)
library(gplots)
library("scales")
library(tibble)
library(ggrepel)

setwd("/scratch/project_2004676/hanna_sigeman/ants/wood-ant-supergene")

text_size_colour = list(theme_bw(base_family="Helvetica", base_size = 12) + 
                          theme(axis.text.x= element_text(colour="black", size=12)) +
                          theme(axis.text.y= element_text(colour="black", size=12)) +
                          theme(axis.title.x = element_text(colour="black",size=14)) + 
                          theme(axis.title.y = element_text(colour="black",size=14)) + 
                          theme(axis.ticks = element_line(colour = "black", size = 0.5)) +
                          theme(panel.border = element_rect(colour = "black", fill=NA)) +
                          theme(plot.title=element_text(family="Courier", size=14, colour="black", hjust = 0.5)) +
                          theme(panel.grid.major = element_line(size = 0.15, linetype = 'solid',
                                                                colour = "lightgrey"), 
                                panel.grid.minor = element_line(size = 0.15, linetype = 'solid',
                                                                colour = "lightgrey")))

theme_title <- function(...) {
  theme_gray(base_family = "Helvetica") + 
    theme(plot.title = element_text(face = "bold"))
}
title_theme <- calc_element("plot.title", theme_title())

theme_description <- function(...) {
  theme_gray(base_family = "Helvetica") + 
    theme(plot.title = element_text(face = "plain",size = 12, hjust = 0))
}
theme_description <- calc_element("plot.title", theme_description())

MyColour <- c("#0073C2FF", "#EFC000FF", "#868686FF", "#CD534CFF", "#7AA6DCFF", "#003C67FF", "#8F7700FF")
names(MyColour) <- c("F_aquilonia", "F_lugubris", "F_paralugubris", "F_polyctena", "F_pratensis", "F_rufa", "F_truncorum")
MyShapes <- c(21, 22, 23, 24)
names(MyShapes) <- c("MM", "MP", "M", "P")

mypal <- pal_jco()(9)
mypal

show_col(mypal)

data <- read.delim("intermediate/RNA_seq_GATK_pipeline/Faqxpol_scaffolds_v1/Faqu_FKRN220303431/TPM_output_geneInfo.txt", sep="\t", header = F)

data <- data %>% rename(Scaffold = V1, start = V2, end = V3, TPM = V4, gene = V5)

subset(data$TPM, data$gene=="jg3505" ) # 40.9143
subset(data$TPM, data$gene=="jg3507" ) # 3.15211

p1 <- ggplot(data, aes(x = TPM)) + xlim(c(0,200)) + 
  geom_histogram(binwidth = 2, colour = "black", fill = "purple") + 
  text_size_colour + geom_vline(xintercept = 40.9143, linetype = "dashed") + 
  geom_vline(xintercept = 3.15211, linetype = "dashed") + 
  annotate("label", label="Zasp52", x=40.9143, y=2000) + 
  annotate("label", label="TTLL2", x=3.15211, y=2500)


p2 <- ggplot(data, aes(x = TPM)) + 
  geom_histogram(binwidth = 2, colour = "black", fill = "purple") + 
  ylab("count") +
  text_size_colour

figure1 <- ggarrange(p1, p2) + ggtitle("hej")

pdf("results/plots/TPM_F.aql_Faqxpol_ref_genome.pdf", width = 12, height = 8)
annotate_figure(figure1, top = text_grob("F. aquilonia RNA data aligned to F. aquilonia x polyctena ref genome ", 
                                         color = "black", face = "bold", size = 14))
dev.off()

data <- read.delim("intermediate/RNA_seq_GATK_pipeline/Faql_flye_noContig_1719/Faqu_FKRN220303431/TPM_output_geneInfo.txt", sep="\t", header = F)
data <- data %>% rename(Scaffold = V1, start = V2, end = V3, TPM = V4, gene = V6)

subset(data$TPM, data$gene~"JG3505") # 40.9143
data[grep("JG3505", data$gene), ]
# 10126  contig_552  2577 7882 19.9085 nbis-gene-16075 Faqxpol_JG3505.T1_R2
# 17326 contig_1305  1889 8579 27.0303 nbis-gene-16074 Faqxpol_JG3505.T1_R1

data[grep("JG3507", data$gene), ]
# 2439   contig_189 11516 17288 1.91913 nbis-gene-16077 Faqxpol_JG3507.T1_R2
# 18658 contig_1603  3054  8722 1.35019 nbis-gene-16076 Faqxpol_JG3507.T1_R0


p1 <- ggplot(data, aes(x = TPM)) + xlim(c(0,200)) + ylim(0,3000) + 
  geom_histogram(binwidth = 2, colour = "black", fill = "purple") + 
  geom_vline(xintercept = 19.9085, linetype = "dashed") + 
  geom_vline(xintercept = 27.0303, linetype = "dashed") + 
  annotate("label", label="Zasp52-P", x=19.9085, y=1500) + 
  annotate("label", label="Zasp52-M", x=27.0303, y=2000) + 
  geom_vline(xintercept = 1.91913, linetype = "dashed") + 
  geom_vline(xintercept = 1.35019, linetype = "dashed") + 
  annotate("label", label="TTLL2-P", x=1.91913, y=2500) + 
  annotate("label", label="TTLL2-M", x=1.35019, y=3000) + 
  ylab("count") +
  text_size_colour

p2 <- ggplot(data, aes(x = TPM)) + 
  geom_histogram(binwidth = 2, colour = "black", fill = "purple") + 
  ylab("count") +
  text_size_colour

figure1 <- ggarrange(p1, p2) 

pdf("results/plots/TPM_F.aql_F.aql_ref_genome.pdf", width = 12, height = 8)
annotate_figure(figure1, top = text_grob("F. aquilonia RNA data aligned to F.aquilonia ref genome ", 
                                         color = "black", face = "bold", size = 14))
dev.off()

data <- subset(data, data$Scaffold=="Scaffold03")

ggplot(data, aes(x = start/1000000, y = TPM)) +
  annotate("rect", xmin = 9.821508,xmax = 9.864738,
           ymin = -Inf,ymax = Inf, alpha = 0.2, fill = "red") +
  scale_shape_manual(values = MyShapes) +
  geom_segment(aes(x = 9.831705, y = 0.6, xend = 9.831705, yend = 0.5), arrow = arrow(length = unit(0.25, "cm"))) +
  geom_segment(aes(x = 9.845014, y = 0.5, xend = 9.845014, yend = 0.4), arrow = arrow(length = unit(0.25, "cm"))) +
  geom_segment(aes(x = 9.856233, y = 0.6, xend = 9.856233, yend = 0.5), arrow = arrow(length = unit(0.25, "cm"))) +
  geom_text_repel(data=filter(data, gene=="jg3505"), aes(label=gene), nudge_y = 3000) +
  # geom_text_repel(data=filter(minimal_P_avg, CDS=="jg3507.t1"), aes(label=CDS), nudge_x = 1, nudge_y = 1) +
  # geom_text_repel(data=filter(minimal_P_avg, CDS=="jg3510.t1"), aes(label=CDS), nudge_x = 1, nudge_y = 1) +
  geom_point() + scale_colour_jama() + scale_fill_jama() + 
  text_size_colour + xlab("Scaffold03 (Mb)") 


minimal_P_exp <- subset(data, c(data$start>9601610 & data$start<10148566))


ggplot(minimal_P_exp, aes(x = start/1000000, y = TPM)) +
  annotate("rect", xmin = 9.821508,xmax = 9.864738,
           ymin = -Inf,ymax = Inf, alpha = 0.2, fill = "red") +
#  geom_segment(aes(x = 9.831705, y = 0.6, xend = 9.831705, yend = 0.5), arrow = arrow(length = unit(0.25, "cm"))) +
#  geom_segment(aes(x = 9.845014, y = 0.5, xend = 9.845014, yend = 0.4), arrow = arrow(length = unit(0.25, "cm"))) +
#  geom_segment(aes(x = 9.856233, y = 0.6, xend = 9.856233, yend = 0.5), arrow = arrow(length = unit(0.25, "cm"))) +
  geom_label_repel(data=filter(minimal_P_exp, gene=="jg3505"), aes(label=gene), nudge_y = 1500) +
  geom_label_repel(data=filter(minimal_P_exp, gene=="jg3507"), aes(label=gene), nudge_y = 1200) +
  geom_label_repel(data=filter(minimal_P_exp, gene=="jg3510"), aes(label=gene), nudge_y = 900) +
  # geom_text_repel(data=filter(minimal_P_avg, CDS=="jg3507.t1"), aes(label=CDS), nudge_x = 1, nudge_y = 1) +
  # geom_text_repel(data=filter(minimal_P_avg, CDS=="jg3510.t1"), aes(label=CDS), nudge_x = 1, nudge_y = 1) +
  geom_point(pch = 21, fill = "purple") +
  text_size_colour + xlab("Scaffold03 (Mb)")
```
