## Supplementary Code for "The loss of a supergene in obligately polygynous *Formica* wood ant species": Code for supergene genotyping 83354545e9a141d4a9e6aa114193b9dd.html

### Code for supergene genotyping

```
# Genotype counts all samples
export LC_ALL=C
join -1 1 -2 1 -t $'\t' <(bcftools query -f '[%SAMPLE\t%GT\n]' intermediate/joint_vcfs/Scaff03_final.woodAnt.hetSites.vcf.gz | sort | uniq -c | awk '{print $2,$3,$1}' | sed 's/ /\t/g' | sort -k1,1) <(sort -k1,1 config/samples_supergene_haplotype_species.tsv) | awk '{if($2=="0/0" || $2=="0/1"  || $2=="1/1") print }' > results/Scaff03_final.woodAnt.hetSites.genotypeCounts.tsv
```

Rcode plotting

```
library(ggplot2)
library(plyr)
library(dplyr)
library(doBy)
library(reshape2)
library(psych)
library(ggsci)
library(data.table)
library(ggpubr)
library(cowplot)
library(vcfR)
library(tidyr)
library("scales")

set.seed(999)
args <- commandArgs(trailingOnly = TRUE)

text_size_colour = list(theme_bw(base_family="Helvetica", base_size = 12) + 
                          theme(axis.text.x= element_text(colour="black", size=12)) +
                          theme(axis.text.y= element_text(colour="black", size=12)) +
                          theme(axis.title.x = element_text(colour="black",size=14)) + 
                          theme(axis.title.y = element_text(colour="black",size=14)) + 
                          theme(axis.ticks = element_line(colour = "black", size = 0.5)) +
                          theme(panel.border = element_rect(colour = "black", fill=NA)) +
                          theme(plot.title=element_text(family="Courier", size=14, colour="black", hjust = 0.5)) +
                          theme(panel.grid.major = element_line(size = 0.15, linetype = 'solid',
                                                                colour = "lightgrey"), 
                                panel.grid.minor = element_line(size = 0.15, linetype = 'solid',
                                                                colour = "lightgrey")))

theme_title <- function(...) {
  theme_gray(base_family = "Helvetica") + 
    theme(plot.title = element_text(face = "bold"))
}
title_theme <- calc_element("plot.title", theme_title())

theme_description <- function(...) {
  theme_gray(base_family = "Helvetica") + 
    theme(plot.title = element_text(face = "plain",size = 12, hjust = 0))
}
theme_description <- calc_element("plot.title", theme_description())


setwd("/scratch/project_2004676/hanna_sigeman/ants/wood-ant-supergene/results")
samples <- read.delim("../config/samples_supergene_haplotype_species.tsv", header = F)
het <- read.table("heterozygosity/Scaff03.deepvariant.biallelic.minQ20.minDP3.het", header = T)
perSampleGenotypes <- read.table("heterozygosity/genotypeCounts.all.samples", sep="\t", header = T)
ok_samples <- read.table("ok_samples.list")
ok_samples <- plyr::rename(ok_samples, c("V1"="sample"))
genes <- read.table("../meta/Faqxpol_genome_annotation_v1.agat.longest.isoform.gff")
vcf <- read.vcfR("../intermediate/joint_vcfs/Scaff03_final.woodAnt.hetSites.CDS.hom10.vcf")
countSNPs <- read.delim("Scaff03_final.woodAnt.hetSites.genotypeCounts.tsv", header = F)
repeats <- read.delim("repeatCumSum.20kb.out", header = F)

samples <- samples %>% rename(sample = V1, Species = V2, supergene = V3)
countSNPs <- countSNPs %>% rename(sample = V1, genotype = V2, count = V3, Species = V4, supergene = V5)
repeats <- repeats %>% rename(chr = V1, start = V2, end = V3, repeat_bp = V4)


vcf_df <- data.frame(
  CHROM = vcf@fix[, "CHROM"],
  POS = vcf@fix[, "POS"],
  ID = vcf@fix[, "ID"],
  REF = vcf@fix[, "REF"],
  ALT = vcf@fix[, "ALT"],
  QUAL = vcf@fix[, "QUAL"],
  INFO = vcf@fix[, "INFO"]
)


mypal <- pal_jco()(9)
mypal
#> [1] "#E64B35B2" "#4DBBD5B2" "#00A087B2" "#3C5488B2" "#F39B7FB2" "#8491B4B2"
#> [7] "#91D1C2B2" "#DC0000B2" "#7E6148B2"

show_col(mypal)

MyColour <- c("#0073C2FF", "#EFC000FF", "#868686FF", "#CD534CFF", "#7AA6DCFF", "#003C67FF", "#8F7700FF")
names(MyColour) <- c("F_aquilonia", "F_lugubris", "F_paralugubris", "F_polyctena", "F_pratensis", "F_rufa", "F_truncorum")

MyShapes <- c(21, 22, 23, 24)
names(MyShapes) <- c("MM", "MP", "M", "P")


genes <- subset(genes, genes$V3=="mRNA")
genes <- subset(genes, genes$V1=="Scaffold03")


het <- plyr::rename(het, c("INDV"="sample"))
het_samples <- merge(het, samples , by = ("sample"))
het_samples <- merge(het_samples, ok_samples , by = ("sample"))
samples_het <- merge(samples, het , by = ("sample"))

perSampleGenotypes2 <- merge(perSampleGenotypes, het , by = ("sample"), all = T)
perSampleGenotypes2 <- merge(perSampleGenotypes2, ok_samples , by = ("sample"))
perSampleGenotypes2 <- merge(perSampleGenotypes2, samples , by = ("sample"))

perSampleGenotypes2[is.na(perSampleGenotypes2)] <- "Unknown"
altHet <- subset(perSampleGenotypes2, perSampleGenotypes2$variable=="altHet")
altHom <- subset(perSampleGenotypes2, perSampleGenotypes2$variable=="altHom")

se_data_altHet <- altHet %>% group_by(Species, start, supergene) %>% summarize(mean_value = mean(value), se = sd(value) / sqrt(n()))
se_data_altHet <- se_data_altHet[order(match(rownames(se_data_altHet), MyColour)), , drop = FALSE]

se_data_altHom <- altHom %>% group_by(Species, start, supergene) %>% summarize(mean_value = mean(value), se = sd(value) / sqrt(n()))
se_data_altHom <- se_data_altHom[order(match(rownames(se_data_altHom), MyColour)), , drop = FALSE]

se_data_altHet_diploids <- subset(se_data_altHet, c(se_data_altHet$supergene=="MP" | se_data_altHet$supergene=="MM"))
se_data_altHom_haploids <- subset(se_data_altHom, c(se_data_altHom$supergene=="M" | se_data_altHom$supergene=="P"))
se_data_haploid_diploids <- rbind(se_data_altHet_diploids, se_data_altHom_haploids)
se_data_haploid_diploids <- se_data_haploid_diploids[order(match(rownames(se_data_haploid_diploids), MyColour)), , drop = FALSE]


# F-score plot

Fscore_plot <- ggplot(het_samples, aes(x = Species, y = F, shape = supergene, fill = Species)) +
  geom_jitter(size = 5) + geom_hline(yintercept = -0.5, linetype = "dashed") + 
  scale_shape_manual(values = MyShapes) +
  scale_fill_manual(values = MyColour) + 
  scale_colour_manual(values = MyColour) + 
  text_size_colour + xlab(" ") + 
  ylab("F") +
  theme(axis.text.x = element_text(angle = 45, vjust = 1, hjust=1))


se_data_haploid_diploids_MP <- subset(se_data_haploid_diploids, se_data_haploid_diploids$supergene=="MP")


supergene_plot <- ggplot(se_data_haploid_diploids_MP, aes(x = start/1000000, y = mean_value, group = Species, fill = Species, color = Species)) +
  annotate("rect", xmin = 1.9, xmax = 11.6, ymin = -Inf, ymax = Inf, alpha = 0.5, fill = "lightgray") +
  geom_line(size = 1) + 
  geom_ribbon(aes(ymin = mean_value - se, ymax = mean_value + se), alpha = 0.9) +
  scale_fill_manual(values = MyColour) + 
  scale_colour_manual(values = MyColour) + 
  text_size_colour +
  xlab(" ") +  ylab(expression(paste("Het. SNPs in MP workers"))) +
  theme(plot.margin = margin(0, 1, 0, 1, "cm")) 

fixed_MP <- ggplot(vcf_df, aes(x = as.numeric(POS)/1000000)) +
  annotate("rect", xmin = 1.9, xmax = 11.6, ymin = -Inf, ymax = Inf, alpha = 0.5, fill = "lightgray") +
  geom_histogram(binwidth = 0.2, colour = "black", fill = "darkgray") +  
  text_size_colour +
  xlab(" ") +  ylab(expression(paste("Fixed MP SNPs"))) +
  theme(plot.margin = margin(0, 1, 0, 1, "cm")) 

combined_supergene_fixed_MP <- ggplot() +
  annotate("rect", xmin = 1.9, xmax = 11.6, ymin = -Inf, ymax = Inf, alpha = 0.5, fill = "lightgray") +
  geom_histogram(data = vcf_df, aes(x = as.numeric(POS)/1000000), binwidth = 0.1, colour = "black", fill = "black") +
  geom_line(data = se_data_haploid_diploids_MP, aes(x = start/1000000, y = mean_value, group = Species, fill = Species, color = Species), size = 1) + 
  geom_ribbon(data = se_data_haploid_diploids_MP, aes(x = start/1000000, y = mean_value, ymin = mean_value - se, ymax = mean_value + se), alpha = 0.9) +
  scale_fill_manual(values = MyColour) + 
  scale_colour_manual(values = MyColour) + 
  text_size_colour +
  xlab("Chromosome 3 (Mb)") +  ylab(expression(paste("Het. SNPs in MP workers"))) +
  theme(plot.margin = margin(0, 1, 0, 1, "cm")) 


countSNPs$proportion <- countSNPs$count / tapply(countSNPs$count, countSNPs$sample, sum)[countSNPs$sample]
# Reshape to wide format
countSNPs_wide <- pivot_wider(countSNPs, 
                              id_cols = c("sample", "Species", "supergene"),
                              names_from = "genotype",
                              values_from = c("proportion", "count"))


countSNPs$supergene_f = factor(countSNPs$supergene, levels=c('MM','MP', 'M', 'P'))
set.seed(123)  # Setting a seed for reproducibility, you can skip this line if you want different results each time

shuffled_df <- countSNPs[sample(nrow(countSNPs)), ]


countSNP_plot <- ggplot(shuffled_df, aes(x = genotype, y = proportion, group = sample, shape = supergene, fill = Species)) +
  geom_line(aes(colour = Species), linetype = "dashed") + 
  geom_jitter(width = 0.1, height = 0.01, size = 5) + 
  text_size_colour + 
  scale_colour_manual(values = MyColour) + 
  scale_fill_manual(values = MyColour) + 
  ylab("Prop. of SNPs") +
  xlab("Genotype") +
  scale_shape_manual(values = MyShapes) + facet_wrap(~supergene_f, nrow = 1)


write.table(countSNPs, "tables/countSNPs.tsv", quote=FALSE, sep="\t", row.names = F, col.names = T, na = "NA")

ggarrange(Fscore_plot + theme(legend.position = 'none'), supergene_plot + theme(legend.position = 'none'), fixed_MP, countSNP_plot + theme(legend.position = 'none'),
          align = "v", axis = "l", heights = c(1, 1, 0.5, 0.7), ncol = 1)

pdf("plots/Figure1_jan2024A_V2.pdf", width = 10, height = 12)
ggarrange(Fscore_plot + theme(legend.position = 'none'), combined_supergene_fixed_MP + theme(legend.position = 'none'), countSNP_plot + theme(legend.position = 'none'),
          align = "v", axis = "l", heights = c(1, 1, 1), ncol = 1)
dev.off()

combined_supergene_fixed_MP


l_supergene_plot <- get_legend(supergene_plot)

l_Fscore_plot <- get_legend(Fscore_plot)

#pdf("plots/Figure1_jan2024A.pdf", width = 10, height = 8)
#ggarrange(Fscore_plot + theme(legend.position = 'none'), supergene_plot + theme(legend.position = 'none'), fixed_MP, 
#          align = "v", heights = c(1, 1, 0.5), ncol = 1)
#dev.off()

ggdraw() + draw_plot(l_supergene_plot, width = 1.9, height = 1.2) +
  draw_plot(l_Fscore_plot, width = 1, height = 0.2)


repeats_plot <- ggplot(repeats, aes(x = start/1000000, y = repeat_bp/200)) +
  annotate("rect", xmin = 1.9,xmax = 11.6, ymin = -Inf,ymax = Inf, alpha = 0.5, fill = "lightgray") +
  geom_area(fill = "darkred") + text_size_colour + xlab(" ") + ylab(expression(paste("% repeats"))) + 
  theme(plot.margin = margin(0, 1, 0, 1, "cm"))
repeats_plot


genes_plot <- ggplot(genes, aes(x = V4/1000000)) +
  annotate("rect", xmin = 1.9,xmax = 11.6, ymin = -Inf,ymax = Inf, alpha = 0.5, fill = "lightgray") +
  geom_histogram(colour = "black", fill = "darkblue", binwidth = 0.1) + text_size_colour + xlab("Scaffold03 (Mb)") + ylab(expression(paste("Nr. of genes"))) + 
  theme(plot.margin = margin(0, 1, 0, 1, "cm"))
genes_plot

pdf("plots/FigureS_repeats_gene_content.pdf", width = 10, height = 8)
ggarrange(repeats_plot + theme(legend.position = 'none'), genes_plot + theme(legend.position = 'none'), 
          align = "hv", heights = c(1, 1), ncol = 1)
dev.off()


se_data_haploid_diploids

pdf("plots/FigureS_altHet.pdf", width = 10, height = 12)
ggplot(se_data_altHet, aes(x = start/1000000, y = mean_value, group = supergene, color = supergene)) +
  annotate("rect", xmin = 1.9, xmax = 11.6, ymin = -Inf,ymax = Inf, alpha = 0.5, fill = "lightgray") +
  geom_line() + geom_ribbon(aes(ymin = mean_value - se, ymax = mean_value + se, fill = supergene), alpha = 0.3) +
  xlab("Chromosome 3 (Mb)") +  ylab(expression(paste("Nr. of heterozygous SNPs (20 kb windows)"))) +
  scale_colour_jama() + scale_fill_jama() + text_size_colour + facet_wrap(~Species, ncol = 1)
dev.off()

pdf("plots/FigureS_altHom.pdf", width = 10, height = 12)
ggplot(se_data_altHom, aes(x = start/1000000, y = mean_value, group = supergene, color = supergene)) +
  annotate("rect", xmin = 1.9, xmax = 11.6, ymin = -Inf,ymax = Inf, alpha = 0.5, fill = "lightgray") +
  geom_line() + geom_ribbon(aes(ymin = mean_value - se, ymax = mean_value + se, fill = supergene), alpha = 0.3) +
  xlab("Chromosome 3 (Mb)") +  ylab(expression(paste("Nr. of alt. homozygous SNPs (20 kb windows)"))) +
  scale_colour_jama() + scale_fill_jama() + text_size_colour + facet_wrap(~Species, ncol = 1)
dev.off()


F_polyctena <- subset(altHet, altHet$Species=="F_polyctena")
se_data <- F_polyctena %>% group_by(supergene, start) %>% summarize(mean_value = mean(value), se = sd(value) / sqrt(n()))
ggplot(se_data, aes(x = start/1000000, y = mean_value, group = supergene, color = supergene)) +
  annotate("rect", xmin = 2.1,xmax = 11.5, ymin = -Inf,ymax = Inf, alpha = 0.5, fill = "lightgray") +
  geom_line() + geom_ribbon(aes(ymin = mean_value - se, ymax = mean_value + se, fill = supergene), alpha = 0.3) +
  scale_colour_jama() + scale_fill_jama() + text_size_colour + xlab("Scaffold03 (Mb)")


F_aquilonia <- subset(altHet, altHet$Species=="F_aquilonia")
se_data <- F_polyctena %>% group_by(supergene, start) %>% summarize(mean_value = mean(value), se = sd(value) / sqrt(n()))
ggplot(se_data, aes(x = start/1000000, y = mean_value, group = supergene, color = supergene)) +
  annotate("rect", xmin = 2.1,xmax = 11.5, ymin = -Inf,ymax = Inf, alpha = 0.5, fill = "lightgray") +
  geom_line() + geom_ribbon(aes(ymin = mean_value - se, ymax = mean_value + se, fill = supergene), alpha = 0.3) +
  scale_colour_jama() + scale_fill_jama() + text_size_colour + xlab("Scaffold03 (Mb)")

F_pratensis <- subset(altHet, altHet$Species=="F_pratensis")
F_pratensis <- subset(F_pratensis, F_pratensis$supergene!="P")
se_data <- F_pratensis %>% group_by(supergene, start) %>% summarize(mean_value = mean(value), se = sd(value) / sqrt(n()))
ggplot(se_data, aes(x = start/1000000, y = mean_value, group = supergene, color = supergene)) +
  annotate("rect", xmin = 2.1,xmax = 11.5, ymin = -Inf,ymax = Inf, alpha = 0.5, fill = "lightgray") +
  geom_line() + geom_ribbon(aes(ymin = mean_value - se, ymax = mean_value + se, fill = supergene), alpha = 0.3) +
  scale_colour_jama() + scale_fill_jama() + text_size_colour + xlab("Scaffold03 (Mb)")

F_polyctena <- subset(altHet, altHet$Species=="F_polyctena")
se_data <- F_polyctena %>% group_by(supergene, start) %>% summarize(mean_value = mean(value), se = sd(value) / sqrt(n()))
ggplot(se_data, aes(x = start/1000000, y = mean_value, group = supergene, color = supergene)) +
  annotate("rect", xmin = 2.1,xmax = 11.5, ymin = -Inf,ymax = Inf, alpha = 0.5, fill = "lightgray") +
  geom_line() + geom_ribbon(aes(ymin = mean_value - se, ymax = mean_value + se, fill = supergene), alpha = 0.3) +
  scale_colour_jama() + scale_fill_jama() + text_size_colour + xlab("Scaffold03 (Mb)")

F_lugubris <- subset(altHet, altHet$Species=="F_lugubris")
se_data <- F_lugubris %>% group_by(supergene, start) %>% summarize(mean_value = mean(value), se = sd(value) / sqrt(n()))
ggplot(se_data, aes(x = start/1000000, y = mean_value, group = supergene, color = supergene)) +
  annotate("rect", xmin = 2.1,xmax = 11.5, ymin = -Inf,ymax = Inf, alpha = 0.5, fill = "lightgray") +
  geom_line() + geom_ribbon(aes(ymin = mean_value - se, ymax = mean_value + se, fill = supergene), alpha = 0.3) +
  scale_colour_jama() + scale_fill_jama() + text_size_colour + xlab("Scaffold03 (Mb)")
```
