## Supplementary Code for "The loss of a supergene in obligately polygynous *Formica* wood ant species": Code to produce de novo genome assemblies and orth 685f4d7d05284cfa9ea5671f8e8dacc5.html

Code to produce de novo genome assemblies and orthology analyses 

### Code to produce de novo genome assemblies and orthology analyses

```
# ============== FAQU ============== # 

#!/bin/bash
#SBATCH --job-name=Faq_flye
#SBATCH --output=Faq_flye.out.log
#SBATCH --error=Faq_flye.err.log
#SBATCH --account=project_2004676
#SBATCH --partition=large
#SBATCH --time=2-00:00:00
#SBATCH --ntasks=1
#SBATCH --mem-per-cpu=6000
#SBATCH --cpus-per-task=40

flye --pacbio-raw scratch/Faq.subreads.fastq.gz --out-dir intermediate/assembly/flye/Faql --threads 8


# ============== FPOL ============== # 

#!/bin/bash
#SBATCH --job-name=Fpol_flye
#SBATCH --output=Fpol_flye.out.log
#SBATCH --error=Fpol_flye.err.log
#SBATCH --account=project_2004676
#SBATCH --partition=large
#SBATCH --time=2-00:00:00
#SBATCH --ntasks=1
#SBATCH --mem-per-cpu=6000
#SBATCH --cpus-per-task=40

flye --pacbio-raw scratch/Fpol.subreads.fastq.gz --out-dir intermediate/assembly/flye/Fpol --threads 35

# ============== Faqxpol ============== # 

#!/bin/bash
#SBATCH --job-name=Faqxpol_flye
#SBATCH --output=Faqxpol_flye.out.log
#SBATCH --error=Faqxpol_flye.err.log
#SBATCH --account=project_2004676
#SBATCH --partition=large
#SBATCH --time=2-00:00:00
#SBATCH --ntasks=1
#SBATCH --mem-per-cpu=6000
#SBATCH --cpus-per-task=40

mkdir -p intermediate/assembly/flye/Faqxpol

flye --pacbio-raw ../data/external_raw/fastq/ERR5383095_Sequel_sequencing_Raw_reads_Faqxpol.fastq.gz --out-dir intermediate/assembly/flye/Faqxpol --threads 35
```

Purge\_dups

```
# Installed the tools according to instructions here: https://github.com/dfguan/purge_dups

echo "/scratch/project_2004676/hanna_sigeman/ants/wood-ant-supergene/scratch/Faq.subreads.fastq.gz" > fofn.list

/projappl/project_2004676/hanna_sigeman/purge_dups/scripts/pd_config.py -n Faql_purgedups.asm intermediate/assembly/flye/Faql/assembly.fasta fofn.list


#!/bin/bash
#SBATCH --job-name=purge_dups
#SBATCH --account=project_2004676
#SBATCH --time=20:00:00
#SBATCH --ntasks=1
#SBATCH --nodes=1
#SBATCH --cpus-per-task=40
#SBATCH --mem-per-cpu=4G
#SBATCH --partition=small

python /projappl/project_2004676/hanna_sigeman/purge_dups/scripts/run_purge_dups.py Faql_purgedups.asm /projappl/project_2004676/hanna_sigeman/purge_dups/src Fpol -p bash
```

Gene annotation liftover

```
proj="/projappl/project_2004676/hanna_sigeman"
gff="intermediate/manual_curation/Faqxpol_genome_annotation_v1.ZaspEdit"
ref="../data/external_raw/genome/Faqxpol_scaffolds_v1.fa"


apptainer exec -B /scratch:/scratch ${proj}/agat_1.0.0--pl5321hdfd78af_0.sif agat_sp_add_start_and_stop.pl -gff ${gff}.gff3 --fasta $ref --out ${gff}.MP.start.stop.gff3


grep Scaffold03 ${gff}.MP.start.stop.gff3 | awk '$3=="mRNA" {print}' |  cut -f 1,9 | sed 's/ID=//' | sed 's/;/\t/g' | cut -f 2 > ${gff}.Scaffold03.trans.list


# ====================== # 

#!/bin/bash
#SBATCH --job-name=gemoma_Fpol
#SBATCH --account=project_2004676
#SBATCH --partition=small
#SBATCH --time=08:00:00
#SBATCH --ntasks=1
#SBATCH --mem-per-cpu=6000
#SBATCH --cpus-per-task=40

module load biokit

conda activate gemoma

target=intermediate/assembly/flye/Fpol/assembly.fasta
ref=../data/external_raw/genome/Faqxpol_scaffolds_v1.fa
gff=intermediate/manual_curation/Faqxpol_genome_annotation_v1.ZaspEdit.start.stop.gff3
out=intermediate/gemoma/Fpol_flye_ZaspEdit

prog=/projappl/project_2004676/hanna_sigeman/envs/gemoma/share/gemoma-1.6.4-1/GeMoMa-1.6.4.jar

java -jar $prog CLI GeMoMaPipeline threads=38 outdir=$out GeMoMa.Score=ReAlign AnnotationFinalizer.r=NO o=true t=$target i=Faqxpol a=$gff g=$ref

# ====================== # 

#!/bin/bash
#SBATCH --job-name=gemoma_Faql
#SBATCH --account=project_2004676
#SBATCH --partition=small
#SBATCH --time=08:00:00
#SBATCH --ntasks=1
#SBATCH --mem-per-cpu=6000
#SBATCH --cpus-per-task=40

module load biokit

conda activate gemoma

target=intermediate/assembly/flye/Faql/assembly.fasta
ref=../data/external_raw/genome/Faqxpol_scaffolds_v1.fa
gff=intermediate/manual_curation/Faqxpol_genome_annotation_v1.ZaspEdit.start.stop.gff3
out=intermediate/gemoma/Faql_flye_ZaspEdit

prog=/projappl/project_2004676/hanna_sigeman/envs/gemoma/share/gemoma-1.6.4-1/GeMoMa-1.6.4.jar

java -jar $prog CLI GeMoMaPipeline threads=38 outdir=$out GeMoMa.Score=ReAlign AnnotationFinalizer.r=NO o=true t=$target i=Faqxpol a=$gff g=$ref

# ====================== # 

#!/bin/bash
#SBATCH --job-name=gemoma_Faqxpol
#SBATCH --account=project_2004676
#SBATCH --partition=small
#SBATCH --time=08:00:00
#SBATCH --ntasks=1
#SBATCH --mem-per-cpu=6000
#SBATCH --cpus-per-task=40

module load biokit

source activate gemoma

target=intermediate/assembly/flye/Faqxpol/assembly.fasta
ref=../data/external_raw/genome/Faqxpol_scaffolds_v1.fa
gff=intermediate/manual_curation/Faqxpol_genome_annotation_v1.ZaspEdit.start.stop.gff3
out=intermediate/gemoma/Faqxpol_flye_ZaspEdit

prog=/projappl/project_2004676/hanna_sigeman/envs/gemoma/share/gemoma-1.6.4-1/GeMoMa-1.6.4.jar

java -jar $prog CLI GeMoMaPipeline threads=38 outdir=$out GeMoMa.Score=ReAlign AnnotationFinalizer.r=NO o=true t=$target i=Faqxpol a=$gff g=$ref
```

```
# Installed the tools according to instructions here: https://github.com/dfguan/purge_dups

echo "/scratch/project_2004676/hanna_sigeman/ants/wood-ant-supergene/scratch/Faq.subreads.fastq.gz" > fofn.list

/projappl/project_2004676/hanna_sigeman/purge_dups/scripts/pd_config.py -n Faql_purgedups.asm intermediate/assembly/flye/Faql/assembly.fasta fofn.list


#!/bin/bash
#SBATCH --job-name=purge_dups
#SBATCH --account=project_2004676
#SBATCH --time=20:00:00
#SBATCH --ntasks=1
#SBATCH --nodes=1
#SBATCH --cpus-per-task=40
#SBATCH --mem-per-cpu=4G
#SBATCH --partition=small

python /projappl/project_2004676/hanna_sigeman/purge_dups/scripts/run_purge_dups.py Faql_purgedups.asm /projappl/project_2004676/hanna_sigeman/purge_dups/src Fpol -p bash
```

MinP liftover

```
intermediate/manual_curation/Faqxpol_genome_annotation_v1.minP.gff3

intermediate/manual_curation/Faqxpol_genome_annotation_v1.minP.start.stop.gff3

sinteractive --account project_2004676 --cores 2 --mem 5000 --tmp 100

module load biokit

conda activate gemoma
prog=/projappl/project_2004676/hanna_sigeman/envs/gemoma/share/gemoma-1.6.4-1/GeMoMa-1.6.4.jar

target=intermediate/assembly/flye/Fpol/assembly.fasta
ref=../data/external_raw/genome/Faqxpol_scaffolds_v1.fa
gff=intermediate/manual_curation/Faqxpol_genome_annotation_v1.minP.start.stop.gff3
out=intermediate/gemoma/Fpol_minP

java -jar $prog CLI GeMoMaPipeline threads=2 outdir=$out GeMoMa.Score=ReAlign AnnotationFinalizer.r=NO o=true t=$target i=Faqxpol a=$gff g=$ref GAF.a="pAA>=0.6" GeMoMa.sil=false


target=intermediate/assembly/flye/Faql/assembly.fasta
ref=../data/external_raw/genome/Faqxpol_scaffolds_v1.fa
gff=intermediate/manual_curation/Faqxpol_genome_annotation_v1.minP.start.stop.gff3
out=intermediate/gemoma/Faql_minP

java -jar $prog CLI GeMoMaPipeline threads=2 outdir=$out GeMoMa.Score=ReAlign AnnotationFinalizer.r=NO o=true t=$target i=Faqxpol a=$gff g=$ref GAF.a="pAA>=0.6" GeMoMa.sil=false

target=intermediate/assembly/flye/Faqxpol/assembly.fasta
ref=../data/external_raw/genome/Faqxpol_scaffolds_v1.fa
gff=intermediate/manual_curation/Faqxpol_genome_annotation_v1.minP.start.stop.gff3
out=intermediate/gemoma/Faqxpol_minP

java -jar $prog CLI GeMoMaPipeline threads=2 outdir=$out GeMoMa.Score=ReAlign AnnotationFinalizer.r=NO o=true t=$target i=Faqxpol a=$gff g=$ref GAF.a="pAA>=0.6" GeMoMa.sil=false
```

Make amino acid gene tree for Zasp52 and TTLL2 (Figure 4)

```
# Get all protein sequences into one file
for sp in Faql Fpol Faqxpol; do cat intermediate/gemoma/${sp}_minP/final_annotation.gff | cut -f 1,9 | grep score | grep -v SOFTWARE | sed 's/ID=//' | sed 's/;/\t/' | cut -f 1,2 | sed 's/Faqxpol/FAQXPOL/' | while read contig trans ; do samtools faidx intermediate/gemoma/${sp}_minP/predicted_proteins.fasta $trans | sed "s/>/>${sp}_${contig}_/" ; done ; done > results/gene_seq/minP_predicted_proteins.fasta

# Zasp52
for sp in Faql Fpol Faqxpol; do cat intermediate/gemoma/${sp}_minP/final_annotation.gff | cut -f 1,9 | grep score | grep ZASP | grep -v SOFTWARE | sed 's/ID=//' | sed 's/;/\t/' | cut -f 1,2 | sed 's/Faqxpol/FAQXPOL/' | while read contig trans ; do samtools faidx intermediate/gemoma/${sp}_minP/predicted_proteins.fasta $trans | sed "s/>/>${sp}_${contig}_/" ; done ; done > results/gene_seq/minP_predicted_proteins_ZASP52.fasta

### Downloaded orthoDB genes from here: https://data.orthodb.org/v11/fasta?id=126710at7399&species= 
### To file: results/gene_seq/ZASP52_orthoDB.fasta

cat results/gene_seq/ZASP52_orthoDB.fasta | sed 's/organism_name":"/\t/' | sed 's/>[^=]*\t/>/' | sed 's/"/\t/' | cut -f 1 | sed 's/ /_/g' | sed '/^$/d' > results/gene_seq/ZASP52_orthoDB.clean.fasta

# TTLL2
for sp in Faql Fpol Faqxpol; do cat intermediate/gemoma/${sp}_minP/final_annotation.gff | cut -f 1,9 | grep score | grep JG3507 | grep -v SOFTWARE | sed 's/ID=//' | sed 's/;/\t/' | cut -f 1,2 | sed 's/Faqxpol/FAQXPOL/' | while read contig trans ; do samtools faidx intermediate/gemoma/${sp}_minP/predicted_proteins.fasta $trans | sed "s/>/>${sp}_${contig}_/" ; done ; done > results/gene_seq/minP_predicted_proteins_TTLL2.fasta

### Downloaded orthoDB genes from here: https://data.orthodb.org/v11/fasta?id=97990at7399&species=
### To file: results/gene_seq/TTLL2_orthoDB.fasta

cat results/gene_seq/TTLL2_orthoDB.fasta | sed 's/organism_name":"/\t/' | sed 's/>[^=]*\t/>/' | sed 's/"/\t/' | cut -f 1 | sed 's/ /_/g' | sed '/^$/d' > results/gene_seq/TTLL2_orthoDB.clean.fasta

cat results/gene_seq/minP_predicted_proteins_ZASP52.fasta results/gene_seq/ZASP52_orthoDB.clean.fasta | tr -d "*" > results/gene_seq/allSpecies_ZASP52.fasta

cat results/gene_seq/minP_predicted_proteins_TTLL2.fasta results/gene_seq/TTLL2_orthoDB.clean.fasta | tr -d "*" > results/gene_seq/allSpecies_TTLL2.fasta

clustalo -i results/gene_seq/ZASP52_orthoDB.clean.fasta -t Protein --threads 5 > results/gene_seq/ZASP52_orthoDB.clean.aln

clustalo -i results/gene_seq/TTLL2_orthoDB.clean.fasta -t Protein --threads 5 > results/gene_seq/TTLL2_orthoDB.clean.aln

clustalo -i results/gene_seq/allSpecies_ZASP52.fasta -t Protein --threads 5 > results/gene_seq/allSpecies_ZASP52.aln

clustalo -i results/gene_seq/allSpecies_TTLL2.fasta -t Protein --threads 5 > results/gene_seq/allSpecies_TTLL2.aln


echo "Pseudomyrmex_gracilis
Cardiocondyla_obscurior
Harpegnathos_saltator
Odontomachus_brunneus
Dinoponera_quadriceps
Ooceraea_biroi
Linepithema_humile
Camponotus_floridanus
Formica_exsecta
Nylanderia_fulva
Lasius_niger
Pogonomyrmex_barbatus
Temnothorax_curvispinosus
Monomorium_pharaonis
Vollenhovia_emeryi
Solenopsis_invicta
Wasmannia_auropunctata
Cyphomyrmex_costatus
Acromyrmex_echinatior
Trachymyrmex_zeteki
Trachymyrmex_cornetzi
Trachymyrmex_septentrionalis
Atta_colombica" > results/gene_seq/ant_species.list

seqkit grep -n -f results/gene_seq/ant_species.list results/gene_seq/ZASP52_orthoDB.clean.fasta | seqkit seq -m 600 -g > results/gene_seq/ZASP52_orthoDB.clean.ants.noShort.fasta

seqkit grep -n -f results/gene_seq/ant_species.list results/gene_seq/TTLL2_orthoDB.clean.fasta | seqkit seq -m 400 -g > results/gene_seq/TTLL2_orthoDB.clean.ants.noShort.fasta

# Combine files (but only do the short Zasp52 gene seq and delete contig_1719)
seqkit grep -p 'ZASP52_DUP' results/gene_seq/minP_predicted_proteins_ZASP52.fasta -r | seqkit grep -p "contig_1719" -r -v | cat - results/gene_seq/ZASP52_orthoDB.clean.ants.noShort.fasta | tr -d "*" > results/gene_seq/allAnts_ZASP52.fasta.tmp

# Get the reference sequence as well 
ref=../data/external_raw/genome/Faqxpol_scaffolds_v1.fa
gff=intermediate/manual_curation/Faqxpol_genome_annotation_v1.minP.start.stop.gff3
proj=/projappl/project_2004676/hanna_sigeman

apptainer exec -B /scratch:/scratch ${proj}/agat_1.0.0--pl5321hdfd78af_0.sif agat_sp_extract_sequences.pl -gff $gff -f $ref -p -o results/gene_seq/minP_Faqxpol_scaffolds.fasta

seqkit grep -p 'Zasp52_DUP' results/gene_seq/minP_Faqxpol_scaffolds.fasta | cat - results/gene_seq/allAnts_ZASP52.fasta.tmp > results/gene_seq/allAnts_ZASP52.fasta
rm results/gene_seq/allAnts_ZASP52.fasta.tmp


seqkit grep -p "contig_1719" -r -v results/gene_seq/minP_predicted_proteins_TTLL2.fasta | cat - results/gene_seq/TTLL2_orthoDB.clean.ants.noShort.fasta | tr -d "*" > results/gene_seq/allAnts_TTLL2.fasta.tmp

seqkit grep -p 'jg3507' -r results/gene_seq/minP_Faqxpol_scaffolds.fasta | cat - results/gene_seq/allAnts_TTLL2.fasta.tmp > results/gene_seq/allAnts_TTLL2.fasta
rm results/gene_seq/allAnts_TTLL2.fasta.tmp

clustalo -i results/gene_seq/allAnts_ZASP52.fasta -t Protein --threads 5 > results/gene_seq/allAnts_ZASP52.aln

clustalo -i results/gene_seq/allAnts_TTLL2.fasta -t Protein --threads 5 > results/gene_seq/allAnts_TTLL2.aln

trimal -in results/gene_seq/allAnts_ZASP52.aln -out results/gene_seq/allAnts_ZASP52.trim.aln -strictplus

trimal -in results/gene_seq/allAnts_TTLL2.aln -out results/gene_seq/allAnts_TTLL2.trim.aln -strictplus

trimal -in results/gene_seq/allAnts_ZASP52.trim.aln -out results/gene_seq/allAnts_ZASP52.trim2.aln -gt 0.8

trimal -in results/gene_seq/allAnts_TTLL2.trim.aln -out results/gene_seq/allAnts_TTLL2.trim2.aln -gt 0.8

iqtree2 -s results/gene_seq/allAnts_ZASP52.trim2.aln -m LG+F+G -nt AUTO

iqtree2 -s results/gene_seq/allAnts_TTLL2.trim2.aln -m LG+F+G -nt AUTO

python code/scripts/seq_similarity.py results/gene_seq/allAnts_ZASP52.trim2.aln > results/gene_seq/allAnts_ZASP52.trim2.seq_similarity.txt

python code/scripts/seq_similarity.py results/gene_seq/allAnts_TTLL2.trim2.aln > results/gene_seq/allAnts_TTLL2.trim2.seq_similarity.txt


cat results/gene_seq/allSpecies_TTLL2.sequences.tsv | cut -f 6 | grep -v pub > results/gene_seq/allSpecies_TTLL2.sequences.list

while read -r gene_id; do
  esearch -db nuccore -query "$gene_id" < /dev/null | efetch -format fasta 
done < results/gene_seq/allSpecies_TTLL2.sequences.list


cat results/gene_seq/allSpecies_TTLL2.sequences.tsv | cut -f 6 | grep -v pub | while read -r gene ; do esearch -db nuccore -query "${gene}" | efetch -format fasta | seqkit grep -p "transcript" -r -n ; done


cat results/gene_seq/allSpecies_TTLL2.sequences.tsv | cut -f 6 | grep -v pub | while read gene ; do esearch -db nuccore -query "${gene}" < /dev/null | efetch -format fasta | seqkit grep -p "transcript" -r -n | awk '/^>/{if(length(seq)>max_length){max_length=length(seq);max_seq_id=seq_id;} seq_id=$0; seq=""; next} {seq=seq$0} END{if(length(seq)>max_length){print max_seq_id; print seq;}else{print seq_id; print seq;}}' ; done > results/gene_seq/allSpecies_TTLL2.nucl.fasta

cat results/gene_seq/allSpecies_ZASP52.sequences.tsv | cut -f 6 | grep -v pub | while read gene ; do esearch -db nuccore -query "${gene}" < /dev/null | efetch -format fasta | seqkit grep -p "transcript" -r -n | awk '/^>/{if(length(seq)>max_length){max_length=length(seq);max_seq_id=seq_id;} seq_id=$0; seq=""; next} {seq=seq$0} END{if(length(seq)>max_length){print max_seq_id; print seq;}else{print seq_id; print seq;}}' ; done > results/gene_seq/allSpecies_ZASP52.nucl.fasta
```
